# Unraveling activation-related rearrangements and intrinsic divergence from ligand-induced conformational changes of the dopamine D3 and D2 receptors

**DOI:** 10.1101/2023.11.11.566699

**Authors:** Kuo Hao Lee, Lei Shi

## Abstract

Effective rational drug discovery targeting a specific protein hinges on understanding their functional states and distinguishing it from homologs. However, for the G protein coupled receptors, both the activation-related conformational changes (ACCs) and the intrinsic divergence among receptors can be misled or obscured by ligand-induced conformational changes (LCCs). Here, we unraveled ACCs and intrinsic divergence from LCCs of the dopamine D3 and D2 receptors (D3R and D2R), by analyzing their experimentally determined structures and the molecular dynamics simulation results of the receptors bound with different ligands. In addition to the ACCs common to other aminergic receptors, we revealed unique ACCs for these two receptors including TM5e and TM6e shifting away from TM2e and TM3e, with a subtle rotation of TM5e. In identifying intrinsic divergence, we found pronounced outward tilting of TM6e in the D2R compared to the D3R in both experimental structures and simulations with ligands in different scaffolds. This tilting was drastically reduced in the simulations of the receptors bound with nonselective full agonist quinpirole, suggesting a misleading impact of LCCs. Further, in the quinpirole-bound simulations, TM1 showed a greater disparity between these receptors, indicating that LCCs may obscure intrinsic divergence. In addition, our analysis showed that the impact of the nonconserved TM1 propagated to conserved Trp^7.40^ and Glu^2.65^, both are ligand binding residues. We also found that the D2R exhibited heightened flexibility compared to the D3R in the extracellular portions of TMs 5, 6, and 7, potentially associated with its greater ligand binding site plasticity. Our results lay the groundwork for crafting ligands specifically targeting D2R or D3R with more precise pharmacological profiles.

## Introduction

The dopamine D2 and D3 receptors (D2R and D3R) are G-protein coupled receptors (GPCRs), belonging to the D2-like subgroup of the dopamine receptor subfamily in the class A rhodopsin-like GPCRs.^1^ Along the G protein signaling pathway, both the D2R and D3R activate the Gi/o proteins to inhibit adenylate cyclase. The ligands of these two receptors have been used or are being developed for the treatments of various neuropsychiatric disorders. Specifically, the D2R antagonists are used for schizophrenia,^2^ the D3R antagonists or partial agonists are being actively developed for substance use disorders,^3, 4^ while agonists targeting these receptors have demonstrated promising results in mitigating the symptoms of Parkinson’s disease.^5-7^ The different distributions of the D2R and D3R in the brain regions^8^ and their functional distinctions^9^ prompt the development of subtype-selective ligands with potentially fewer side effects.^3, 10^ However, it has been challenging to rationally develop subtype-selective D2R or D3R ligands with controlled efficacy and specificity for signaling pathways, due to the high sequence homology between these two receptors as well as the difficulty in identifying activation-related conformational changes (ACCs) in their ligand binding pockets.^4, 11-14^

The X-ray crystal structure of the D3R in an inactive conformation became available in 2010,^11^ and was followed by the crystal structures of the D2R in somewhat different inactive conformations.^15-18^ Recently, the cryogenic electron microscopy (cryo-EM) structures of the active conformations of D2R and D3R in complex with the Gi proteins have been solved, providing critical insights with regards to the activation mechanisms of these receptors.^19-21^ However, distinguishing the ACCs from the conformational changes necessary to accommodate specific ligands, i.e., ligand-specific conformational changes (LCCs), poses a significant challenge. This complexity arises because the ligands bound to the active conformations of the D3R and D2R, pramipexole, PD128907, and bromocriptine are of noticeably different scaffolds compared to each other and to the antagonists or inverse agonists bound in the inactive structures (eticlopride, risperidone, haloperidol, and spiperone). As a result, the conformational differences observed between the active and inactive structures may be attributed to either ACCs or LCCs, with potential overlaps between the two.

To effectively utilize these structures to address the challenges of selectively targeting D2R versus D3R, it is crucial to identify the conformational differences that reflect the intrinsic divergence between these two receptors.^13^ Although these divergences naturally stem from the regions with different primary sequences, they can extend and propagate to other parts of the receptors consisting of conserved residues. Therefore, the divergence may also exist and have functional consequence in seemingly conserved regions, such as the ligand binding pockets and the receptor-signaling protein interface. Detection of these divergences in experimentally determined structures can be obscured by either LCCs, ACCs, or both. For example, whereas pramipexole and PD128907 bound the D3R active structures have been reported to possess the D3R over the D2R selectivity,^20, 22, 23^ the extent to which the differences between the D3R and D2R active structures are the intrinsic divergence between these two highly homologous receptors, is not clear. This is because, in addition to the intrinsic divergences, the observed differences may include the elements of LCCs. Furthermore, the efficacy of bromocriptine at the D2R has not been consistently reported, ranging from partial to full agonism.^19, 23, 24^ Therefore, it is possible that the D2R structures bound with bromocriptine may not be as active as the D3R active structures, and their disparities may also include some ACCs.

Recently, we have developed an analysis infrastructure that allows for a superposition-independent comparisons of GPCR structures. Using this infrastructure, we have analyzed the available structures of aminergic receptors to identify common ACCs that are independent of LCCs ^14^. While our analysis reveals no shared ACCs at the extracellular end of the transmembrane (TM) domain, we found common ACCs associated with the Pro^5.50^-Ile^3.40^-Phe^6.44^ (PIF) motif (the superscripts denote Ballesteros-Weinstein numbering^25^) and nearby residues, including the previously proposed toggle switch Trp^6.48^, at the bottom of the ligand binding pocket. We proposed a novel “activation switch” motif integrating the PIF motif and Trp^6.48^, shared by all aminergic receptors.^14^

In this study, we first comprehensively compared the inactive and active conformations revealed by the experimentally determined D3R and D2R structures. In addition to the ACCs shared among aminergic receptors^14^, we aimed to identify ACCs that may be specific to these two receptors. We then adapted the analysis infrastructure to identify conformational differences that could reflect intrinsic divergence between the D3R and D2R. To gain further insights in a more physiologically relevant environment, we built the D2R and D3R active models bound with the ligands bound in the cryo-EM structures, immersed them in the lipid-bilayer environment, and conducted molecular dynamics (MD) simulations at 310 K. Further, to thoroughly investigate the intrinsic divergence between the D2R and D3R, we established and equilibrated the active-state models of both receptors bound with quinpirole, a high-affinity and nonselective full agonist that has been frequently utilized in the D2R and D3R research.^22, 26^ Together, our comprehensive analysis of the conformational differences observed in both the experimentally determined structures and computational simulations shed lights on the activation mechanism, the ligand binding pocket plasticity, and the intrinsic divergence between these two receptors.

## Results

### Common activation conformational changes of the D2R and D3R

In our recent analysis of the experimentally determined structures of aminergic receptors, we developed several superposition-independent and distance-based quantitative metrics to detect commonly occurring conformational changes during receptor activation.^14^ These metrics were designed to evaluate the rearrangements among the extracellular ends, ligand binding residues, or subsegments of the TM domain (see their definitions in Methods). To ensure accuracy, these analyses require the receptor being analyzed to have high-resolution structures available for both the inactive and active states, which is the situation for both the D2R and D3R (**Table S1**). Thus, in the current study, utilizing these metrics and analysis protocols, we first carried out an in-depth investigation to determine whether there are additional ACCs shared between these two receptors in addition to those commonly found for the aminergic family ^14^.

In these analyses, we first computed pairwise distances among specific structural elements for each structure under investigation. For each pair of active and inactive structures of each receptor, such as the active D2R structure 7JVR and the inactive D2R structure 6CM4 (referred to as “7JVR-6CM4” below), we calculated the differences in corresponding distances. These differences were then classified into three categories: 1, -1, and 0, based on a threshold (referred to as distance-difference threshold, DDthreshold), and we integrated the results of all structure pairs by averaging the derived categories (see Methods for details). For instance, in the extracellular end analysis of the D2R, we measured pairwise distances among the centers of mass of the extracellular ends of each D2R structure. For each pair of active and inactive D2R structures (7JVR-6CM4, 7JVR-6LUQ, and 7JVR-7DFP), we calculated the differences in corresponding distances between the two structures. These differences were categorized and then averaged to identify the common changes of the extracellular ends in the D2R (**Fig. S1**). Using the same protocol, we also categorized and averaged the changes of ligand binding residues and subsegments (**Figs. S2-4**). Similarly, the same analyses for the D3R structures determined the trends specific to the D3R (**Figs. S5-8**). By averaging the results of the D2R and D3R analyses, we identified the common changes observed in both receptors (**Figs. S9-12**).

The results of our extracellular end analysis showed that the extracellular loop 2 (EL2) and the extracellular end of TM7 (TM7ee, see definition of extracellular ends in Methods) move closer to each other in the activation of both the D2R and D3R (**Fig. S9**). This suggests a restricted access to the ligand binding pocket in the active state, which has been documented for the β2 adrenergic receptor and a few other GPCRs,^27^ and detected by us for many subfamilies of the aminergic receptors.^14^ In addition to this relatively common change, we observed a common trend in both receptors where TM2ee moves away from TM5ee, TM6ee, and TM7ee by combining the analysis results of both the sidechains and backbones (**Fig. S9**).

In the ligand binding residue analysis (**Figs. S2, S6, and S10**), in addition to the common ACC identified at the aminergic family level that are associated with the activation switch (formed by residues at positions 3.36-3.40, 5.46-5.50, and 6.44-6.48),^14^ we found other changes associated with the switch common to both the D2R and D3R. Specifically, the sidechain of Ser^5.42^ gets further away from Cys^3.36^, while that of Ser^5.43^ gets closer to Thr^3.37^ and Ile^3.40^ (**Fig. 1B**). This trend is also observed for the backbone when DDthreshold is reduced to 0.2 Å (**Fig. S10**), indicating a small rotation of TM5e in receptor activation, shifting Ser^5.43^ towards the ligand binding pocket while moving Ser^5.42^ away. Note that Ser^5.42^ and Ser^5.43^ form hydrogen bonds (H-bonds) with the agonists in the D3R active structures (PDB: 7CMU and 7CMV) but not with the bromocriptine bound in the D2R active structure (PDB: 7JVR) (**Fig. 1B**). Thus, this rotation of TM5e is likely intrinsically associated for the activation of these two receptors – while some specific receptor-agonist interactions may facilitate this rotation, such interactions may not be required.

**Figure 1.**
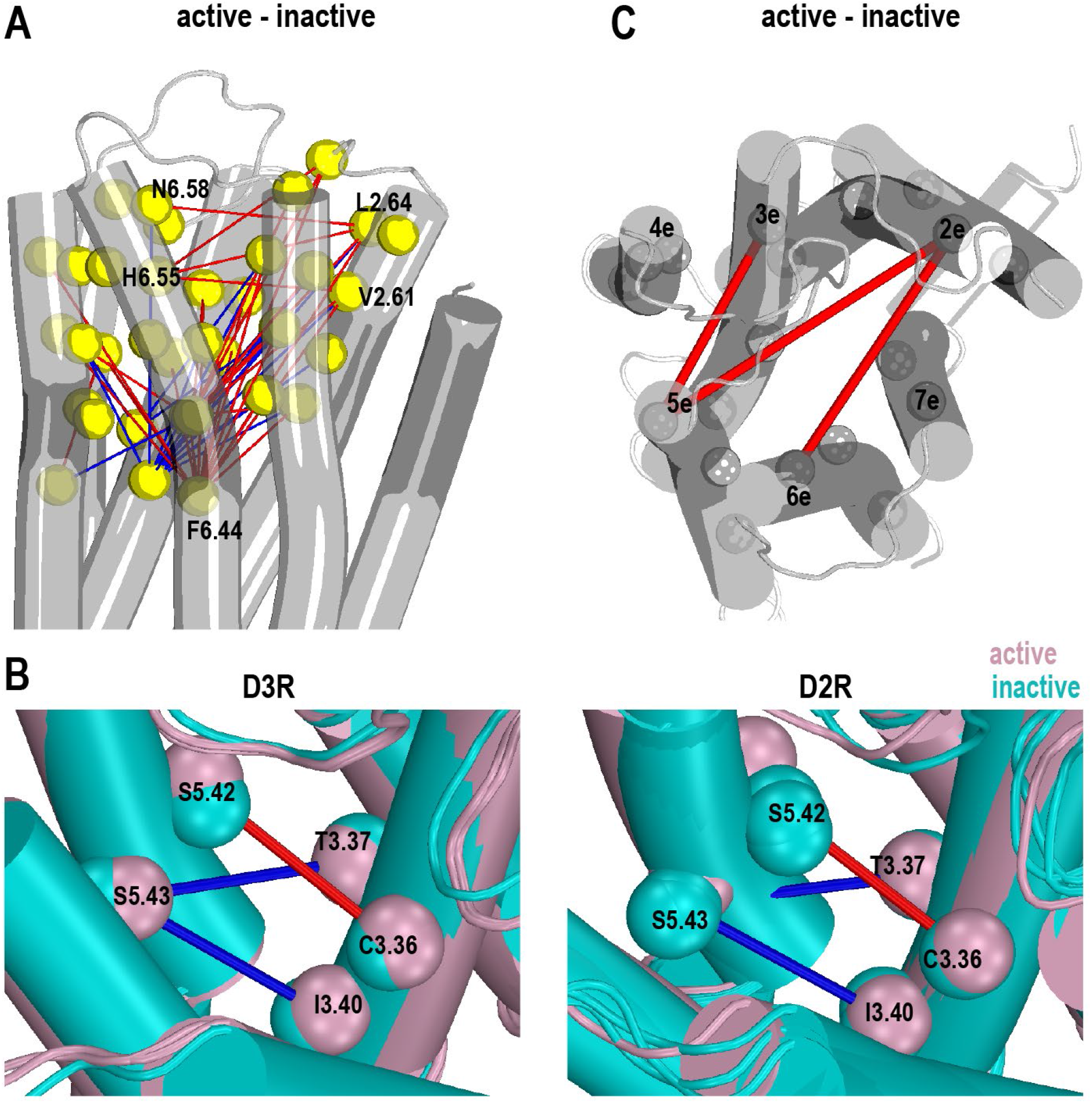
Common activation conformational changes in the D2R and D3R structures. Panel A maps the categorized pairwise distance differences among binding site residues according to the results using a DDthreshold of 1.0 Å (see **Fig. S10**) on the D3R inactive structure (PDB 7CMU). Yellow spheres are the Cα atoms of binding site residues. The red and blue lines indicate that the pairwise distances larger than 1.0 Å in the active and inactive structures, respectively. Panel B compares the common changes of the serine residues in TM5e in both D2R and D3R structures. The spheres represent the centers of mass (COMs) of the indicated sidechain residues. Ser^5.43^ is positioned closer to Thr^3.37^ and Ile^3.40^ in the inactive structure, while Ser^5.42^ and Cys^3.36^ have greater distances in the active structures. Panel C maps the categorized pairwise distance changes among subsegments (see method for subsegments definition) using DDthreshold of 0.6 Å (see **Fig. S12**). The COMs of the subsegments are shown by the gray spheres. The red and blue lines indicate greater and smaller distances in the active and inactive conformations, respectively.

We previously defined the residues commonly interacting with all orthosteric ligands as the orthosteric binding site (OBS) residues, which form the core of the ligand binding pocket.^1^ Among the OBS residues, in addition to the prominent longer distance between Trp^6.48^ and Asp^3.32^ observed at the aminergic family level,^14^ the sidechain of Trp^6.48^ also moves away from those of Thr^7.39^ and Tyr^7.43^ during the activation (**Figs. S3, S7, and S11**). Note that Tyr^7.43^ forms a conserved H-bond with Asp^3.32^, while Thr^7.39^ is one helical turn above Tyr^7.43^.

Among the ligand binding residues not related to the activation switch, by combining the analysis results of both the sidechains and backbones, we found that His^6.55^ and Asn^6.58^ move away from Val^2.61^ and/or Leu^2.64^ in both the D2R and D3R (**Fig. 1A**). These changes are consistent with the trend of TM2ee moving away from TM6ee in the extracellular end analysis (**Fig. S9**), as well as the longer distance of TM2e-TM6e in the subsegment analysis (see below).

In the subsegment analysis (**Figs. S4, S8, and S12**), similar to the patterns observed in other GPCRs (reviewed in ref ^14, 28^), the most prominent intracellular changes of these two receptors in activation are an outward swinging of the intracellular portion of TM6 (TM6i) towards the lipids and an inward movement of TM7i towards the core of the TM domain. These changes are critical in opening an intracellular space of the receptors to accommodate the binding of signaling proteins. Among the middle subsegments forming the activation switch, TM3m gets subtly distant from TMs 5m and 6m during the activation (**Fig. S12**). In addition to these ACCs shared with other GPCRs, among the extracellular subsegments that contain most of the ligand binding residues, TM5e moves away from TM2e and TM3e, while TM6e also moves away from TM2e (**Figs. 1C, S12**). As no common movement among the extracellular subsegments was detected at the aminergic family level, such rearrangements among TMs 2e, 3e, 5e, and 6e can potentially be the ACCs specific for the D2R and D3R.

### Common differences between the D2R and D3R structures in both inactive and active states

We then employed the same metrics and analysis protocol to investigate the divergence between the D2R and D3R, by comparing the differences among the pairs of their inactive structures and among the pairs of their active structures. Specifically, to assess the differences between the D3R and D2R inactive structures, we considered all possible D3R-D2R pairs: 3PBL-6CM4, 3PBL-6LUQ, and 3PBL-7DFP. After subtracting the metrics of the D2R structure from those of the D3R structure in each pair, we categorized the resulting differences. By averaging the derived categories across these pairs, we identified their consistent differences in the inactive states (**Figs. S13-S15**). Similarly, we compared the active structures of these two receptors, specifically 7CMU-7JVR and 7CMV-7JVR, to detect the trends in the active states (**Figs. S16-S18**). To identify consistent difference patterns in both the inactive and active states, we averaged the categories from these two states (**Figs. S19-21**).

While the D3R and D2R are highly homologous in their TM domain, noticeable differences are present in their sequence alignment. Specifically, the regions of TM1m-TM1i, IL2-TM4i, TM5e, and TM6e-EL3-TM7e contain multiple nonconserved residues with diverse physicochemical properties (**Table S2**). Notably, both TM6e and TM7e harbor divergent residues (positions 6.53, 6.56, and 6.59 in TM6e, and 7.33 and 7.38 in TM7e), while they are connected by highly divergent EL3 that features only two common disulfide bonded cysteines. It is anticipated that somewhat different structural conformations can be originated from these regions.

On the intracellular side, in the comparison of the inactive structures of the D3R and D2R, which do not involve interactions with any other protein, we observed longer distances of TM4i to TMs 3i, 5i, and 6i in the D3R at the DDthreshold of 1.0 Å in the subsegment analysis (**Fig. S15**). However, surprisingly, when comparing the active structures of the two receptors, which are all in complex with the Gi protein, only the TM2i-TM6i distance was found to be longer in the D3R at the same threshold (**Fig. S18**). The smaller divergence on the intracellular side in the active state suggests that binding to the same Gi protein may have a conforming effect, masking some of the divergences between the receptors. Combining the results of the inactive and active states together, there is no common difference among the intracellular subsegments or among the middle subsegments at the DDthreshold of 1.0 Å (**Fig. S21**).

On the extracellular side, the subsegment analysis revealed common differences between the D3R and D2R structures in both inactive and active states. Specifically, TM6e in the D2R has longer distances to TM3e and TM4e, indicating a more outward tilting of TM6e in the D2R compared to the D3R (**Fig. 2A**), while TM7e also demonstrates a slight inward movement at lower DDthreshold (**Fig. S21**). In the ligand binding residue analysis, we found a consistent trend showing that the TM6e residues (6.55-6.59) exhibit longer distances to various ligand binding residues, while Trp^7.40^ in TM7e displays shorter distances to many other residues in the D2R (**Figs. 2B and S19**). To further investigate the tilting of the extracellular subsegments, we measured the bend angles of the conserved proline kinks (Prokinks) near Pro^5.50^, Pro^6.50^, and Pro^7.50^.^29, 30^ We found that the TM6 Prokink of the D3R has significantly smaller bend angles than that of the D2R in both the inactive and active states, with the differences of -11.7° and - 16.9°, respectively. In addition, the bend angle of the TM7 Prokink in the active state is noticeably large in the D3R structures than in the D2R structure, with an average difference of 6.8° (**Table S3**).

**Figure 2.**
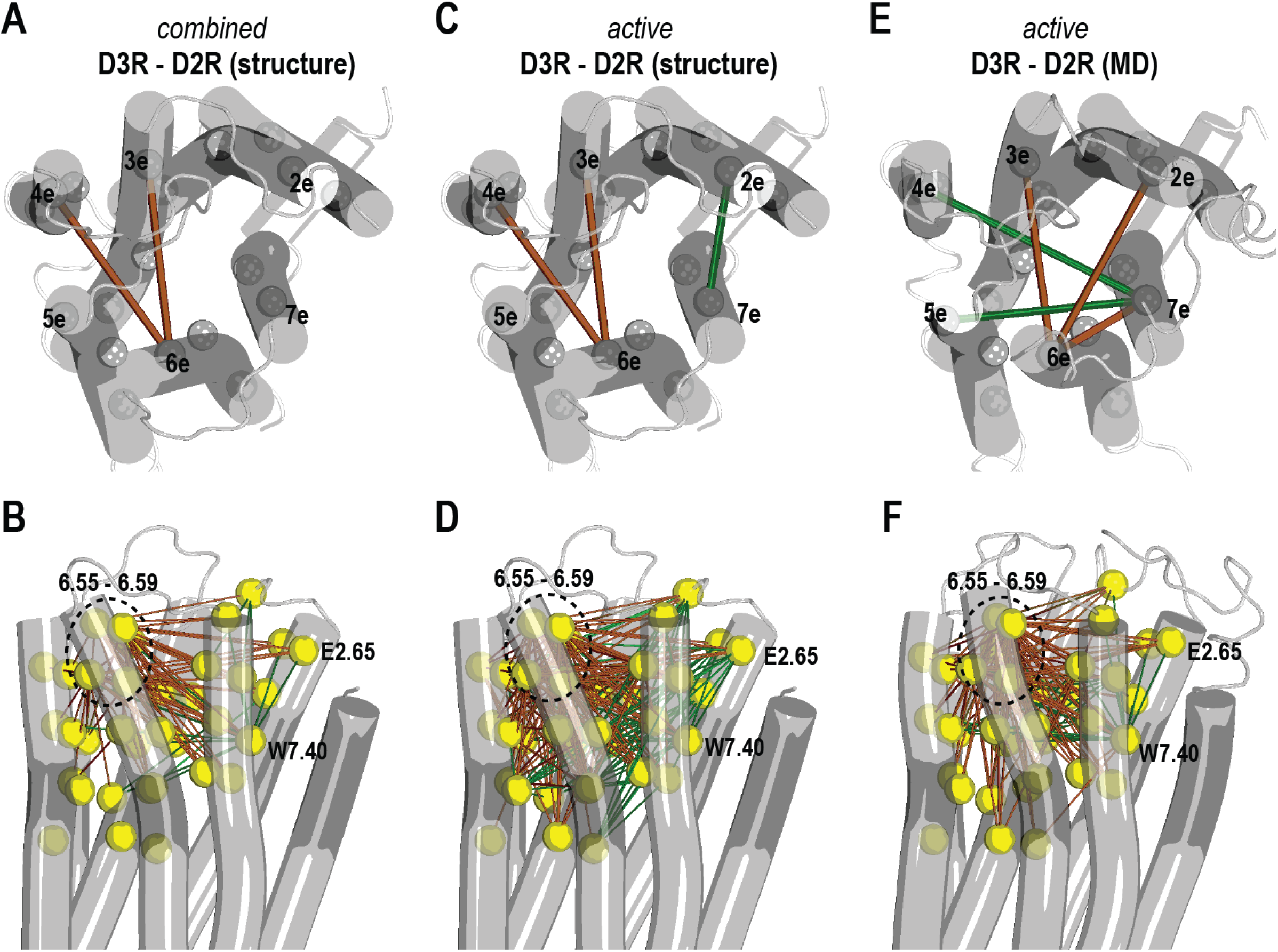
Differences between the experimentally determined D2R and D3R structures, active structures and corresponding MD simulations. Panel A shows the common distance differences among extracellular subsegments between the D3R and D2R by combining the results of both the inactive and active structure comparisons. The mapping was generated according to the results using a DDthreshold of 1.0 Å (see **Fig. S21**), and on a D3R structure in the active state (PDB 7CMU). The COMs of the subsegments are shown with the gray spheres. The green and brown lines indicate larger distances observed in the D3R and D2R, respectively. In panel B, the categorized pairwise distance differences among ligand binding residues using a DDthreshold of 1.0 Å (see **Fig. S19**) are mapped on the same structure. The color scheme employed is the same with that of panel A. The yellow spheres represent the Cα atoms of ligand binding residues. The mapping of the comparison results specifically on the active structures of the D3R and D2R using the same DDthreshold (see **Figs. S18 and S16**) are shown in panels C and D, with the same color and representation schemes. The mapping of the comparison results of the corresponding simulations of these active structures (**Figs. S28 and S26**) are shown in panels E and F.

The observation of a more outward tilting of TM6e in the D2R in both the active and inactive states suggest that this feature may be independent of the LCCs required to accommodate different ligand scaffolds. It is tempting to speculate that it is an intrinsic divergence between the D2R and D3R. However, our previous MD simulations showed that when a D2R inactive model was bound with eticlopride, a nonselective antagonist that binds to the D3R inactive structure (3PBL), TM6e exhibited an inward tilting compared to the situation when the D2R was bound with risperidone (6CM4).^18^ This finding suggests that the conformation of TM6e can vary depending on the specific ligand bound to the receptor. Furthermore, in the active D2R structure (7JVR), we found that the accommodation for the bulky 2-methylpropyl group of the bound nonselective agonist, bromocriptine, which interacts closely with His^6.55^ in TM6e, may contribute partially to the observed outward tilting of TM6e. Therefore, LCCs may coincidentally lead to similar outward tilting of TM6e of the D2R in both inactive and active states, which may not necessarily reflect the either the direction or the extent of the intrinsic divergence between the D2R and D3R.

### MD simulations revealed similar trends observed in the active structures but nuances in a lipid bilayer environment and at 310 K

To specifically gain insights into the active conformations of the D2R and D3R in a more physiologically relevant environment, we built models based on the active-state structures of these two receptors in complex with Gi protein, immersed the models in the lipid bilayer environment, and performed extensive MD simulations at 310 K (see Methods). We first investigated the models bound with the agonists present in the experimentally determined structures, namely D2R/Gi-bromocriptine, D3R/Gi-pramipexole, and D3R/Gi-PD128907 **(Table S4**). Overall, the simulations showed that the protein conformations and the binding modes of the bound agonists revealed by the cryo-EM structures were highly stable. To assess the potential influence of different environments as well as temperature on the observed structural differences between the D2R and D3R, we carried out similar analysis of the simulations results as those for the experimentally determined structures (See Methods).

At the subsegment level, we observed similar trends between the active-state experimental structures and their corresponding MD simulations (**Fig. 2C,E**). Specifically, only limited differences were found between these two receptors among the intracellular subsegments. On the extracellular side, we found long distances of TM1m to several extracellular segments, the outward tilting of TM6e, and a slight inward movement of TM7e in the D2R (**Fig. 2C,E**). However, the directions of the TM1m, TM6e, and TM7e rearrangements are not exactly the same in the experimental structures and the MD simulations (**Fig. 2C,E**). For example, in the case of TM6e, rather than having longer distances to TM3e and TM4e in the D2R compared to the D3R active structures, it has longer distances to TM2e and TM3e in the D2R than the D3R in the simulations.

In the comparison of the Prokinks, we found significant bend angle differences of both the TM6 and TM7 Prokinks between the D3R and D2R in this set of simulations, which were comparable to those between the corresponding D3R and D2R active structures (**Fig. S22, Table S3**).

In the analysis of ligand binding residues, the most prominent differences are that positions 6.55-6.59 are farther away from most of other ligand binding residues in the D2R compared to the D3R. Other obvious disparities include longer distances of Tyr^7.35^ and shorter distances of Trp^7.40^ to many other ligand binding residues in the D2R (**Fig. 2D,F**). However, similar to the subsegment analysis, nuances can be observed between the experimental structures and the simulations. In particular, the longer distances of TM2e residues (positions 2.61-2.64) to those of TM6m (6.44-6.48) in the D2R in the comparison of the experimental structures were not detected in the simulations (**Fig. 2D,F**).

The nuances between the comparison results of experimental structures and the MD simulations are likely due to that the simulations were carried out at 310 K and lipid environment. As the experimental structures were determined at lower cryogenic temperature (<123 K), the higher temperature used in the simulations might amplify some divergences. These amplified divergences may not be obvious in the experimental structures yet could overshadow some of those initially observed in those structures during the simulations. In addition, the adaptations of these two receptors to the lipid environment could differ, which may contribute to the discrepancy as well. On the other hand, this set of simulations did not provide further clues to discern the contributions of the LCC and intrinsic divergence to the observed differences. We need to eliminate the impact of LCC by comparing the D3R and D2R active models bound with the same nonselective full agonist.

### Identification of the binding pose of the nonselective full agonist quinpirole at the D3R and D2R

To further differentiate the divergence between the D2R and D3R especially in the ligand binding pocket, we modeled and simulated the D3R/Gi and D2R/Gi models in complex with a high-affinity agonist quinpirole, known for its non-selectivity and full efficacy at the D2R and D3R.^22, 26^ Depending on the direction of protonation of the pyramidal nitrogen, quinpirole has two diastereomers (5S- and 5R-isomers, **Fig. 3A,B**). While these two isomers are in an equilibrium in solution, the ionic interaction between the pyramidal nitrogen and Asp^3.32^ of the D3R or D2R would position the ligand in different orientations in the ligand binding pocket, and therefore one isomer is likely preferred. We first carried out a docking analysis of both the 5S- and 5R-isomers at the D3R (see Methods). Interestingly, the docking poses of both diastereomers in the D3R appeared to be feasible. Thus, to evaluate which isomer is preferred, we carried out extensive MD simulations of the D3R/Gi complex bound with either of these two isomers to evaluate which isomer is more stable (see Methods and **Table S4**).

**Figure 3.**
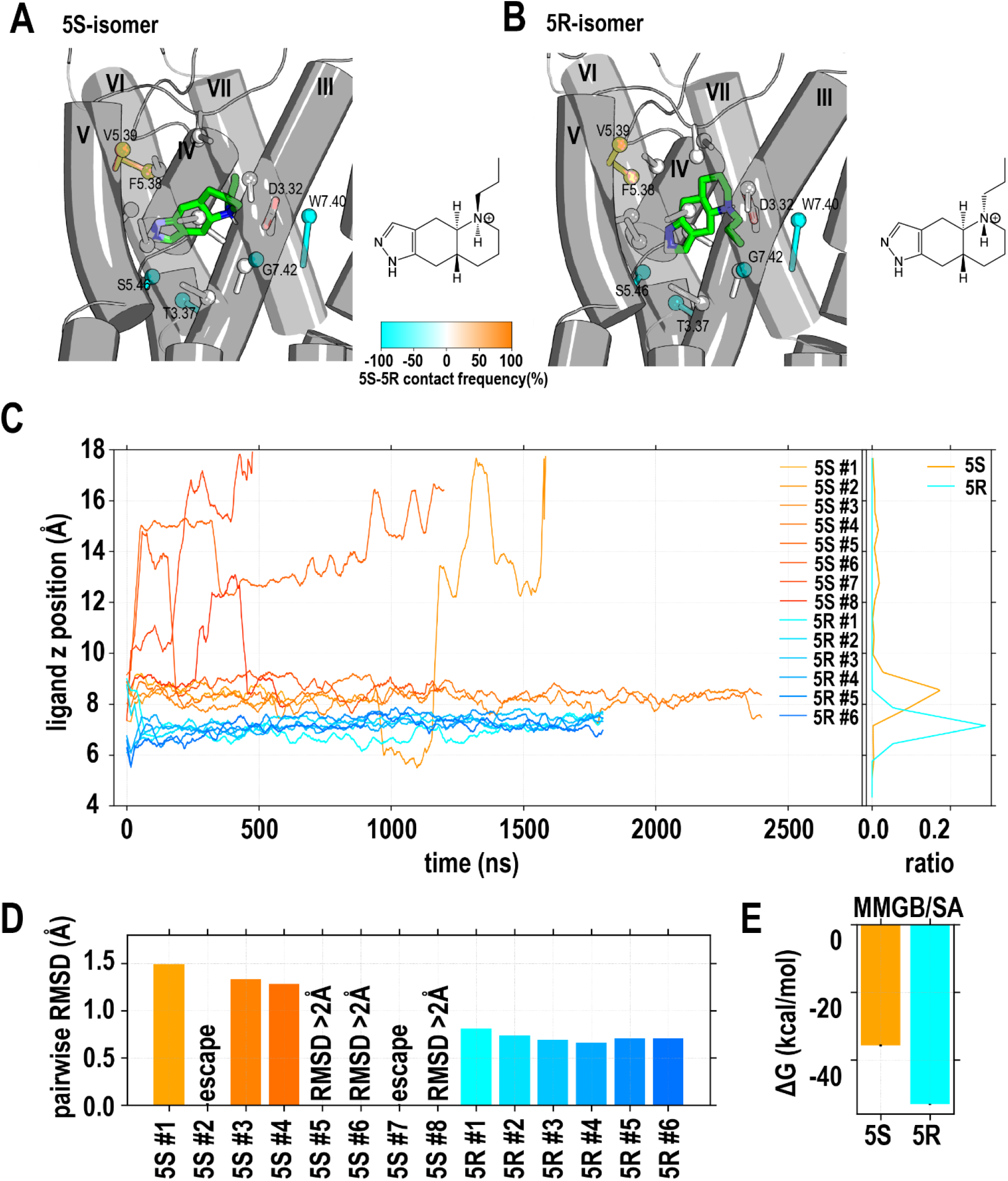
Quinpirole prefers to be in the 5R-isomer in the ligand binding site of D3R. Two possible protonation states of the pyramidal nitrogen of quinpirole, the 5S-isomer (A) and 5R-isomer (B), result in different orientations in ligand the binding pocket, when this nitrogen forms a hydrogen bond with Asp^3.32^. Orange and cyan indicate the residues with higher contact frequencies in the 5S- and 5R-isomer (see **Table S5**). The sticks denote the vectors from the Cα atom of a residue to the COM (represented as a sphere) of the heavy atoms of the sidechain. In panel C, the evolutions of the z positions (i.e., along the axis perpendicular to the membrane) of the ligands in both 5S-isomer and 5R-isomer are shown, while the combined distributions of the z positions for these two isomers are shown on the right. When the binding pose of the 5S-isomer was relatively stable, it still exhibited higher z position values compared to the 5R-isomer. The averaged pairwise ligand RMSDs for each MD trajectory are presented in panel D, indicating that the 5R-isomer was more stable than the 5S-isomer. In panel E, the results of the MMGB/SA calculations using the stable trajectories of the 5S- and 5R-isomers are shown, demonstrating the 5R-isomer was bound tighter than the 5S-isomer.

For both the D3R/Gi-5R-quinpirole and D3R/Gi-5S-quinpirole complexes, we collected multiple trajectories. In all our six D3R/Gi-5R-quinpirole trajectories, the bound 5R-quinpirole quickly relaxed and converged to a define pose, with the pairwise ligand root mean square deviation (RMSD) < 1.0 Å. In contrast, 5S-quinpirole was not stable in all eight trajectories, including two trajectories in which the ligand escaped from the OBS after having lost the interaction with Asp^3.32^ (**Fig. 3C, movies 1 and 2**), while the other six trajectories have noticeably larger pairwise ligand RMSDs than those of the D3R/Gi-5R-quinpirole trajectories (**Fig. 3D**). The preference of the 5R-isomer of quinpirole in the D3R binding pocket was further confirmed by the estimations of the ligand binding free energy with MM/GBSA calculations (see Methods), which showed that the binding energy of the 5R-isomer is approximately 15 kcal/mol lower than that of the 5S-isomer at the D3R (**Fig. 3E**). Interestingly, when we examined the stabilized poses of the 5R- and 5S-isomers in the D3R, the 5R-isomer made more interactions with the residues deeper in the binding pocket, such as Thr^3.37^, Ser^5.46^ and Gly^7.42^, while the 5S- isomer interacted more with more extracellularly located Phe^5.38^ and Val^5.39^ (**Fig. 3A,B and Table S5**).

Similarly, in a set of MD simulations for the D2R/Gi complex bound with either of the two isomers, we found a similar trend that the 5R-isomer was preferred and could be stable in the binding pocket, while the 5S-isomer had a significant chance to escape from the binding pocket or had higher pairwise ligand RMSDs when stayed (**Fig. S23** and **Table S5**).

### The divergence between the D3R and D2R revealed by binding to quinpirole

Between the conformations of the D3R and D2R bound with the same nonselective agonist quinpirole, we assumed that the impact of LCC can be canceled out, and the remaining differences observed would be solely attributed to the intrinsic divergence between the D3R and D2R. Thus, using the representative frame ensembles of the D3R/Gi-5R-quinpirole and D2R/Gi- 5R-quinpirole MD simulations, we carried out similar comparative analyses of subsegments and ligand binding residues as above **(Figs. S26-28)**.

As expected, the subsegment analysis showed that the distance differences are predominant associated with the regions of the D3R and D2R with divergent primary sequences (**Fig. 4A**, **Table S2**). In particular, the diversity in TM1m, where only four of nine residues are conserved, contributed to its longer distances to most of other extracellular and middle subsegments in the D2R. Interestingly, this TM1m-related divergence between the two receptors was surprisingly less pronounced in the comparisons of the active D3R and D2R structures and MD simulations when they are bound with agonists of different scaffolds (**Fig. S28**, note that these comparisons are collectively referred to as the “different-scaffold comparisons” below). Similarly, the divergent TM5e residues are likely responsible for the shorter TM4e-TM5e distances, which was interestingly not observed in the “different-scaffold comparisons”. Thus, the receptor accommodations to the different ligand scaffolds can obscure the intrinsic divergence.

**Figure 4.**
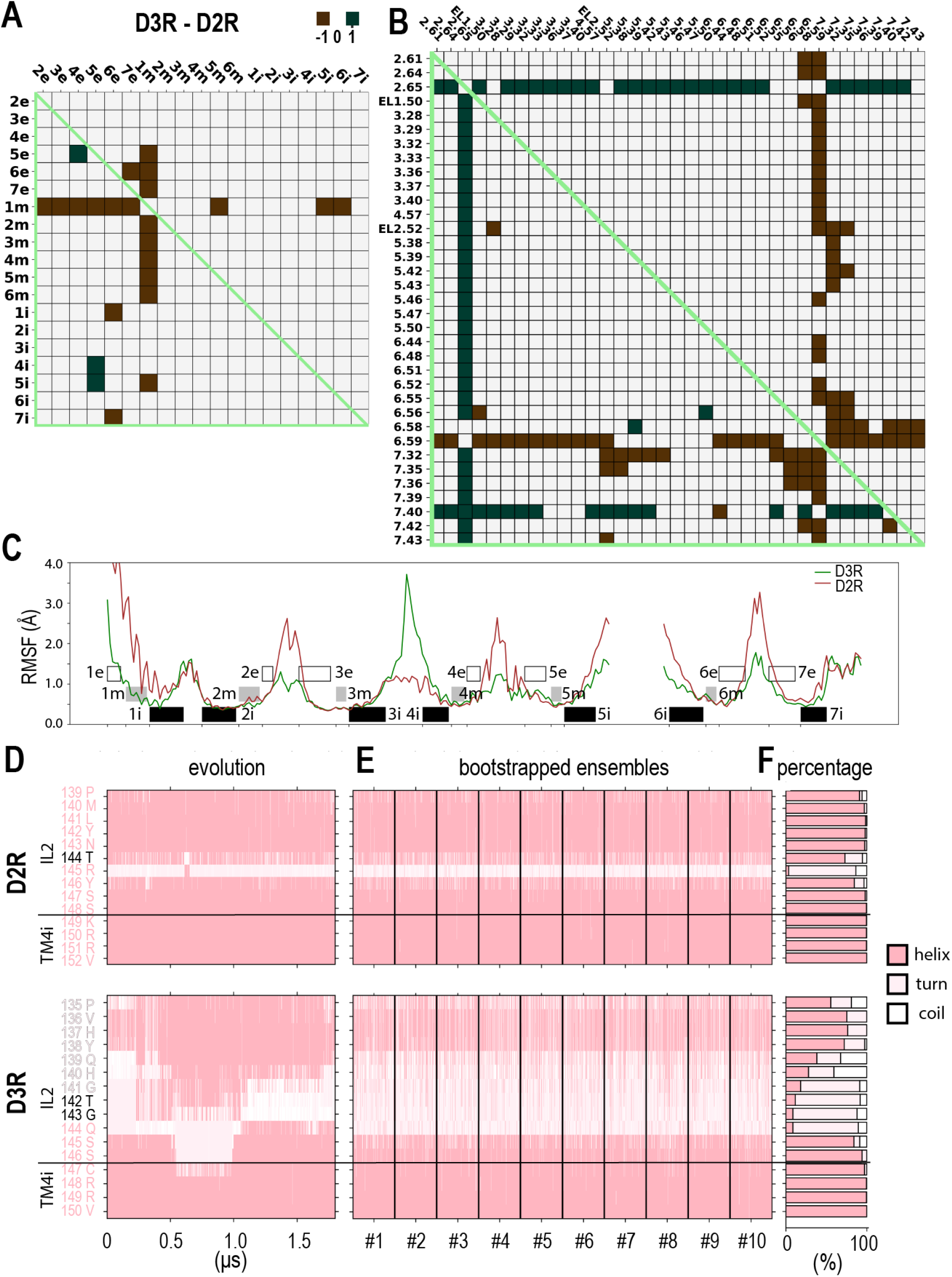
Intrinsic divergence between the D2R and D3R revealed from the comparisons of the 5R-quinpirole bound MD simulations. Panels A and B are heatmap plots showing the categorized pairwise distance differences among the subsegment (A) and ligand binding residues (B) for the D2R and D3R, according to the results using a DDthreshold of 1.0 Å (see **Fig. S28 and S26**). The color and representation schemes are the same as Fig. 2. In panel C, the RMSFs along the indicated regions are shown. The calculations used the bootstrapped frames from D2R/Gi-5R-quinpirole and D3R/Gi-5R- quinpirole simulations. Black shades denote intracellular subsegments, gray shades indicate middle subsegments, and the white box represent extracellular subsegments. In panels E to F, the secondary structures of the IL2-TM4i regions are shown for the evolution along representative trajectories (E), ten bootstrapped frame ensembles (F), and the percentages of the identified secondary structures in the ensembles (G). The secondary structures were calculated with Stride (see Method). The helical, turn, and coil are represented in light pink, lavender blush, and white, respectively.

However, there is no drastic outward tilting of TM6e or the inward movement of TM7e in the D2R as in the “different-scaffold comparisons”. Among the TM6e differences, only longer TM6e- TM7e distances in the D2R is observed at or above the DDthreshold of 0.6 Å (**Figs. S28 and 4A**). On the intracellular side, consistent with the conforming effect of the binding to the Gi protein observed in the “different-scaffold comparisons”, we did not detect noticeable divergence between the two receptors among the intracellular subsegments (**Fig. 4A**).

In the ligand binding residue analysis, we found many of the differences are associated with Glu^2.65^ and Trp^7.40^, two residues conserved between the D2R and D3R. Both Glu^2.65^ and Trp^7.40^ had larger distances to many other binding site residues, including a larger Glu^2.65^-Trp^7.40^ distance itself, in the D3R (**Fig. 4B**). By examining the resulting models from the simulations as well as the experimentally determined structures, we found that the sidechain orientation of Trp^7.40^ exhibited significant differences between the D3R and D2R. Specifically, Trp^7.40^ formed a water-mediated hydrogen bond with Ser^1.38^ of TM1m in the D3R but directly interacted with Glu^2.65^ in the D2R, in which the aligned Leu^1.38^ could not form any H-bond (**Figs. 5A, S25A**). Correspondingly, the χ1 angle of Trp^7.40^ displays distinct distributions, *trans* in the D3R and *gauche+* in the D2R (**Fig. S24B**). Notably, these distributions are consistent with those observed in the experimentally determined D3R and D2R structures, whether in inactive or active states (**Figs. S24A, S25B**).

**Figure 5.**
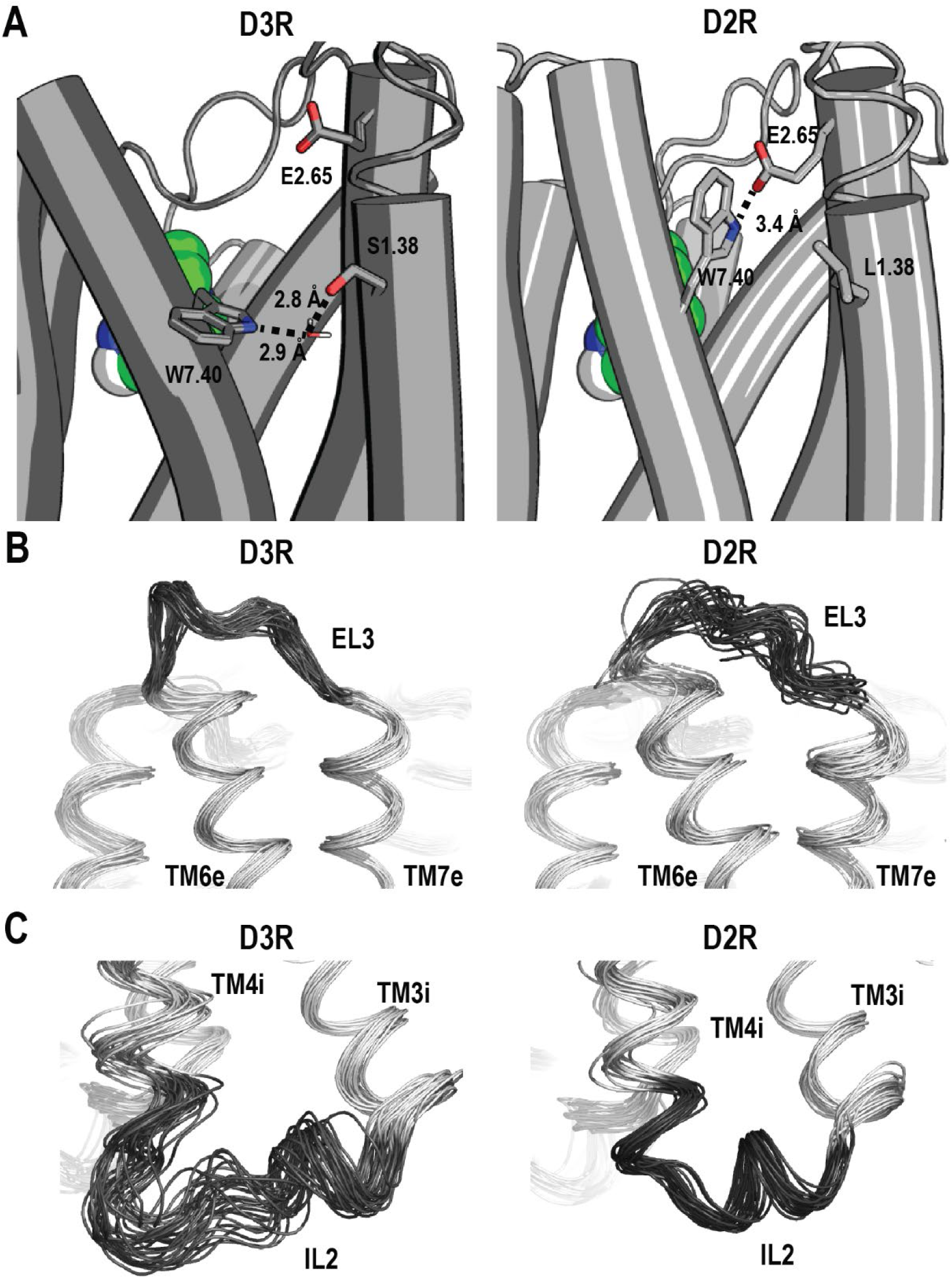
The intrinsic divergence between the D2R and D3R at the selected regions. Panel A shows the divergence at residue position 1.38 leads to distinct interactions with Trp^7.40^ in the D3R and D2R. Specifically, a water-mediated hydrogen bond interaction between Trp^7.40^ and Ser^1.38^ was observed in the D3R, while Trp^7.40^ formed a direct interaction with Glu^2.65^ (Å) in the D2R. Panel B demonstrates that EL3 has a larger fluctuation in the D2R than the D3R, which corresponds to the comparisons of their RMSFs (Fig. 4C) and the distributions of the bend angles of TM6 and TM7 Prokinks (**Fig. S22**). Panel C illustrates that in the IL2-TM4i region, IL2 exhibited a predominantly helical conformation in the D2R, while oscillated between coil and helical conformations in the D3R, the dynamics of which propagated to the TM4i region.

In addition, residues 6.58, 6.59, 7.32, and 7.35 are associated with some smaller distances in the D3R (**Fig. 4B**), consistent with the divergent sequences of the TM6e-EL3-TM7e region between these two receptors. In addition to the highly divergent EL3, this region includes several non-conserved residues (**Table S2**). In particular, the residues located at the 6.59, 7.33, and 7.38 are of distinct physiochemical properties. Nevertheless, the bend angle difference of the TM6 Prokink was -5.7° for the D3R/Gi-5R-quinpirole versus D2R/Gi-5R-quinpirole comparison (**Fig. S22, Table S3**), which was drastically smaller than the “different-scaffold comparisons”. Specifically, the angle is 20.0° in D2R/Gi-5R-quinpirole, which was smaller than 22.4° of D3R/Gi-PD128907 (**Table S3**). These results suggest that while TM6e of D2R may still tend to be more outward tilting than that of the D3R, the extent is much smaller than that revealed by the experimentally determined structures. Furthermore, the bend angle difference of the TM7 Prokink was -3.8° for the D3R/Gi-5R-quinpirole versus D2R/Gi-5R-quinpirole comparison, which was in an opposite direction to that in the “different-scaffold comparisons” (**Table S3**).

### The D3R and D2R exhibit distinct dynamics

The strength of MD simulations lies in their ability to offer insights into potentially functionally relevant protein dynamics beyond what can be revealed by experimentally determined structures, which essentially capture static snapshots of the conformational states.

To characterize the divergence of the D3R and D2R from a dynamic perspective, we first carried out a root mean square fluctuation (RMSF) analysis of our MD simulation results (see Methods). RMSF quantifies a particle’s fluctuations from its mean position over time, whereas RMSD measures the average deviation between different structural conformations. Notably, our RMSF analysis results showed that the middle subsegments, including TMs 3m, 5m, and 6m that form our recently proposed activation switch,^14^ display overall less fluctuation compared to the extracellular subsegments (**Fig. 4C**). In addition, our results demonstrated that EL3 had a higher RMSF peak value of ∼3.3 Å in the D2R/Gi-5R-quinpirole condition, compared to ∼1.7 Å in D3R/Gi-5R-quinpirole, indicating that EL3 as well as neighboring TM6e and TM7e are more dynamic in the D2R (**Figs. 4C, 5B**). In contrast, on the intracellular side, IL2 and its neighboring TM4i exhibited higher RMSF and were therefore more dynamics in the D3R compared to the D2R (**Fig. 4C**).

We then analyzed the secondary structure for each residue of IL2 in our simulated D3R and D2R conditions (see Methods). We observed that IL2 displayed a greater tendency to retain helical conformation in the D2R compared to the D3R, in which IL2 of the D3R oscillates between helical and non-helical (including turn and coil) forms (**Figs. 4D, 5C**). Specifically, compared to the aligned portion in the D2R, residues 135-144 of the D3R exhibit a weaker tendency to adopt a helical conformation. The stronger tendency of IL2 to be in loop conformations corresponds with the higher IL2 RMSF observed in the D3R (**Fig. 4C**).

Remarkably, upon scrutinizing the Prokink distributions of TMs 5, 6, and 7 across all simulations and comparing those of the D2R and D3R (**Fig. S22A-F**), a notable observation emerged: the distributions in the D2R are all notably broader than their counterparts in the D3R. Given that the intracellular portions of these TMs engage in direct interactions with the G_i_ protein and are stabilized by it, this trend strongly indicated that the extracellular portions of these TMs, which are main contributors to the ligand binding site, was intrinsically more flexible in the D2R compared to those in the D3R.

To further validate this finding, we assessed the flexibility of the ligand binding site through the calculations of pairwise RMSDs of the frames within each representative ensemble of the simulated D3R and D2R conditions (**Fig. S22G,H**). Our results demonstrated wider distributions of the pairwise RMSDs in the D2R, suggesting that the ligand bind site of the D2R exhibited greater flexibility than that of the D3R. This difference was particularly pronounced in the comparison between the D3R/Gi-5R-quinpirole and D2R/Gi-5R-quinpirole conditions when both receptors were bound with the same ligand (**Fig. S22H**).

## Discussion

When rationally optimizing a ligand to achieve a desired pharmacological profile using a receptor structure, in addition to retaining distinctive receptor characteristics related to the targeted functional state, it is also important to preserve or improve its selectivity against other homologous receptors. However, both ACCs and intrinsic divergences can be obscured by LCCs.

In this study, we first used distance-based metrics to identify ACCs in the D2R and D3R. Based on our recent study at the aminergic family level,^14^ our current analyses focused on the ACCs that may be specific for these two highly homologous receptors, while also taking the protein dynamics into considerations. In addition to the common activation-related rearrangements observed for many other GPCRs,^14^ our analyses unveiled ACCs unique for these two receptors. Specifically, TM5e exhibited a departure from TM2e and TM3e, while TM6e also moved away from TM2e, in accordance with the movement of His^6.55^ and Asn^6.58^ away from Val^2.61^ and/or Leu^2.64^ (**Fig. 1**). In addition, the shifts of Ser^5.42^ away from Cys^3.36^, and Ser^5.43^ towards to Thr^3.37^ and Ile^3.40^, indicate a subtle rotation of TM5e. This rotation appears to be independent of whether an agonist can form H-bonds with these serine residues, although such interactions may facilitate the rearrangement. Despite the pivotal significance attributed to TM5 serine residues for catecholamine receptors,^31, 32^ our comprehensive analysis of entire aminergic family did not uncover any consistent rotamer changes of TM5 serine and threonine during activation.^14^ This absence of a common trend aligns with the lack of necessity for an agonist to form H-bonds with these serine residues. Rather, the key functional role of TM5 serine residues likely lies in their ability in modulating the backbone conformation during the transition from the inactive to active state.^33^

By adapting the same metrics and analysis protocol, we subsequently examined the experimentally determined structures to discern disparities between these two receptors in both the inactive and active states. Given our grounding in the understanding of their inactive states,^13, 18^ our primary focus in this investigation centers on the differences in their active states, for which we carried out MD simulations as well. While on the intracellular side, the limited intracellular divergence may be due to the masking effect of their coupling to the same Gi protein, on the extracellular side, we observed a more pronounced outward tilting of TM6e in the D2R compared to the D3R in both the experimentally derived structures and corresponding simulations. It is crucial to emphasize that these active structures are bound with ligands of markedly different scaffolds, and we collectively term these comparisons, encompassing both the structures and the corresponding simulations, as the “different-scaffold comparisons”. However, in the MD simulations involving the D3R and D2R models bound with the nonselective full agonist quinpirole – a condition that may mitigate the influence of LCC in comparing these two receptors – the extent of outward tilting observed in TM6e of the D2R was significantly reduced. This suggests that the extent of demonstrated difference in the “different-scaffold comparisons” is coincidently and misleadingly more than the intrinsic divergence.

In contrast, TM1m exhibited notably greater disparity between these two receptors in our quinpirole-bound D3R and D2R simulations than in the “different-scaffold comparisons”, suggesting that the LCCs could also obscure the intrinsic divergence (**Fig. S28**). This observation echoes our prior finding when comparing the β_1_ adrenergic receptor structures bound with same (partial) agonists, where we demonstrated the ACCs common to the aminergic family could be detected at higher DDthreshold, when LCCs could be cancelled out.^14^ Regardless of whether it is obscured, in all the D2R versus D3R comparisons, the impact of the divergence propagating from the nonconserved TM1m to a functionally important conserved region is evident. Specifically, our analyses revealed the formation of an interaction network involving a water-mediated interaction between Ser^1.38^ and Trp^7.40^ in the D3R, while in the D2R, Trp^7.40^ engaged in a direct interaction with Glu^2.65^ (**Fig. 5**). Note that the varied lengths of EL1 that connects TM2 and TM3 may also facilitate different orientations of Glu^2.65^ in the D3R and D2R.^13, 34^ Both Trp^7.40^ and Glu^2.65^ are conserved residues that contribute to the secondary binding pocket at the interface of TMs 1, 2, 3, and 7 of D2R and D3R.^34, 35^ Thus, although the ligand binding residues of the D2R and D3R are highly conserved, there are opportunities to be exploited for the development of selective compounds, such as the bitopic scheme employed in the development of the D3R-selective ligands.^13^

On the intracellular side, the intrinsic divergences of IL2 between the D2R and D3R lead to varying propensities to adopt a helical conformation. Consequently, the distinct conformations and dynamics exhibited by these two receptors in this region, which directly interacts with G proteins, may underlie their coupling specificity,^36^ as well as unique signaling characteristics.^9^ Notably, the intrinsic conformational flexibility of the IL2 has also been observed in a few other homologous aminergic receptors. This flexibility enables it to transition between discrete conformations in response to various cellular signals and ligands to modulate the functional properties of these GPCRs.^37, 38^

The functionally relevant structural divergence may not only exist in the averaged conformations that can be experimentally determined, but also in their dynamic properties that can be characterized through MD simulations. In this study, our simulations and the following analyses unveiled a heightened flexibility of the extracellular portions of TMs 5, 6, and 7 in the D2R than those of the D3R. Notably, we found that the flexibility of this region is associated with the plasticity of the ligand binding site in these two receptors, i.e., the ligand binding site of the D2R is more flexible than that the D3R. Curiously, upon querying the ChEMBL database, we found a significantly larger number of the D3R selective ligands in comparison to the D2R selective compounds (Won et al., manuscript in preparation). Assuming there have been comparable levels of efforts directed towards enhancing both the D3R and D2R selectivities, we speculate that the potentially more rigid D3R binding pocket may account for this distinction.

In this study, we applied an established structural analysis protocol on both experimentally determined structures and MD simulations, to unravel the ACCs of and the intrinsic divergence between D2R and D3R from the LCCs. Our findings will serve as a foundation for the targeted design and optimization of drugs that specifically target D2R and D3R, potentially offering more effective and tailored therapeutic interventions.

## Methods

### Definitions of the structural elements

We defined various structural elements at different locations and resolutions.

#### Transmembrane subsegments

TM1e (the extracellular section I of TM1, 1.30-1.36), TM1m (the middle section (m) of TM1, residues 1.37-1.45),TM1i (the intracellular section (i) of TM1, residues 1.46-1.59), TM2i (residues 2.38-2.51), TM2m (residues 2.52-2.60), TM2e (residues 2.61-2.66), TM3e (residues 3.22-3.35), TM3m (residues 3.36-3.40), TM3i (residues 3.41-3.55), TM4i (residues 4.39-4.49), TM4m (residues 4.50-4.55), TM4e (residues 4.56-4.62), TM5e (residues 5.36-5.45), TM5m (residues 5.46-5.50), TM5i (residues 5.51-5.63), TM6i (residues 6.30-6.43), TM6m (residues 6.44-6.48), TM6e (residues 6.49-6.60), TM7e (residues 7.32-7.43), TM7i (residues 7.44-7.54).

#### Extracellular ends (EE)

We also defined the following residues to be the extracellular ends for each TM: TM1ee (residues 1.30-1.33), TM2ee (residues 2.63-2.66), TM3ee (residues 3.22- 3.25), TM4ee (residues 4.59-4.62), TM5ee (residues 5.36-5.39), TM6ee (residues 6.57-6.60), TM7ee (residues 7.32-7.35), and EL2 (residue EL2.52).

#### Binding site residues

The binding site residues identified in our previous studies ^1^ include 2.61, 2.64, 2.65, EL1.50, 3.28, 3.29, 3.32, 3.33, 3.36, 3.37, 3.40, 4.57, EL2.52, 5.38, 5.39, 5.42, 5.43, 5.46, 5.47, 5.50, 6.44, 6.48, 6.51, 6.52, 6.55, 6.56, 6.58, 6.59, 7.32, 7.35, 7.36, 7.39, 7.40, 7.42, and 7.43. Among them, 3.32, 3.33, 3.36, EL2.52, 5.42, 5.46, 6.48, 6.51, 6.52, 6.55, 7.39, 7.43 are the residues located in the orthosteric binding site (OBS).

### Superposition-independent structural analysis

To detect conformational differences between a pair of structures, we developed several analyses based on pairwise distance of various structural elements defined above.^14^

In each distance analysis for each pair of structures of the same receptor, we first calculated the pairwise distances of all possible pairs of the structural elements in each structure (when a structural element includes more than one heavy atom, we use the center of mass (COM) of included atoms for this calculation), and then computed the differences of the corresponding distances between the two structures. For a pair of inactive and active structures in the ACC analysis, it is calculated as following:

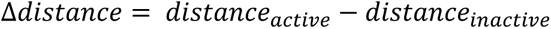

For a pair of the D3R and D2R structures (in the same state) in the intrinsic divergence analysis, it is calculated as following:

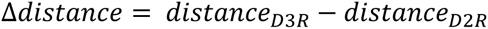

### Classification and integration of the analysis results

We classified the Δ*distance* according to a threshold (referred to as distance-difference threshold) into categories “1”, “0”, and “-1”. Specifically, in the ACC analysis, “1” or “-1” represents that the active structure has a larger or smaller distance than the inactive structure, respectively, when the absolute value of Δ*distance* (|Δ*distance*|) between two structures is larger than the distance-difference threshold (DDthreshold). In the intrinsic divergence analysis, “1” or “-1” represent the D3R structure has a larger or smaller distance than the D2R structure, respectively, when |Δ*distance*| > DDthreshold. 0 means the Δ*distance* is smaller than or equal to the threshold.

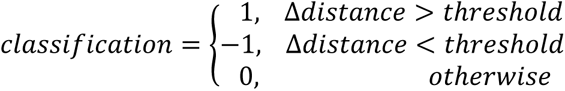

In this study, the distance-difference threshold values were chosen to be 0.0, 0.2, 0.4, 0.6, 0.8, and 1.0 Å.

This classification is necessary when we average the results from all possible pairs of structures of the same comparison (e.g., all the pairs of a receptor for the ACC analysis or the all the inactive pair between the two receptors in the intrinsic divergence analysis) to derive the trend, because without using categories, for example, a very large positive distance difference may cancel out several small negative ones resulting in a misleading conclusion.

For each distance analysis, we carried out both with the backbone Cα atoms and with the sidechain heavy atoms, the results of which are shown on the top-right and bottom-left halves of the same heatmap, respectively. It is important to note that these analyses are independent of how the structures are superimposed. While the specific distances do not have directional information, by calculating the differences of these distances between two conditions and mapping them on a heatmap, we can deduce the trend of rearrangements.

To highlight the common trends on a heatmap showing the categorized distance differences, we applied masks on the corresponding results, with a category-threshold of 0.9 (but not 1.0, to tolerate some uncertainty in the structures, see Discussion). Thus, if the category average of the difference of a given measurement is smaller or equal to 0.9, it is colored as white in the masked distance difference heatmaps.

In the ACC analysis of the D2R and D3R, we considered three D2R pairs: 7JVR-6CM4, 7JVR-6LUQ, and 7JVR-7DFP, and two D3R structure pairs 7CMU-3PBL and 7CMV-3PBL. In the intrinsic divergence analysis of the D2R and D3R, we considered three inactive structures pairs: 3PBL-6CM4, 3PBL-6LUQ, and 3PBL-7DFP, and two active structure pairs 7CMU-7JVR and 7CMV-7JVR.

### Treatment of missing residues, sidechains, and mutations in our analysis of experimentally determined structures

For the structural elements defined in this study (see above), if any residue is missing in a structure, when calculating the COM of the Cα atoms, we used the available Cα atoms of the other residues in the elements. The missing residue(s) for each structure of D2R and D3R are listed in our previous study.^14^ We did not include TM1ee and TM1e in our analysis due to their many missing residues in many structures. Note that no binding site residue in any structure is missing. When a structural element has any missing sidechain heavy atoms, we used the available heavy atom(s) in the element (including the situation when only the Cα atom is available), to calculate the COM. The list of residues missing sidechains for each structure is shown previously.^14^ We did not mutate the residues mutated during the crystallography and cryo-EM studies back to their wildtype residues. The mutations for each structure are shown previously.^14^

### Molecular modeling of the human D3R and D2R in complex with Gi proteins

The cryo-EM structures of the human D3R/Gi complexes (PDB 7CMU and 7CMV) were used as the templates to build the D3R/Gi-pramipexole and D3R/Gi-PD128907 models, respectively. Comparative modeling in MODELLER (version 9.24) was used to add the missing sidechains, and to mutate thermostabilized residues back its wildtype residues.

Of note, in the D3R cryo-EM structure, the N-terminus (residues 1-31) and intracellular loop 3 (IL3) (residues 224-320) were missing. Sufficient number of N-terminal residues can prevent the entry of lipid molecule to the ligand binding pocket in the molecular dynamics (MD) simulations. Therefore, we built part of the missing N-terminus (residues 17-31) using the same region from our previous equilibrated D3R models.^4^ In addition, a previously optimized 9 residues poly-glycine chain was also included in the models to connect TM5 and TM6 to restrain the dynamics resulting from the hanging intracellular ends of these two segments.^4^ For the complexed Gαi subunit, the helical domain (residues 58-176) was not resolved and thus we can only construct the Ras domain with the four thermostabilizing mutations mutated back to the wildtype residues. Final models were selected based on the DOPE score and the proper backbone φ-ψ distribution in Ramachandran plot.

The cryo-EM structure of the D2R/Gi complex (PDB 7JVR) was used as the template to build the human D2R/Gi-bromocriptine model. In the D2R cryo-EM structure, the N-terminus (residues 1-33) and IL3 (residues 226-365) were not resolved. To comparatively investigate the D2R versus D3R, we applied the same modeling strategy as D3R/Gi complexes by adding missing N-terminus (residues 16-33) and a poly-Gly segment (9 residues) in IL3 from previous equilibrated D2R models.^39^ For Gαi subunit, the N-terminus (residues 1-4) and helical domain (residues 56-181) were not solved in structure 7JVR, and we used the N-terminal residues 2-6 from the Gαi subunit in another D2R/Gαi-bromocriptine structure (PDB 6VMS)^19^ as the template. The rat Gβ1 and bovine Gγ2 from the structure 7JVR were used as the templates for modeling human Gβ1 and Gγ2 in the human D2R/Gi model.

### Modeling of the binding poses of quinpirole in the D3R and D2R

The receptor-G protein complexes were taken from the last frame of μs production simulations in D2R/Gi-bromocriptine and D3R/Gi-pramipexole complexes. We then carried out pKa predictions of quinpirole using Jaguar^40^ and Epik^41^ programs in the Schrodinger Suite (version 2021-1). We then conducted docking for both 5S- and 5R-quinpirole to the D3R/Gi complex (PDB 7CMU) using the induced fit docking (IFD) protocol^42^ in Schrodinger. The quinpirole binding orientations were chosen by the IFD score. Subsequently, we use the selected poses in D3R as the references to select IFD docking poses of 5S- and 5R-quinpirole at the D2R/Gi complex (PDB 7JVR). The final binding poses were selected based on the best IFD score obtained.

### Molecular dynamics simulation protocol

The receptor-G protein models with either the originally bound ligands in the cryo-EM structure or the selected binding poses of quinpirole were further processed to build the simulation systems with the Desmond System Builder of Schrodinger suites (version 2021-1 with OPLS4 force field). Briefly, the D3R or D2R complex models were immersed in explicit 1- palmitoyl-2-oleoyl-sn-glycero-3-phosphocholine lipid bilayer (POPC). The simple point charge (SPC) water model was used to solvate the system, the net charge of the system was neutralized by Cl^-^ ions, and then 0.15 M NaCl was added. Residues Asp^2.50^ and Asp^3.49^ are protonated to their neutral forms as assumed in the active state of rhodopsin-like GPCRs.^43^ The process resulted in a system with a dimension of 106×115×148 Å^3^ and total number of atoms of ∼182000. The initial parameters for bromocriptine, quinpirole and PD128907 were further optimized by the force field builder of the Schrodinger Suites (version 2021-1 with OPLS4 force field).

Desmond MD systems (D. E. Shaw Research, New York, NY) was used for the MD simulations. Similar to our previous simulation protocols used for GPCRs,^18^ the system was initially minimized and equilibrated with restraints on the ligand heavy atoms and protein backbone atoms. The NPγT ensemble was used with constant temperature maintained with Langevin dynamics. Specifically, 1 atm constant pressure was achieved with the hybrid Nose-Hoover Langevin piston method on an anisotropic flexible periodic cell with a constant surface tension (x-y plane). In the production runs at 310 K, all restraints on the receptor were released; however, to retain the integrity of the Gi protein while allowing adequate flexibility to interact with the receptor, the heavy atoms of residues 9-30, 35-57, 170-190, 196-214, and 221-307 of Gαi, and the entire Gβ and Gγ subunits were restrained with a force constant of 1 kcal/mol/Å.

For each condition, we collected at least three trajectories starting from different random number seeds. Overall, 43 trajectories with an aggregated simulated time of 69.1 μs were collected (**Table S4**).

### MM/GBSA calculations

The binding free energies between the bound 5S- and 5R-quinpirole in the D3R and D2R were estimated using the Molecular Mechanics/Generalized Born Surface Area (MM/GBSA) calculations. The same force field was employed in the MD simulations for both the proteins and ligands, but with the VSGB2.1 solvation model.^44^ Frames were extracted every 6 ns from the last 600 ns of the production simulations for all the 5S- and 5R-quinpirole bound trajectories. To calculate the binding free energies, the thermal_mmgbsa.py script from the Schrodinger suite was used. The MM/GBSA calculations were carried out on 500 bootstrapped frames for each condition. The reported binding free energies are the averages of these 500 bootstrapping frames for each condition.

### Structural analysis using ensembles constructed through bootstrapping

We aimed to analyze the conformational dynamics of various receptor-ligand complexes, including D3R/Gi-pramipexole, D3R/Gi-PD12890, D2R/Gi-bromocriptine, D3R/Gi-5R-quinpirole, and D2R/Gi-5R-quinpirole. To achieve this, we generated representative ensembles for each complex by employing bootstrapping. Specifically, we randomly selected 1000 frames with replacement from the trajectory data for each condition, and this process was repeated ten times to ensure robustness. These bootstrapped ensembles were then utilized consistently throughout all geometric calculations and analyses. We calculated dihedral angles and distances using VMD-python (version 3.0.6)^45^ and MDAnalysis.^46^ These calculations enabled us to assess the structural variations and interactions in the receptor-ligand complexes under study.

The superposition-independent structural analysis, classification and integration of the analysis results can be applied to MD simulation. For an individual MD frame, we carried out similar analysis as those for the experimentally determined structures by measuring pairwise distances among the defined structural elements. We then averaged the results of bootstrapped representative ensembles of frames from the simulations of each model (referred to as “condition” below), calculated the differences of two conditions being compared, and classified the differences into categories (see Methods).

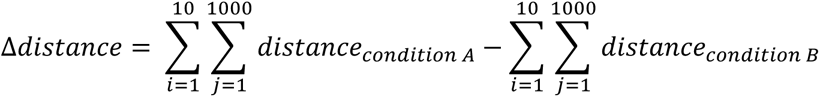

Specifically, we first calculated the difference between the D3R/Gi-pramipexole and D2R/Gi- bromocriptine conditions “D3R/Gi-pramipexole - D2R/Gi-bromocriptine” and “D3R/Gi-PD128907- D2R/Gi-bromocriptine”. We then averaged the derived categories of these two pairs to identify common trends. Similarly, the same category average scheme was used for the condition of “D3R/Gi-5R-quinpirole – D2R/Gi-5R-quinpirole”.

### Conformational analysis

#### Ligand z position and pairwise ligand RMSD

The ligand z position was calculated for each condition of 5S- and 5R-quinpirole in the D3R and D2R simulations. Four different reference coordinates were prepared for protein-ligand complexes and aligned to the same cryoEM reference using seven transmembrane segments. For each condition, the center of mass of ligand heavy atoms was calculated and the z component was reported as the ligand z position. The distribution plot of the ligand z position was using the z position smaller than 12 Å. To calculate pairwise ligand RMSD, the OBS residues in TM (residue positions 3.32, 3.33, 3.36, 5.42, 5.46, 6.48, 6.51, 6.52, 6.55, 7.39, and 7.43) were used to align the whole trajectory for individual run and the pairwise RMSD of ligand heavy atoms was then calculated and the average value and standard error were reported.

#### RMSF calculation

We initially employed the protein Cα atoms of the specific TM regions (residues 63-91, 100-134, 145-170, 186-200, and 362-400 for the D3R and residues 68-96, 104-138, 147-172, 187-201, and 376-414 for the D2R) for structural alignment and then computed the RMSF of bootstrapped ensembles. The RMSF calculation was determined using the MDAnalysis library.^46^ Subsequently, we use the Cα atoms of the residues in the region having RMSF values below 1 Å to re-align the structures and re-compute the RMSF. This iterative process continued until the average value of the region having RMSF values less than 1 Å is converged.

#### Secondary structure determination

Utilizing bootstrapped ensembles of D3R/Gi-5R- quinpirole and D2R/Gi-5R-quinpirole, the estimation of IL2’s secondary structure was assigned. The assignment of IL2’s structural elements was conducted through STRIDE.^47^ After generating IL2 structure assignment for ten bootstrapped ensembles, we subsequently computed an averaged percentage representation of the secondary structure assignment. Note that when categorizing helical assignments, we did not differentiate between α-helix, 310-helix, or π-helix.

#### Prokink analysis

The measurement of Prokink was performed using Simulaid.^48, 49^ The bend angle of the proline kink for TM5, TM6, and TM7 were calculated using the bootstrapped ensembles. The bend angle represents the angle formed between two segments when the helix is kinked along its axis. In the Prokink estimation, the pre- and post-helical segments were defined as the seven residues before and after the proline position for Pro^5.50^ and Pro^6.50^. In case of TM7, the pre- and post-helical segments were defined as the five residues before and after the proline position for Pro^7.50^. We employed an identical averaging approach, consistent with the one employed in structural analyses, for the bend angles and their difference presented in **Table S3**. The average and standard deviation from the bootstrapped ensembles were then calculated for the distribution plots.

#### Pairwise RMSD of ligand binding residues

The same bootstrapped ensembles were used for pairwise RMSD computations. We first aligned this ensemble using the backbone atoms of ligand binding residues defined previously in ref ^1^. We then computed the pairwise RMSD for these ligand binding residues and determined their average and standard deviation from the bootstrapped ensembles for the distribution plots.

## Author Contribution

K.H.L and L.S. designed the study. K.H.L. carried out computational modeling, simulations and analysis. Both authors interpreted the results and wrote the manuscript.

## Acknowledgements

Support for this research was provided by the National Institute on Drug Abuse–Intramural Research Program, Z1A DA000606 (L.S.). This work utilized the computational resources of the NIH HPC Biowulf cluster (http://hpc.nih.gov).

## Supplemental Materials

**Figure S1.**
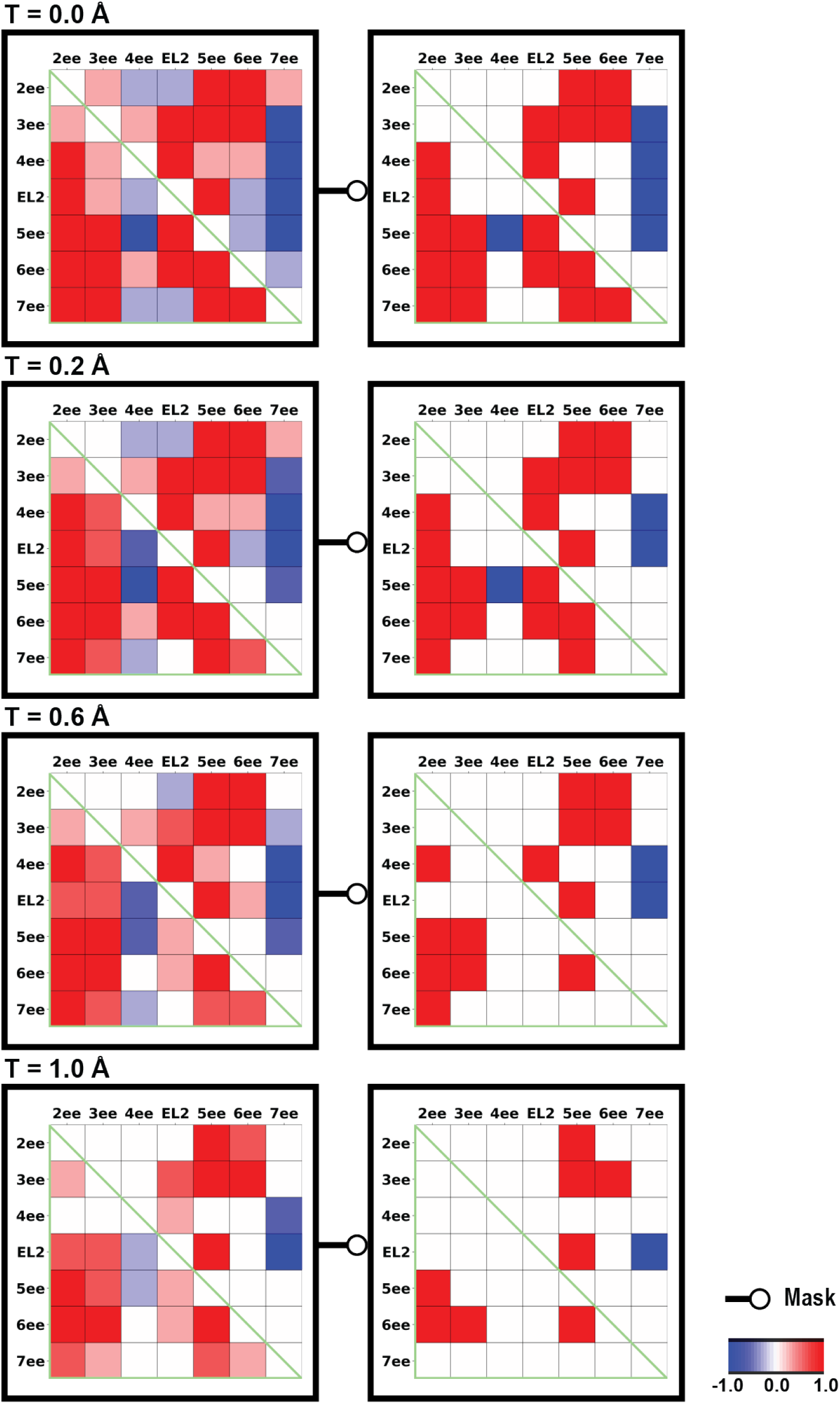
The extracellular-end distance differences of the inactive and active D2R structure pairs. The heatmap shows the difference of pairwise distance among the extracellular-end between the active and inactive structures (D2R active – D2R inactive) and categorized based on various DDthreshold (T). Red represents the active conformation have a larger distance than inactive conformation, and blue represents the inactive conformation have a larger distance than active conformation. If the category average of the difference is smaller or equal to 0.9, it is colored as white in the masked heatmaps.

**Figure S2.**
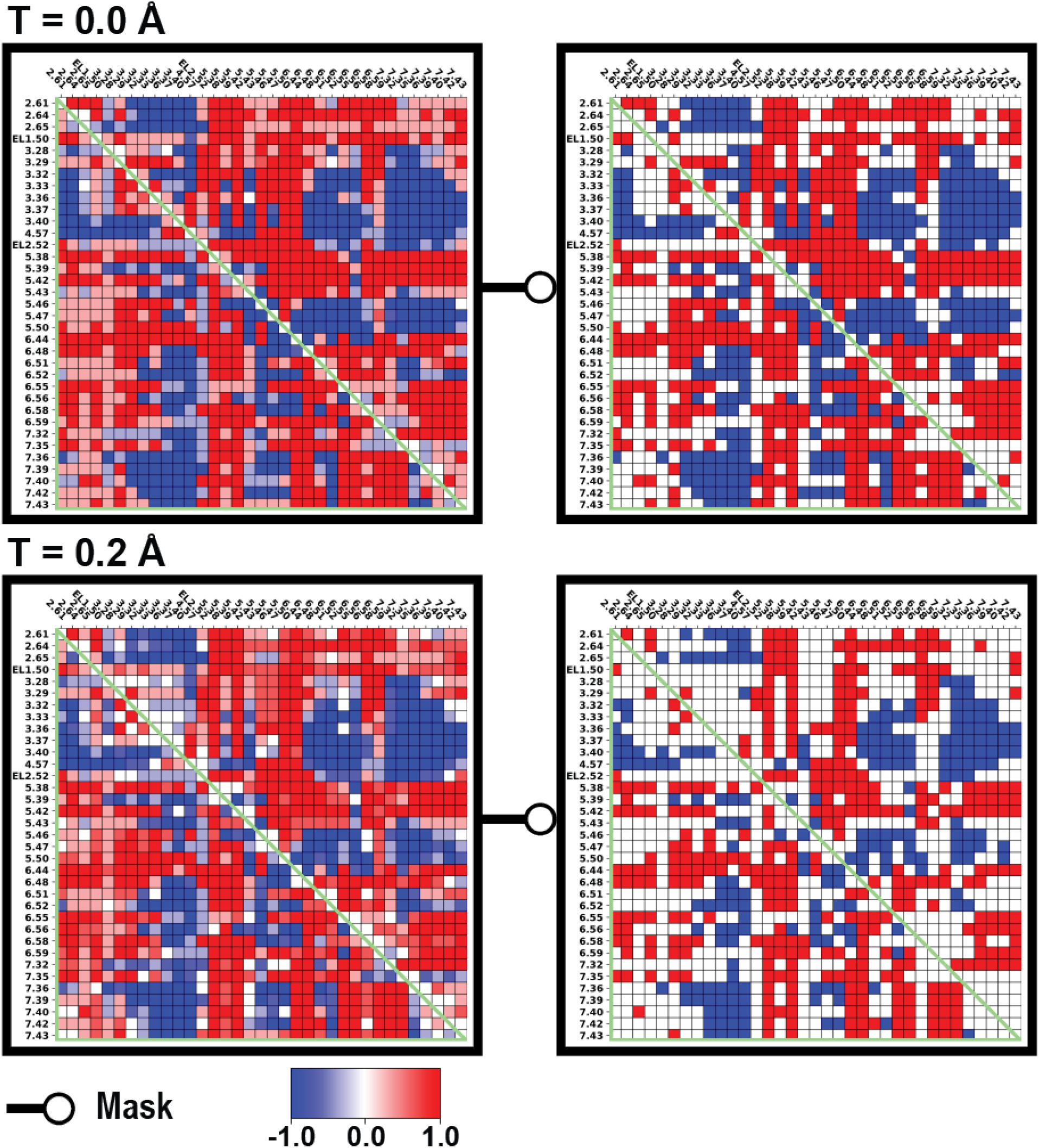
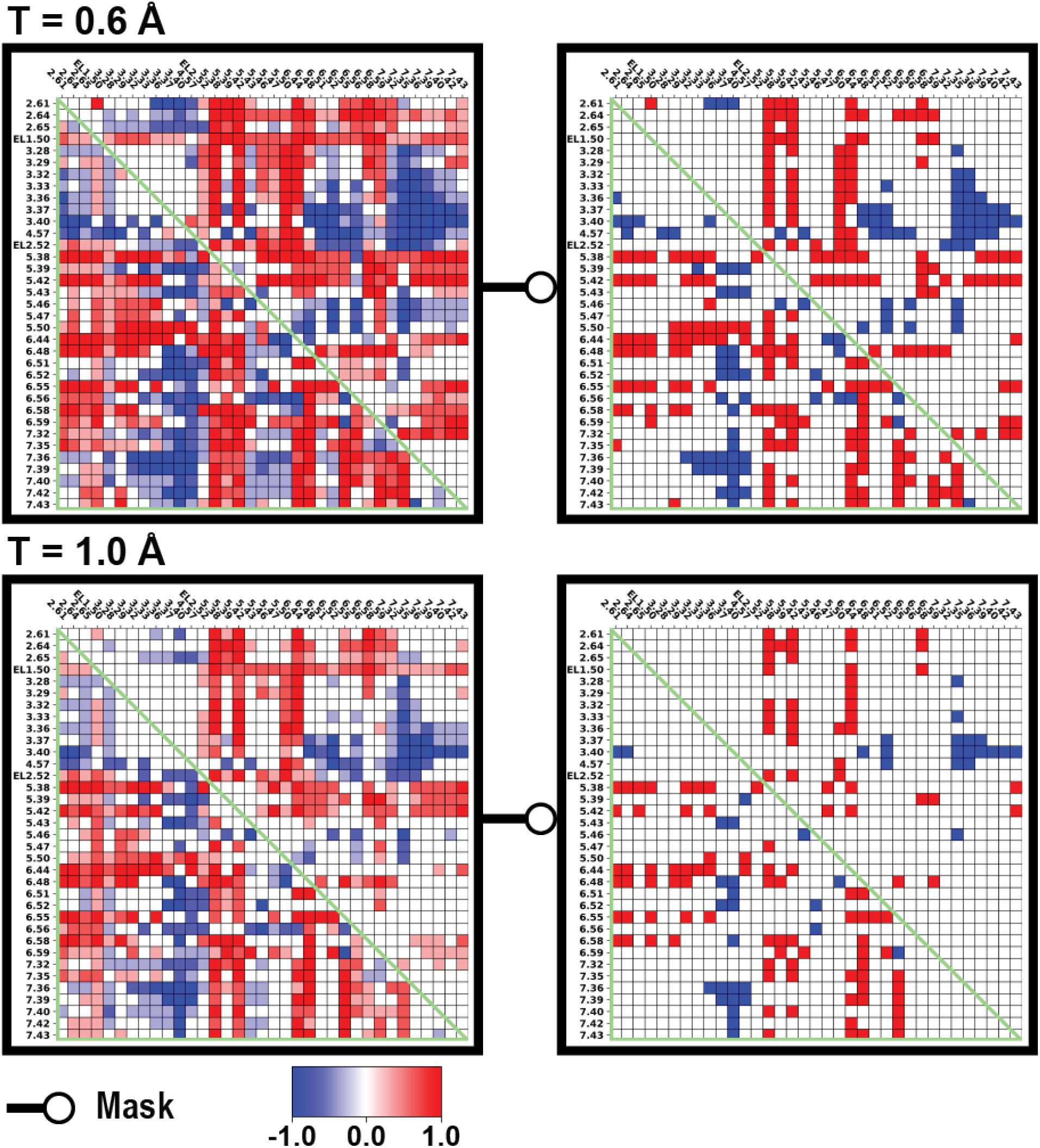
The binding site distance-difference category averages of the inactive and active D2R structure pairs. The heatmap shows the difference of pairwise distance among the binding site residues between the active and inactive structures (D2R active – D2R inactive) and categorized based on various DDthreshold (T). Red represents the active conformation have a larger distance than inactive conformation, and blue represents the inactive conformation have a larger distance than active conformation. If the category average of the difference is smaller or equal to 0.9, it is colored as white in the masked heatmaps.

**Figure S3.**
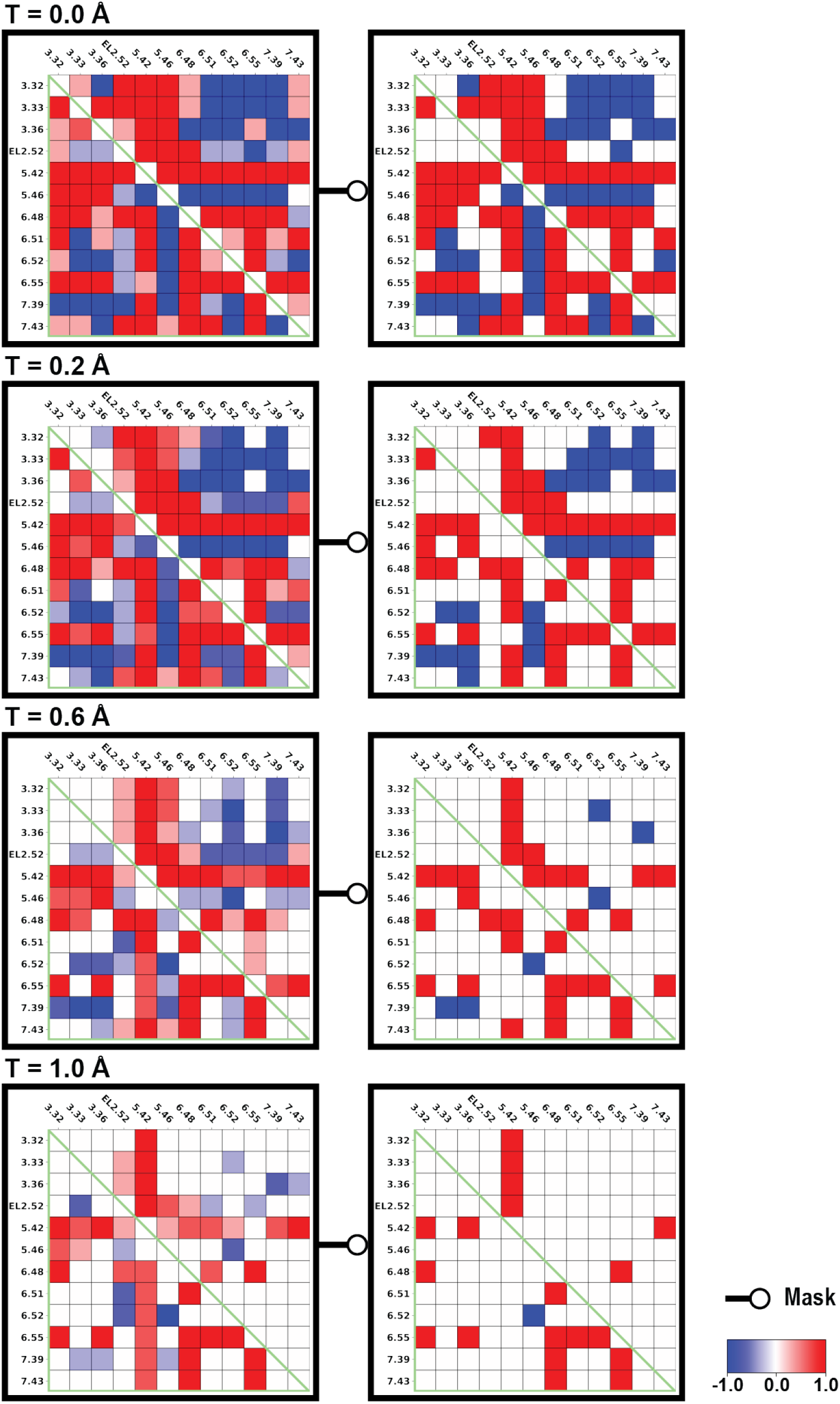
The OBS residue distance-difference category averages of the inactive and active D2R structure pairs. The heatmap shows the difference of pairwise distance among the OBS residues between the active and inactive structures (D2R active – D2R inactive) and categorized based on various DDthreshold (T). Red represents the active conformation have a larger distance than inactive conformation, and blue represents the inactive conformation have a larger distance than active conformation. If the category average of the difference is smaller or equal to 0.9, it is colored as white in the masked heatmaps.

**Figure S4.**
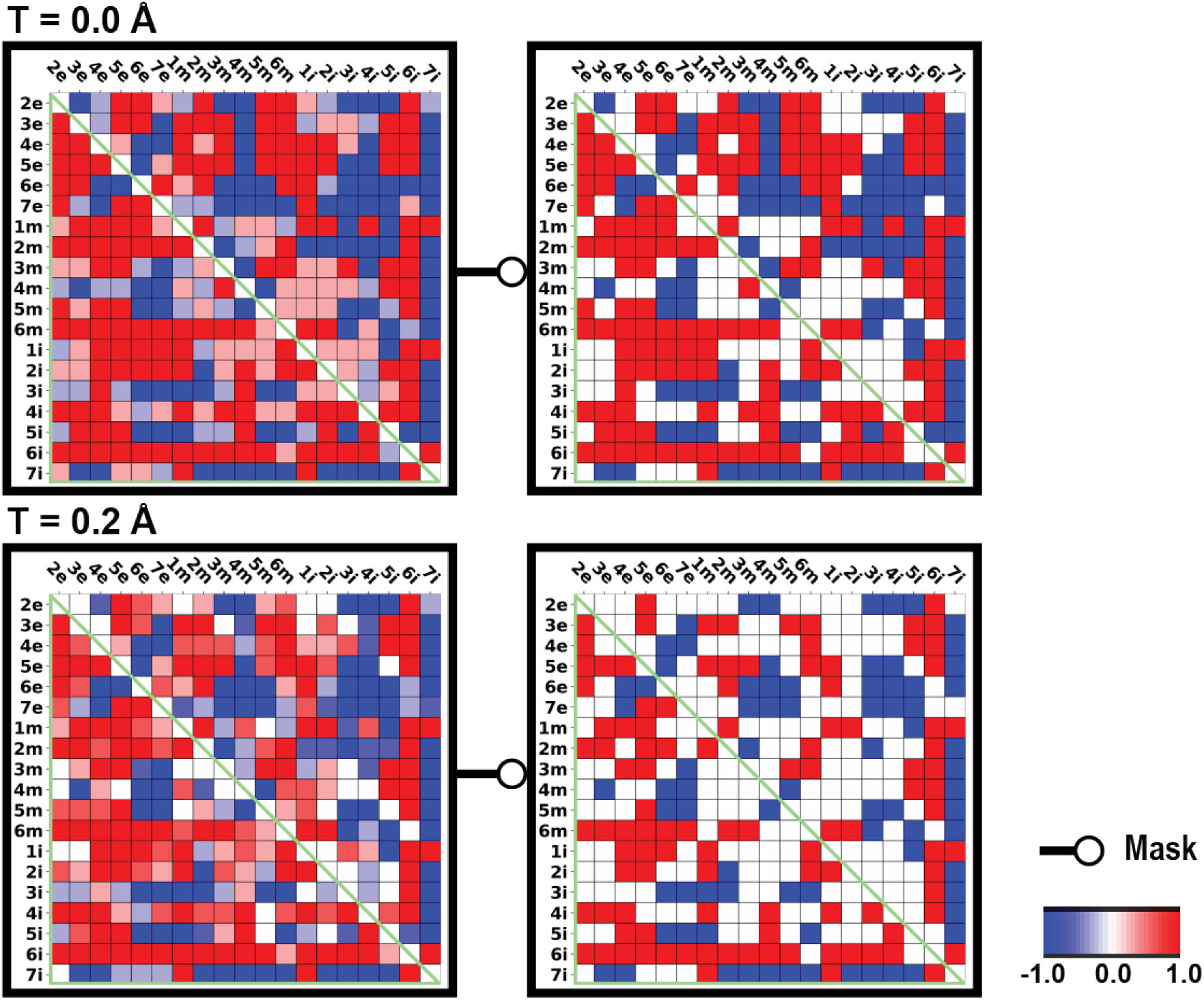
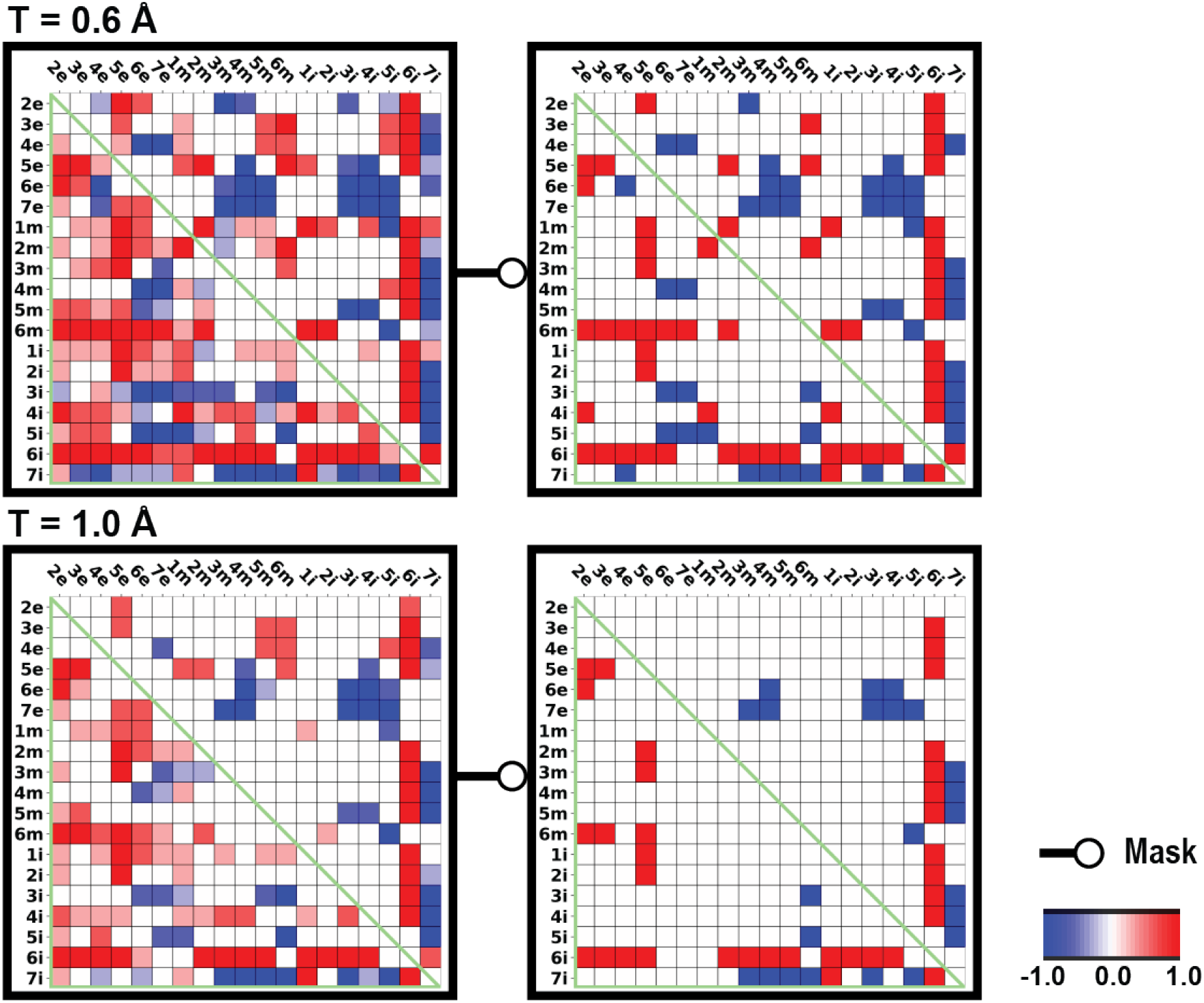
The subsegment distance-difference category averages of the inactive and active D2R structure pairs. The heatmap shows the difference of pairwise distance among the subsegments between the active and inactive structures (D2R active – D2R inactive) and categorized based on various DDthreshold (T). Red represents the active conformation have a larger distance than inactive conformation, and blue represents the inactive conformation have a larger distance than active conformation. If the category average of the difference is smaller or equal to 0.9, it is colored as white in the masked heatmaps.

**Figure S5.**
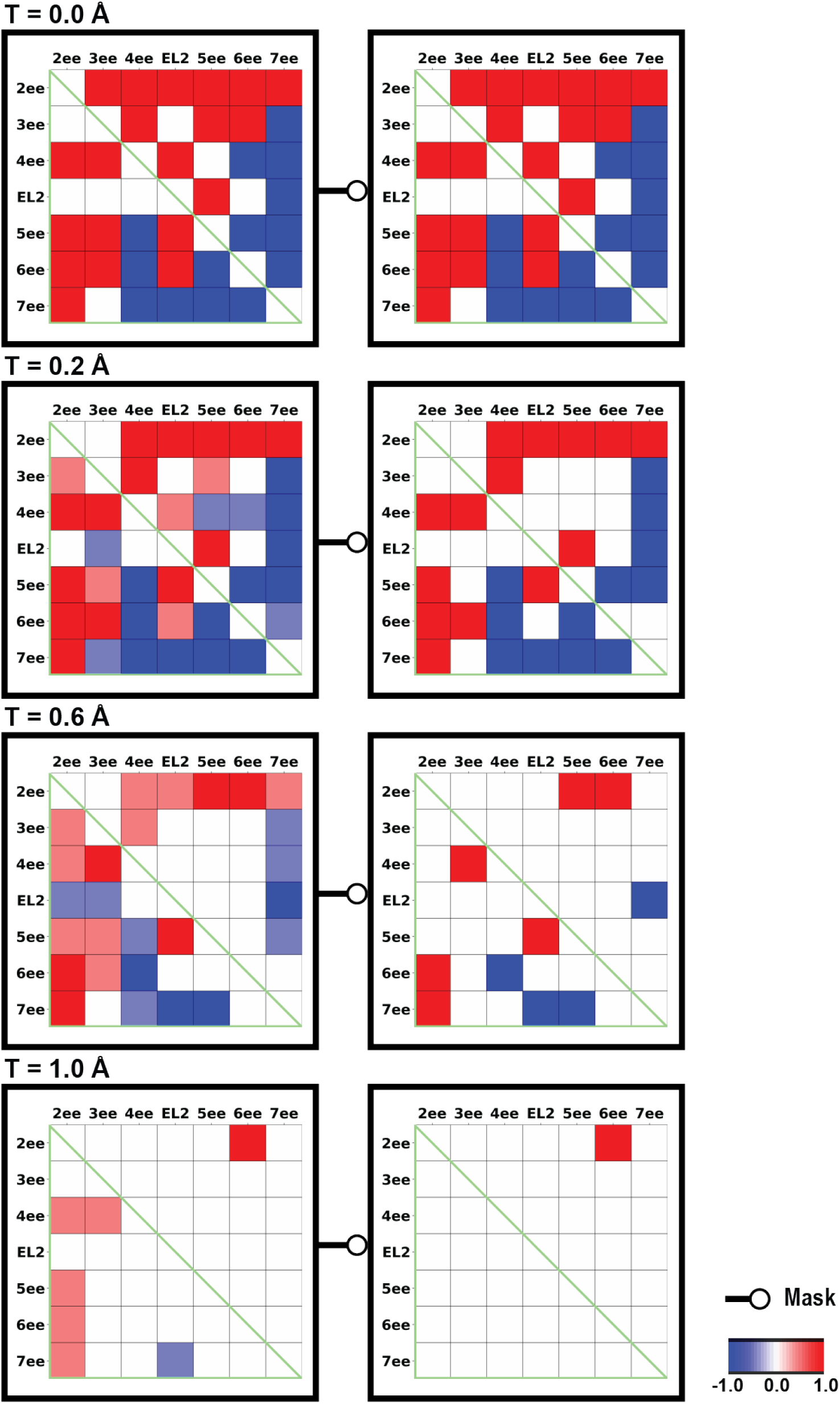
The extracellular-end distance differences of the inactive and active D3R structure pairs. The heatmap shows the difference of pairwise distance among the extracellular-end between the active and inactive structures (D3R active – D3R inactive) and categorized based on various DDthreshold (T). Red represents the active conformation have a larger distance than inactive conformation, and blue represents the inactive conformation have a larger distance than active conformation. If the category average of the difference is smaller or equal to 0.9, it is colored as white in the masked heatmaps.

**Figure S6.**
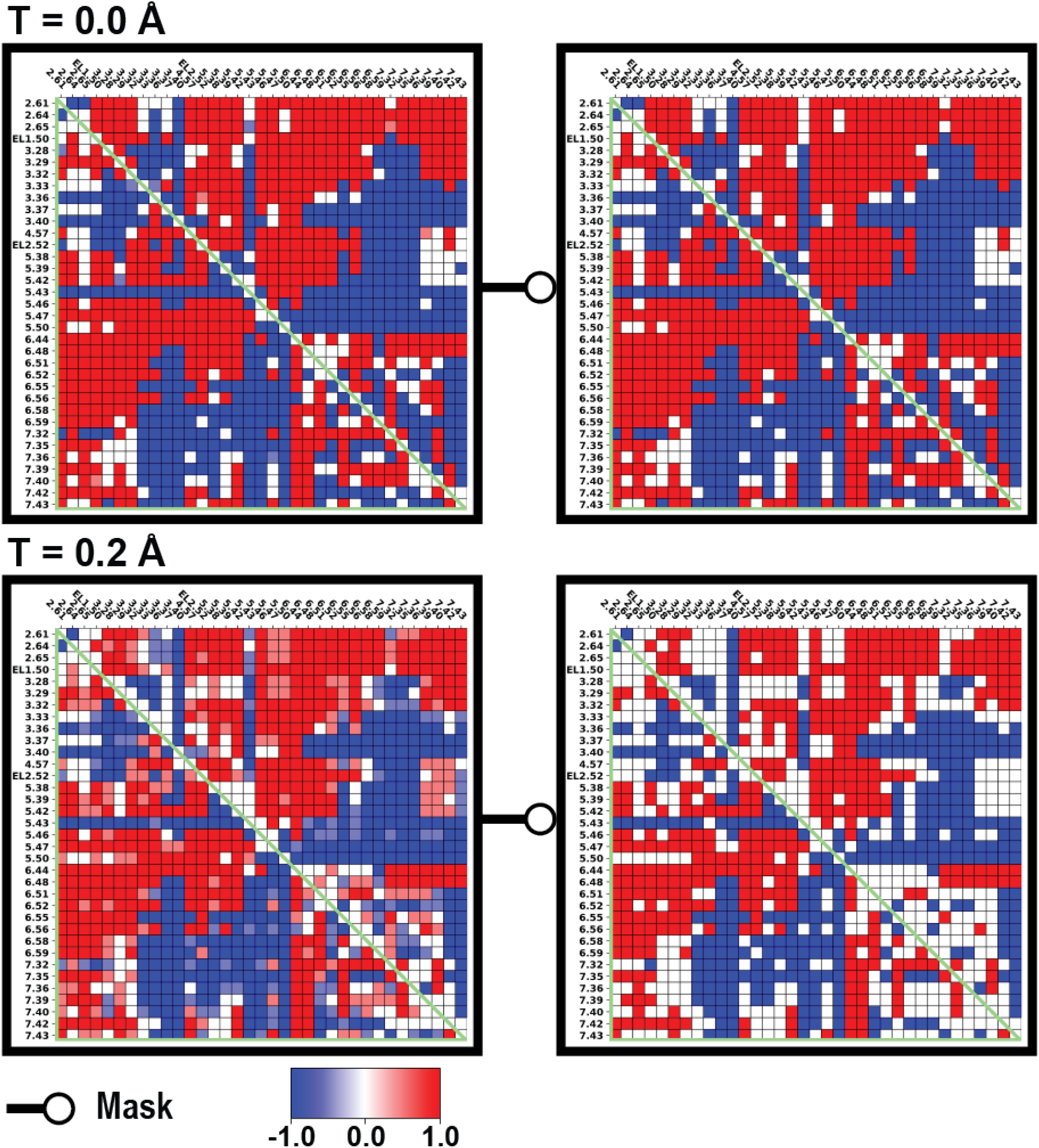
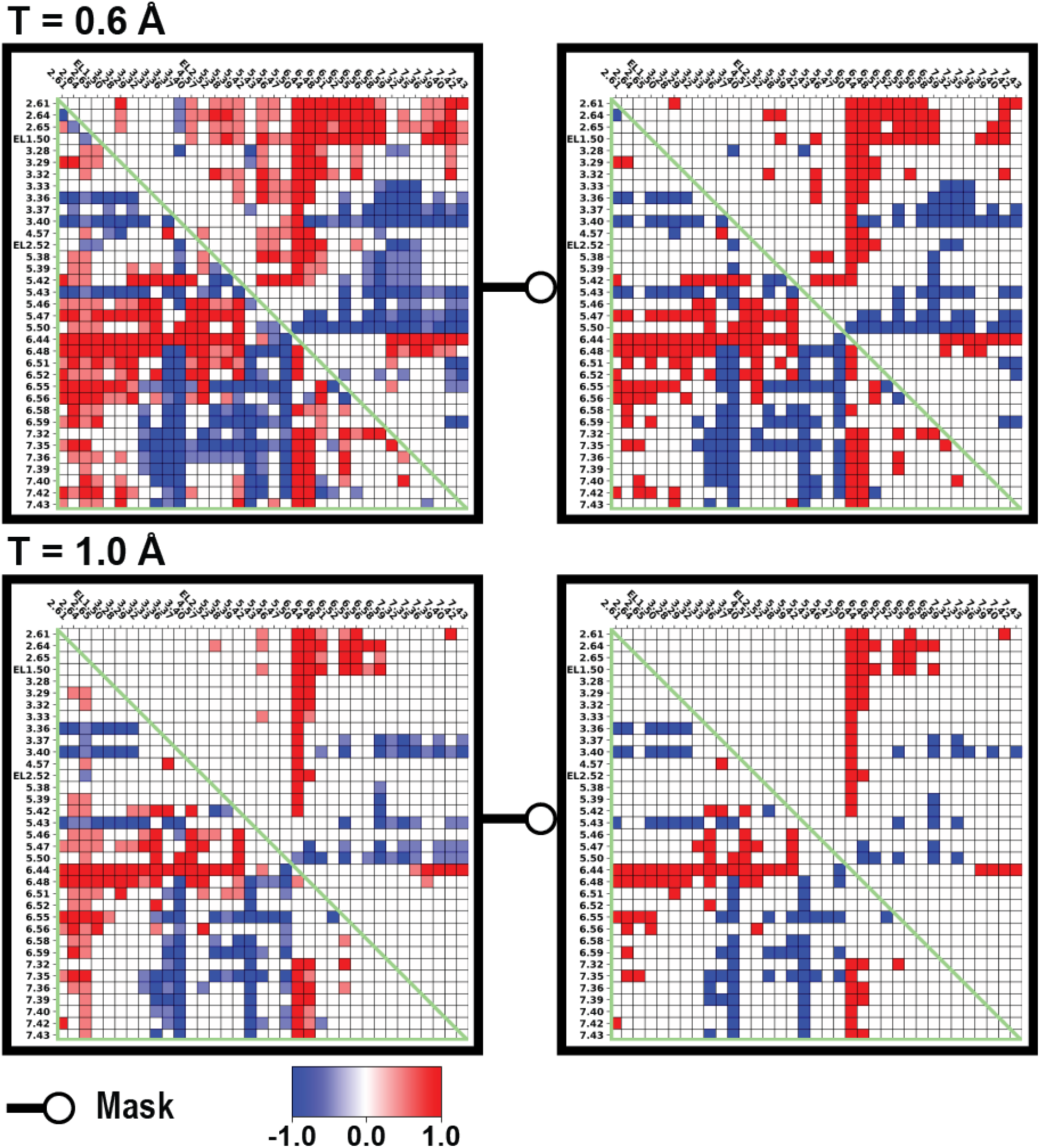
The binding site distance-difference category averages of the inactive and active D3R structure pairs. The heatmap shows the difference of pairwise distance among the binding site residues between the active and inactive structures (D3R active – D3R inactive) and categorized based on various DDthreshold (T). Red represents the active conformation have a larger distance than inactive conformation, and blue represents the inactive conformation have a larger distance than active conformation. If the category average of the difference is smaller or equal to 0.9, it is colored as white in the masked heatmaps.

**Figure S7.**
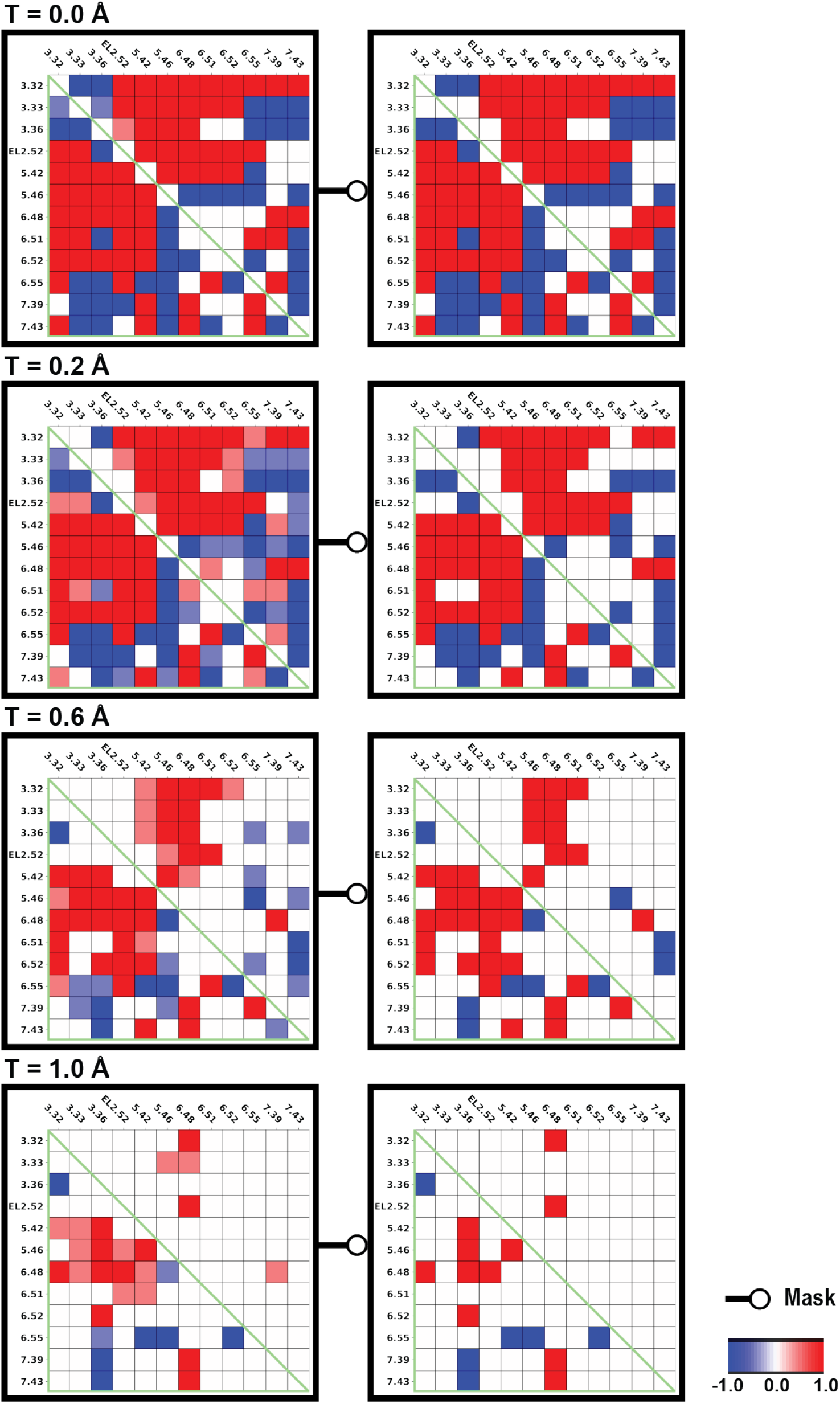
The OBS residue distance-difference category averages of the inactive and active D3R structure pairs. The heatmap shows the difference of pairwise distance among the OBS residues between the active and inactive structures (D3R active – D3R inactive) and categorized based on various DDthreshold (T). Red represents the active conformation have a larger distance than inactive conformation, and blue represents the inactive conformation have a larger distance than active conformation. If the category average of the difference is smaller or equal to 0.9, it is colored as white in the masked heatmaps.

**Figure S8.**
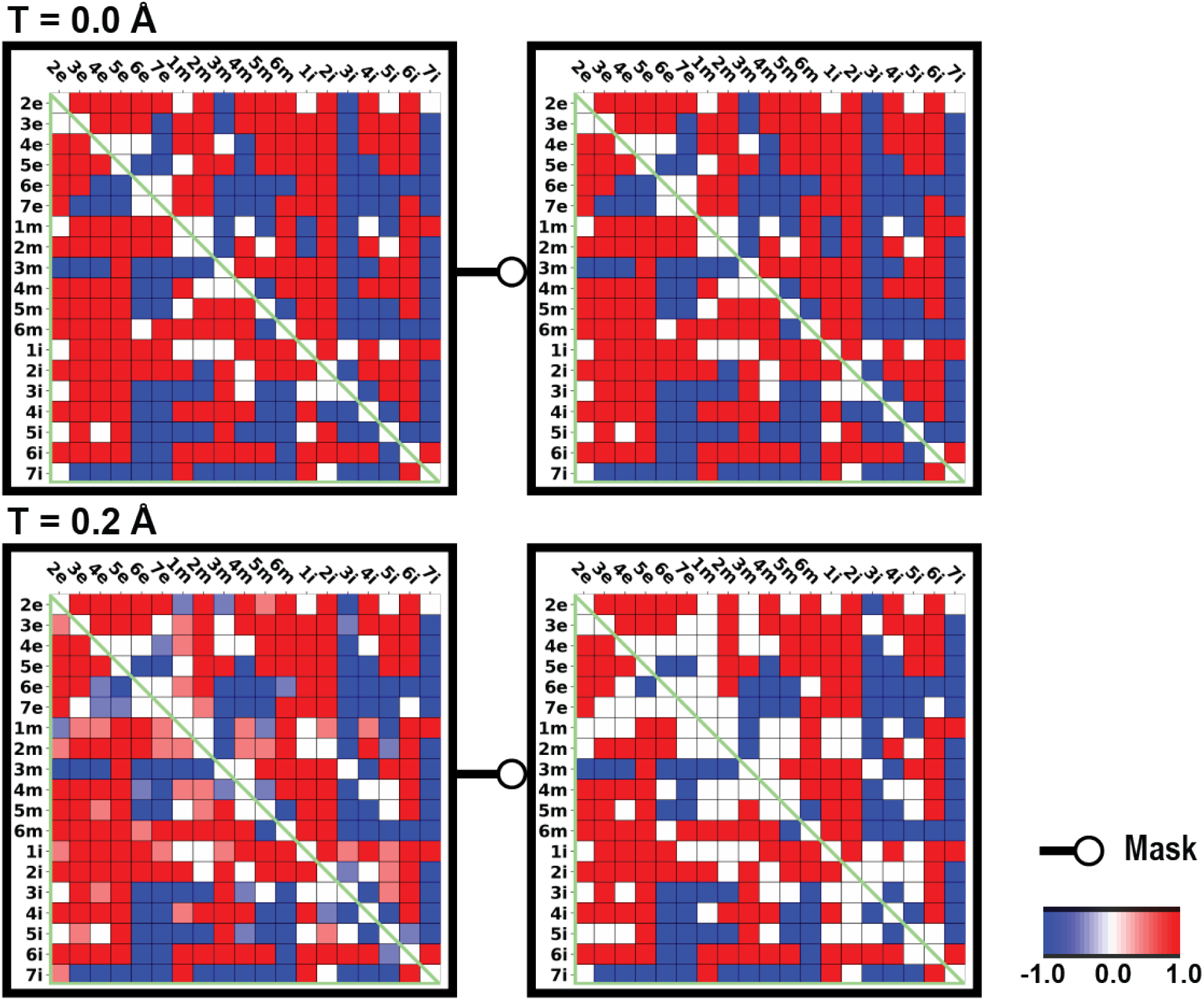
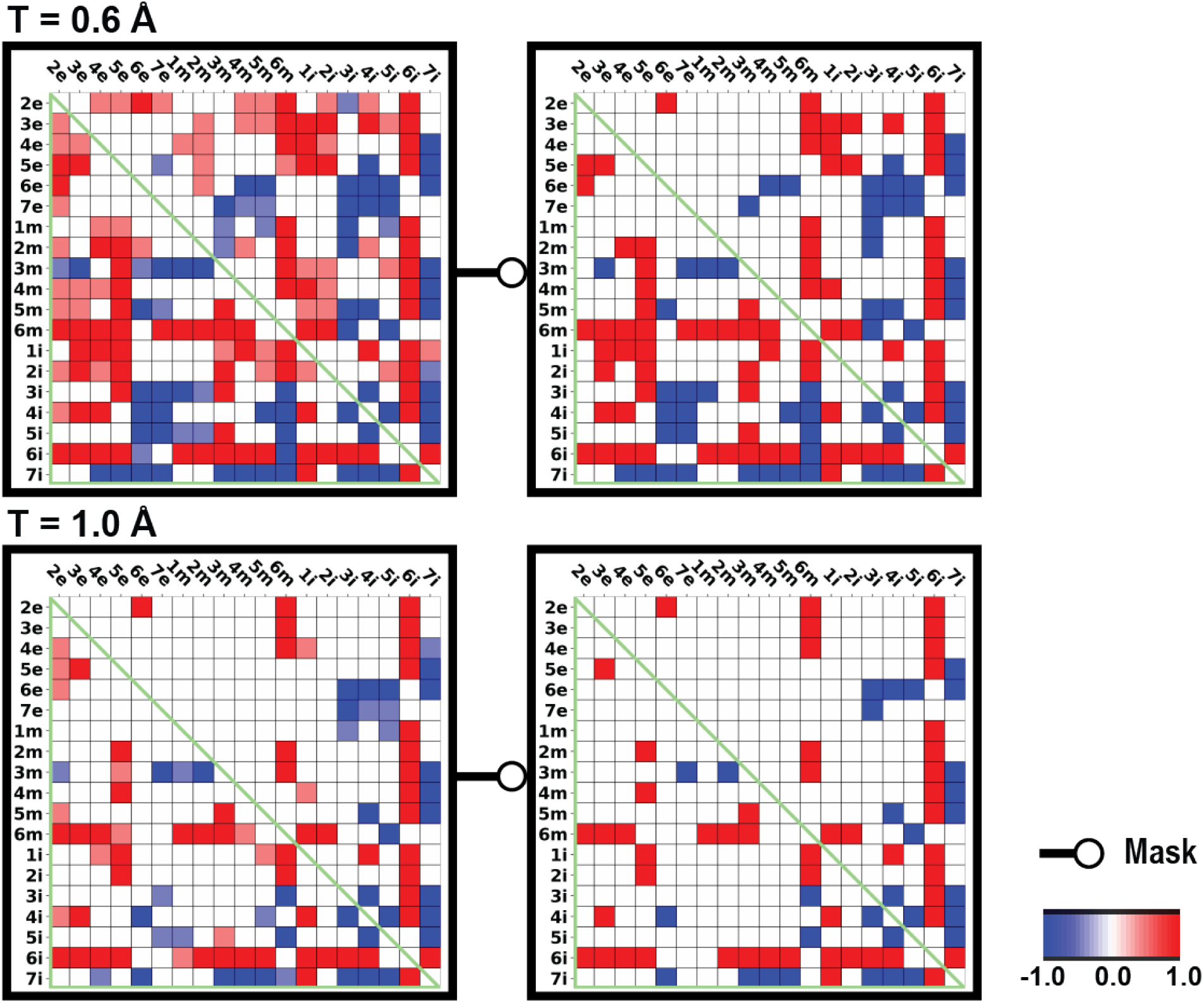
The subsegment distance-difference category averages of the inactive and active D3R structure pairs. The heatmap shows the difference of pairwise distance among the subsegments between the active and inactive structures (D3R active – D3R inactive) and categorized based on various DDthreshold (T). Red represents the active conformation have a larger distance than inactive conformation, and blue represents the inactive conformation have a larger distance than active conformation. If the category average of the difference is smaller or equal to 0.9, it is colored as white in the masked heatmaps.

**Figure S9.**
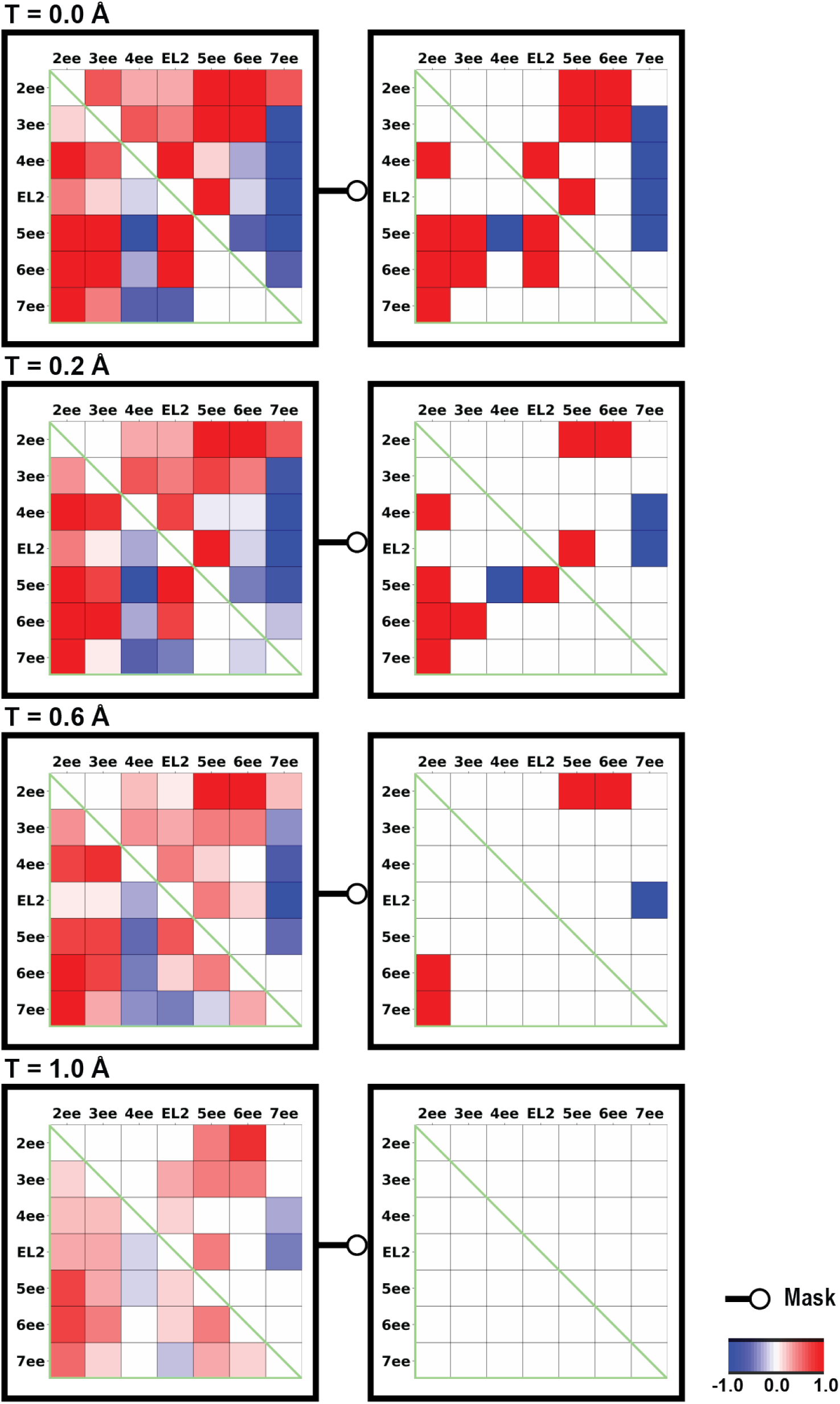
The extracellular-end distance differences of the inactive and active D2R and D3R structure pairs. The heatmap shows the difference of pairwise distance among the extracellular-end between the active and inactive structures (active – inactive) and categorized based on various DDthreshold (T). Red represents the active conformation have a larger distance than inactive conformation, and blue represents the inactive conformation have a larger distance than active conformation. If the category average of the difference is smaller or equal to 0.9, it is colored as white in the masked heatmaps.

**Figure S10.**
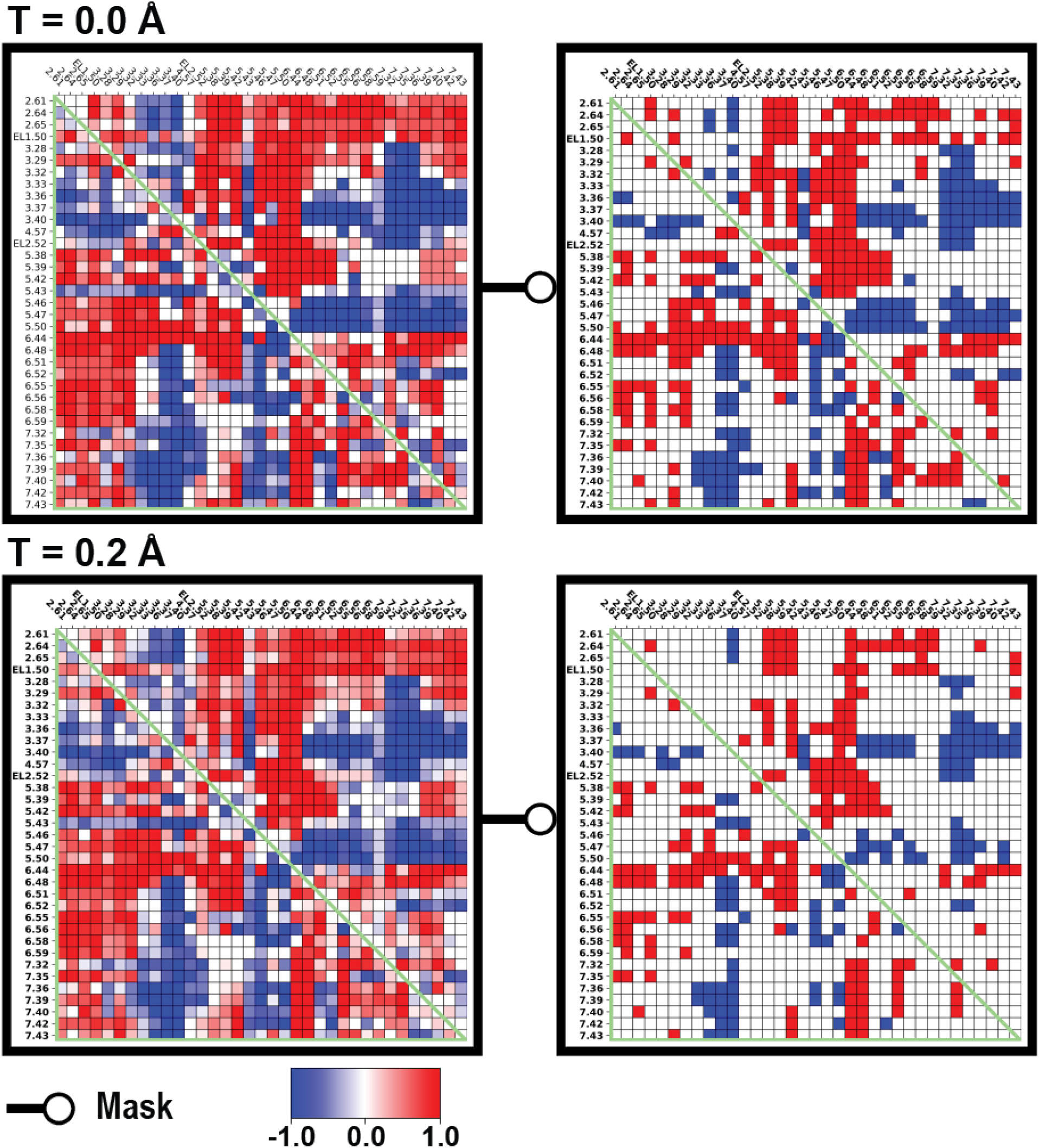
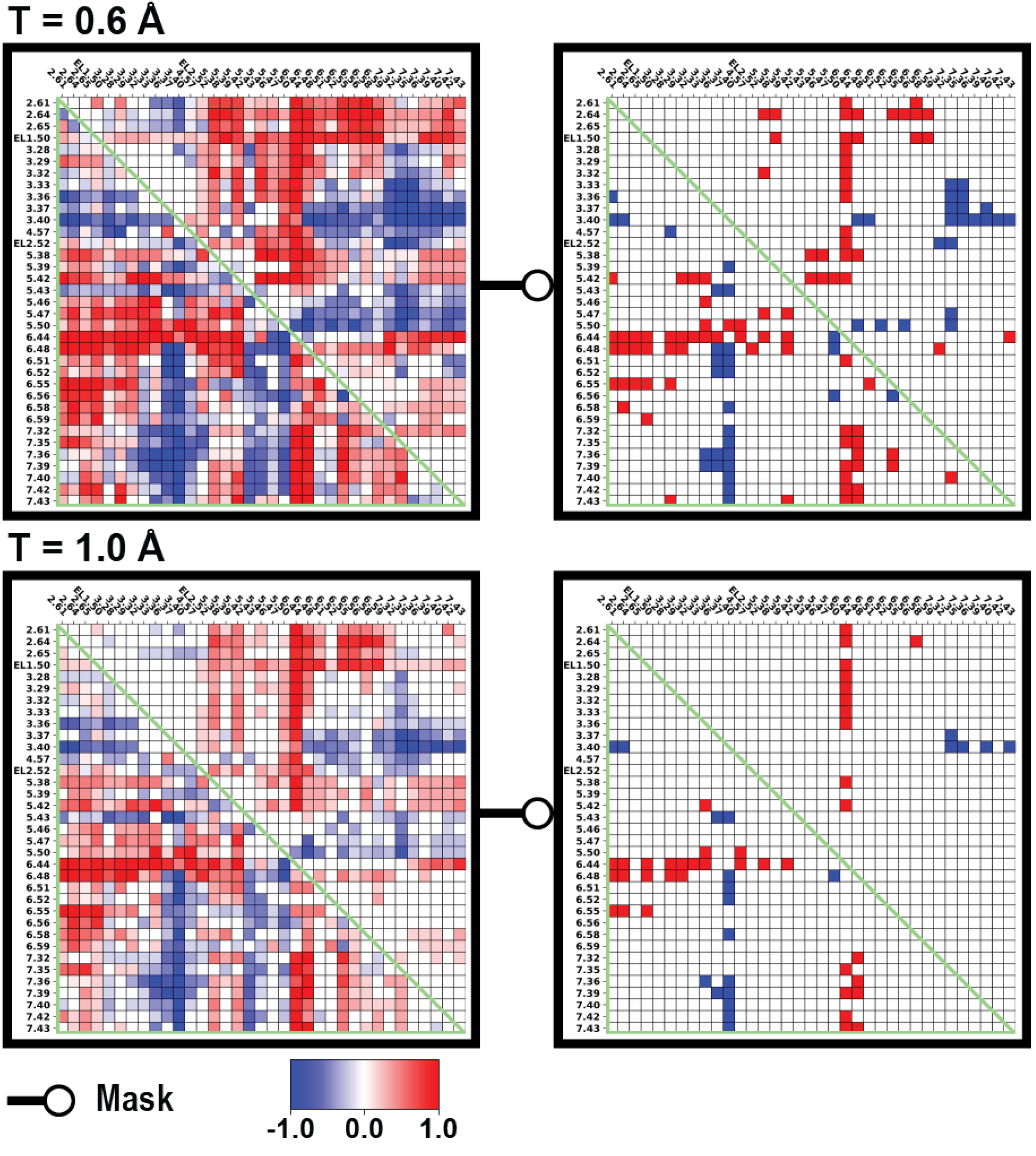
The binding site distance-difference category averages of the inactive and active D2R and D3R structure pairs. The heatmap shows the difference of pairwise distance among the binding site residues between the active and inactive structures (active –inactive) and categorized based on various DDthreshold (T). Red represents the active conformation have a larger distance than inactive conformation, and blue represents the inactive conformation have a larger distance than active conformation. If the category average of the difference is smaller or equal to 0.9, it is colored as white in the masked heatmaps.

**Figure S11.**
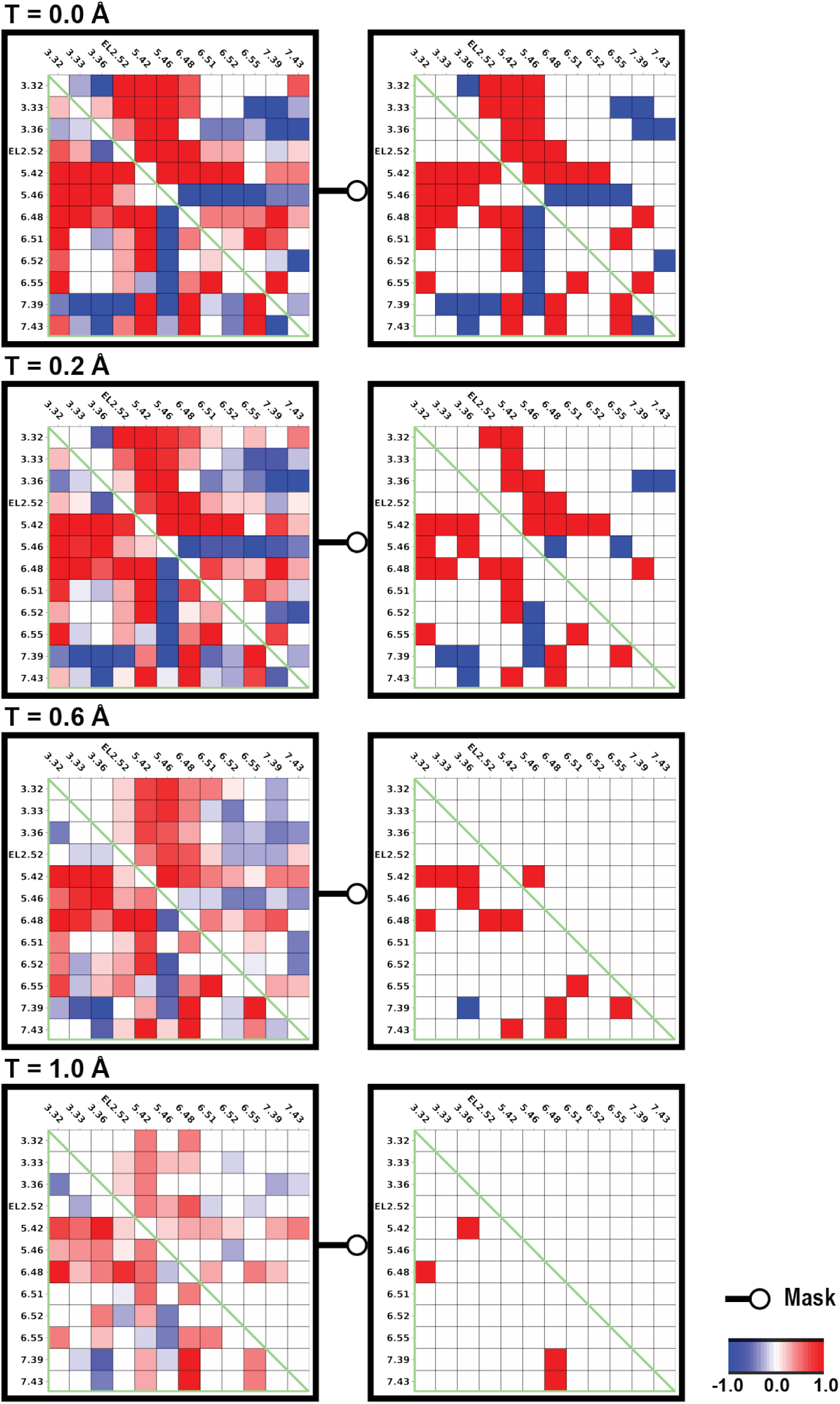
The OBS residue distance-difference category averages of the inactive and active D2R and D3R structure pairs. The heatmap shows the difference of pairwise distance among the OBS residues between the active and inactive structures (active – inactive) and categorized based on various DDthreshold (T). Red represents the active conformation have a larger distance than inactive conformation, and blue represents the inactive conformation have a larger distance than active conformation. If the category average of the difference is smaller or equal to 0.9, it is colored as white in the masked heatmaps.

**Figure S12.**
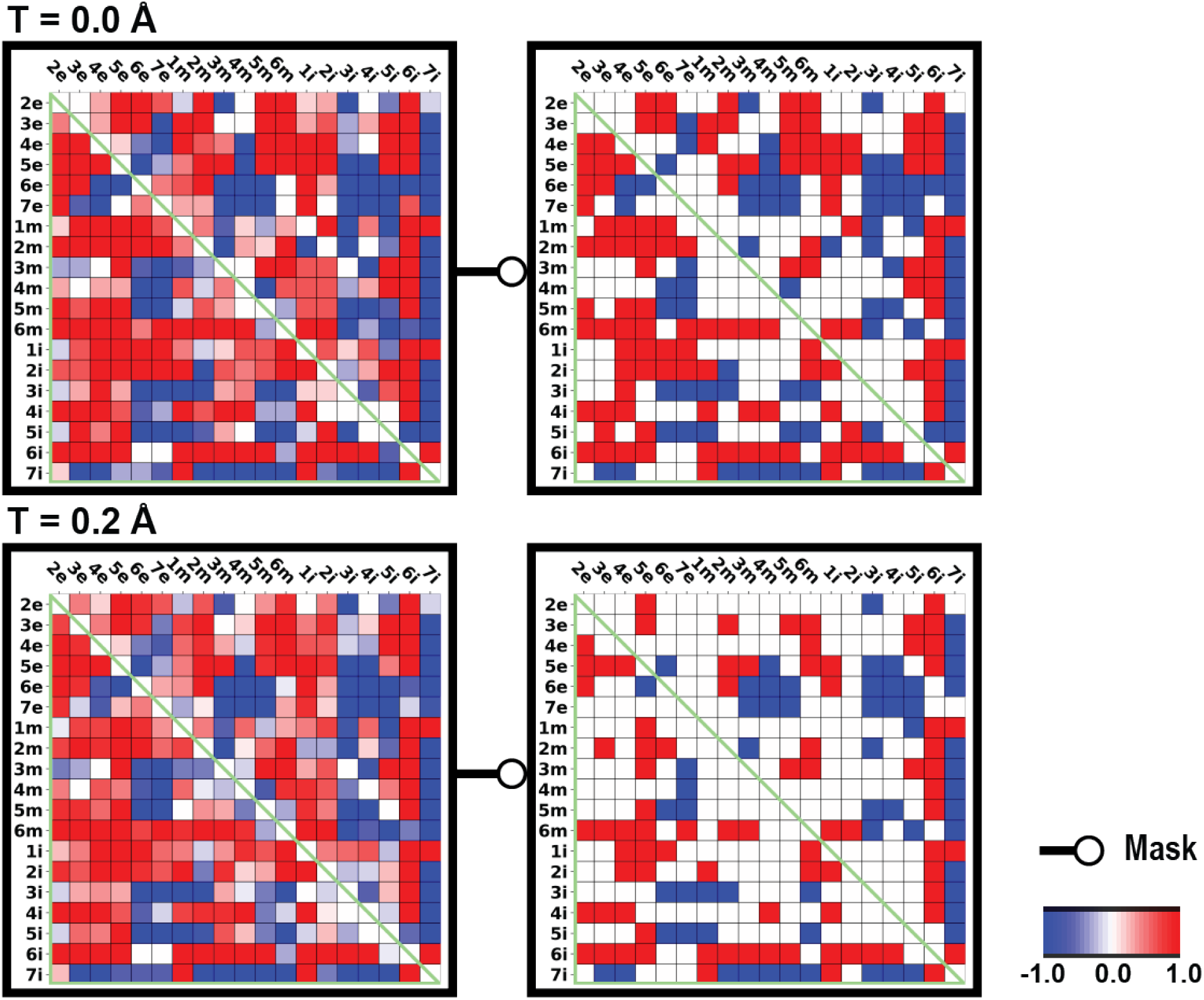
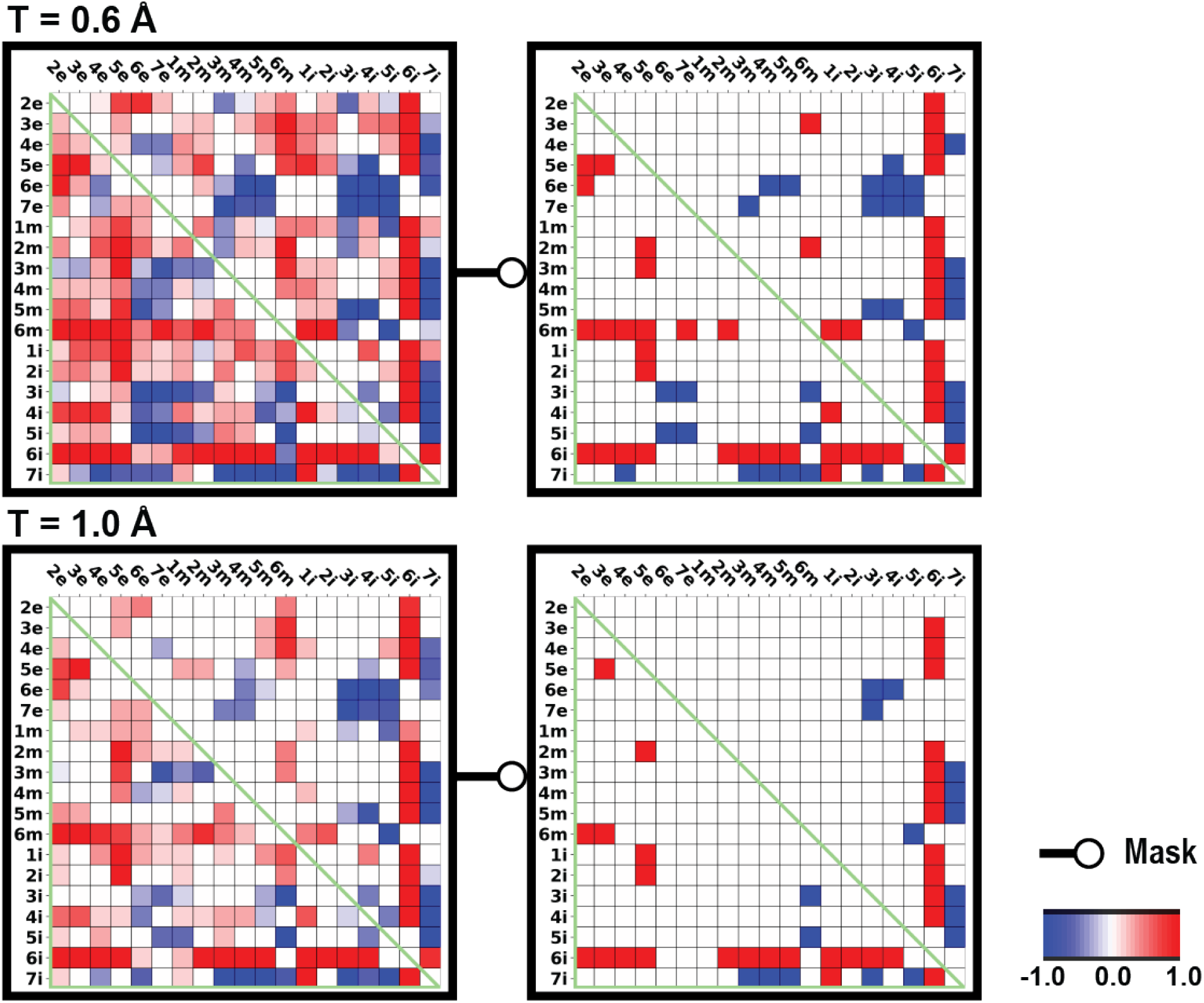
The subsegment distance-difference category averages of the inactive and active D2R and D3R structure pairs. The heatmap shows the difference of pairwise distance among the subsegments between the active and inactive structures (active – inactive) and categorized based on various DDthreshold (T). Red represents the active conformation have a larger distance than inactive conformation, and blue represents the inactive conformation have a larger distance than active conformation. If the category average of the difference is smaller or equal to 0.9, it is colored as white in the masked heatmaps.

**Figure S13.**
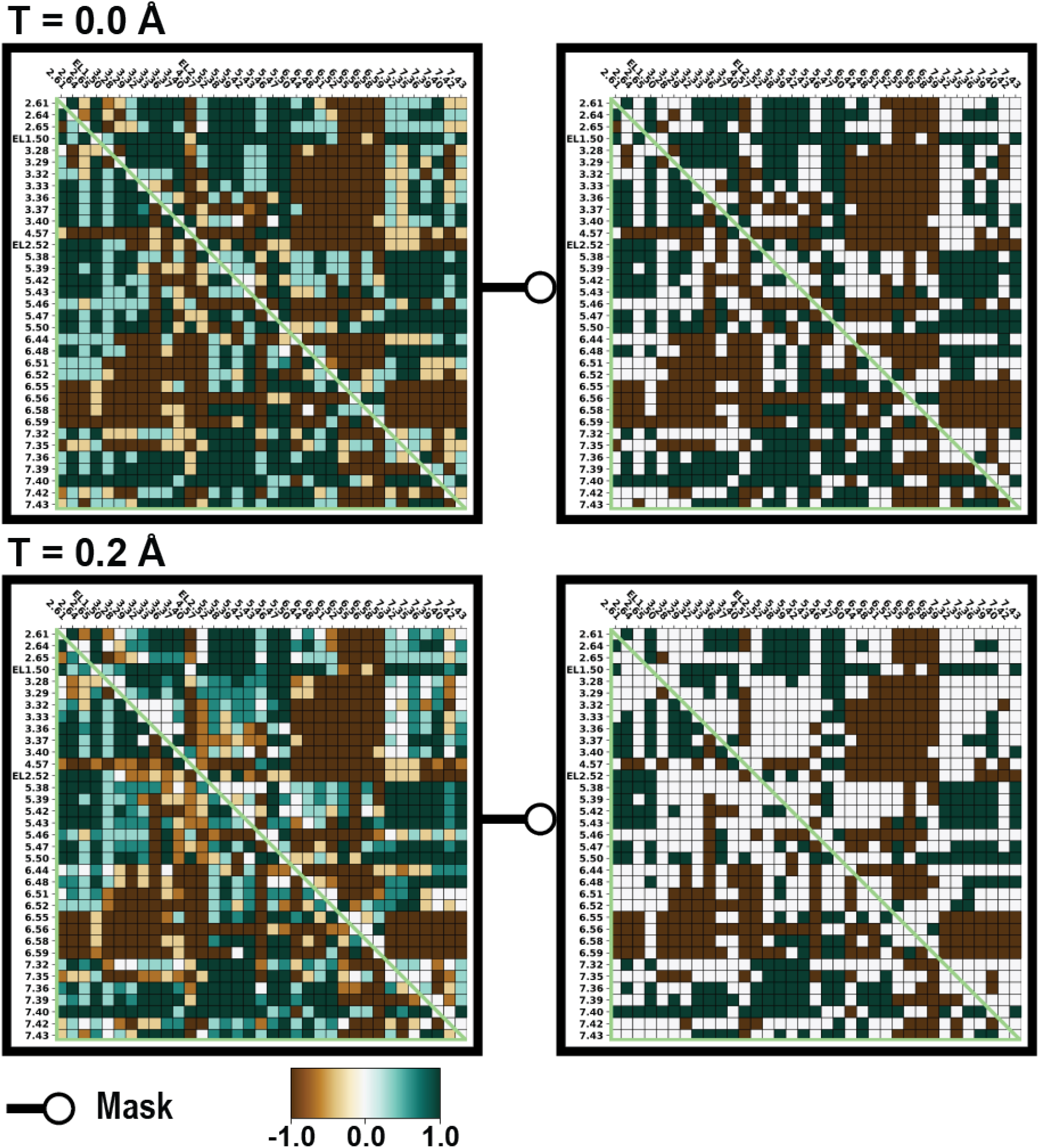
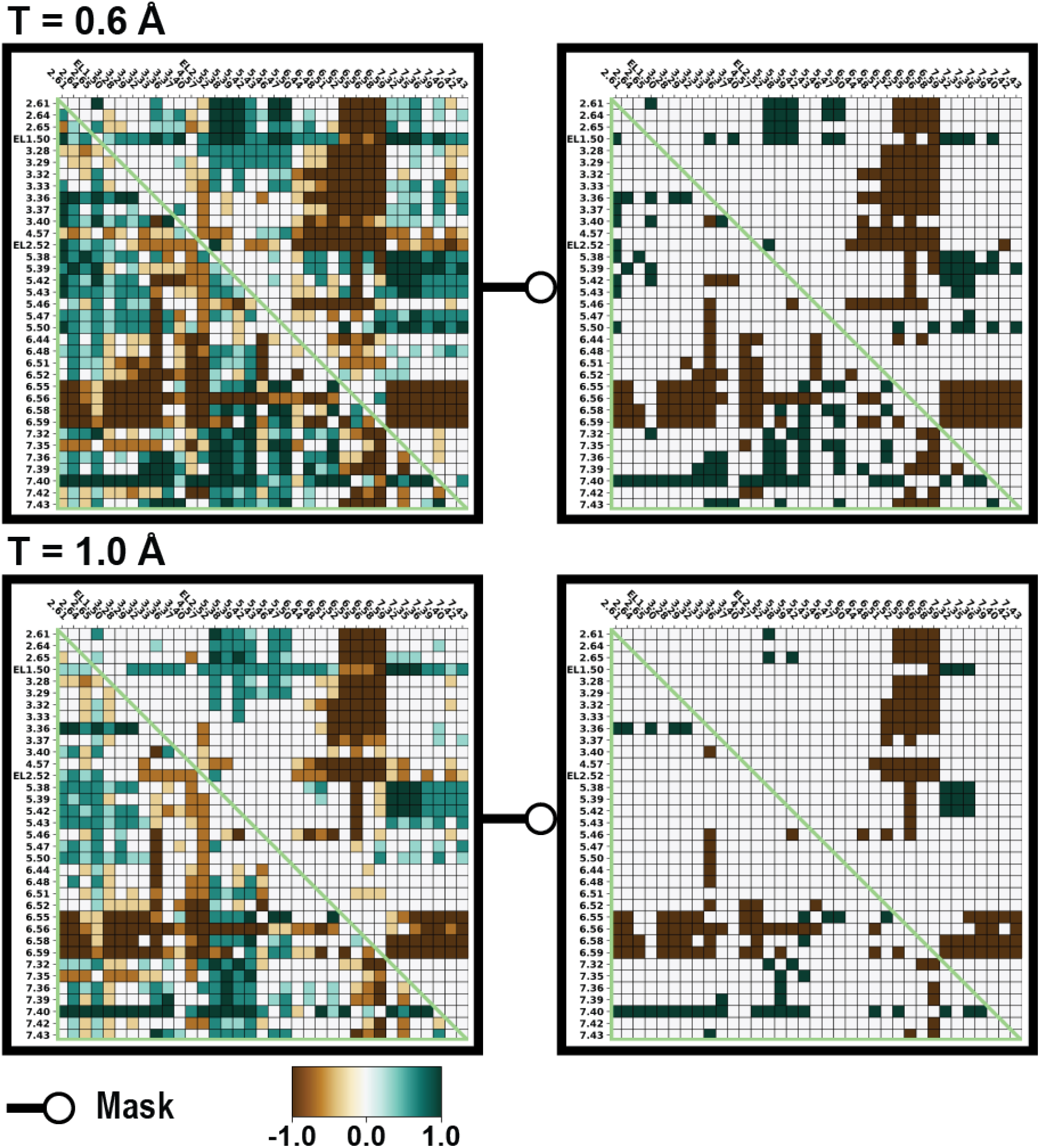
The binding site distance-difference category averages of the comparison between the inactive D3R and D2R structures. The heatmap shows the difference of pairwise distance among the binding site residues between the D3R and D2R using their inactive structures (D3R – D2R) and categorized based on various DDthreshold (T). Green represents the D3R conformation have a larger distance than D2R conformation, and brown represents the D2R conformation have a larger distance than D3R conformation. If the category average of the difference is smaller or equal to 0.9, it is colored as white in the masked heatmaps.

**Figure S14.**
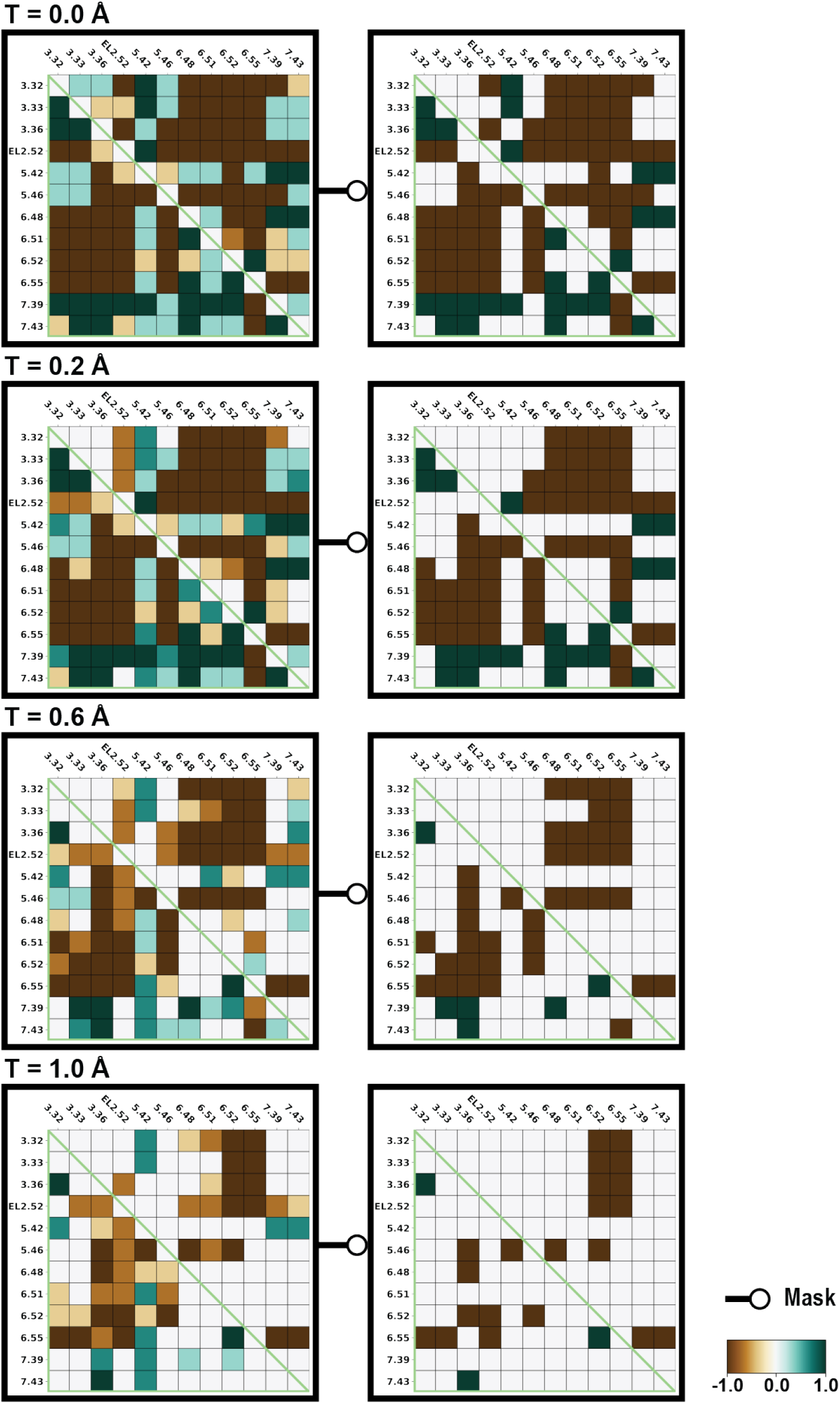
The OBS residue distance-difference category averages of the comparison between the inactive D3R and D2R structures. The heatmap shows the difference of pairwise distance among the OBS residues between the D3R and D2R using their inactive structures (D3R – D2R) and categorized based on various DDthreshold (T). Green represents the D3R conformation have a larger distance than D2R conformation, and brown represents the D2R conformation have a larger distance than D3R conformation. If the category average of the difference is smaller or equal to 0.9, it is colored as white in the masked heatmaps.

**Figure S15.**
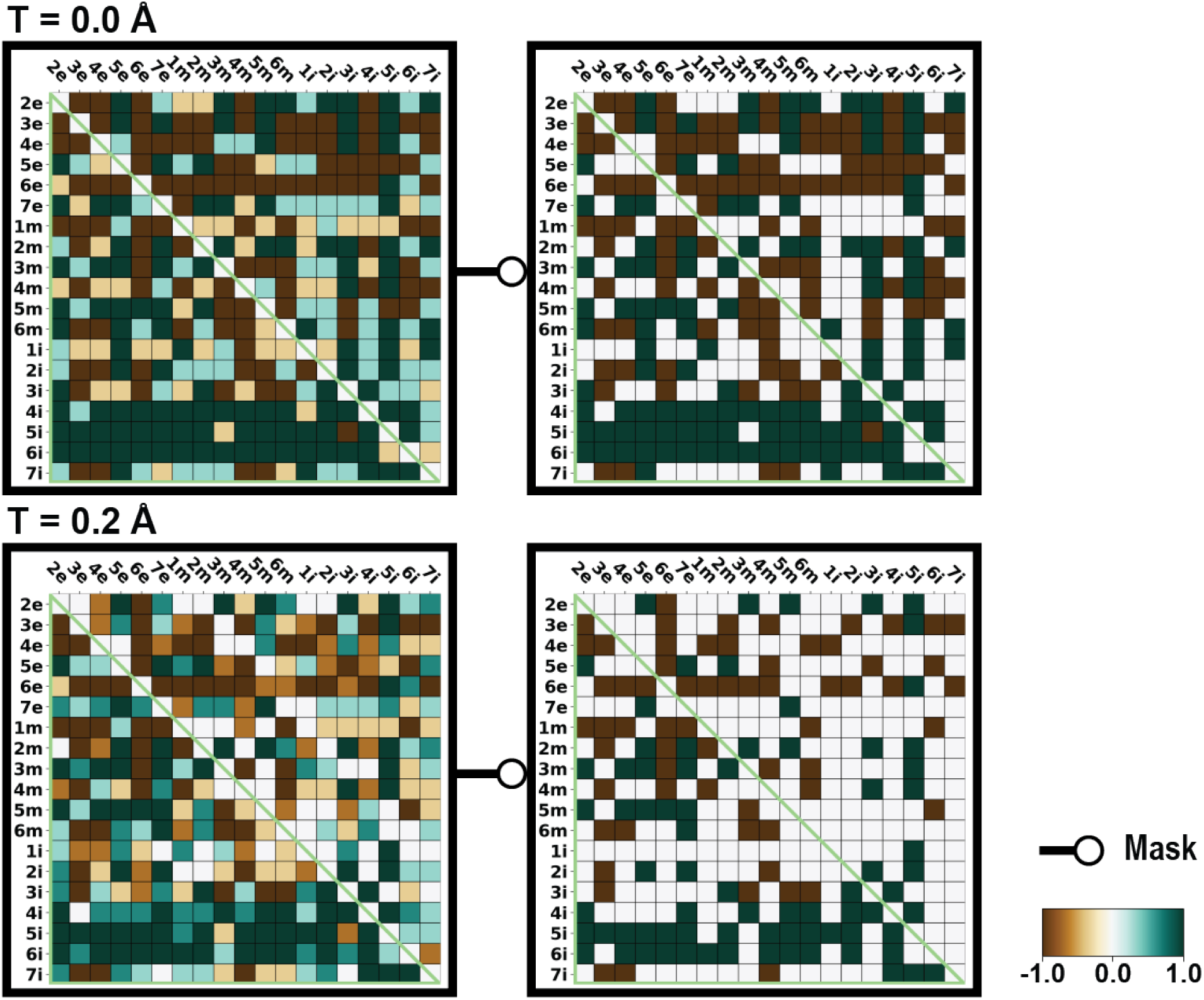
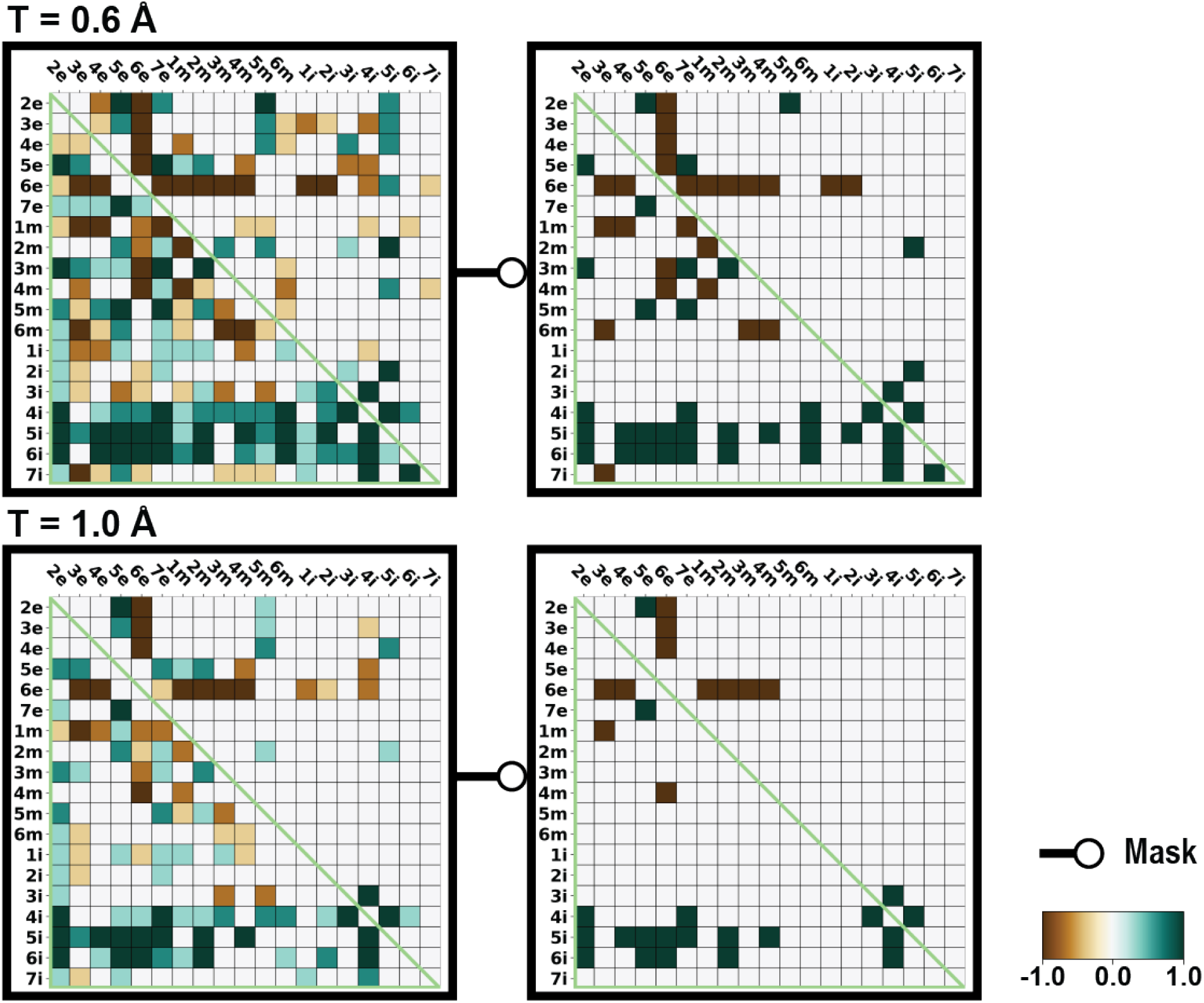
The subsegment distance-difference category averages of the comparison between the inactive D3R and D2R structures. The heatmap shows the difference of pairwise distance among the subsegments between the D3R and D2R using their inactive structures (D3R – D2R) and categorized based on various DDthreshold (T). Green represents the D3R conformation have a larger distance than D2R conformation, and brown represents the D2R conformation have a larger distance than D3R conformation. If the category average of the difference is smaller or equal to 0.9, it is colored as white in the masked heatmaps.

**Figure S16.**
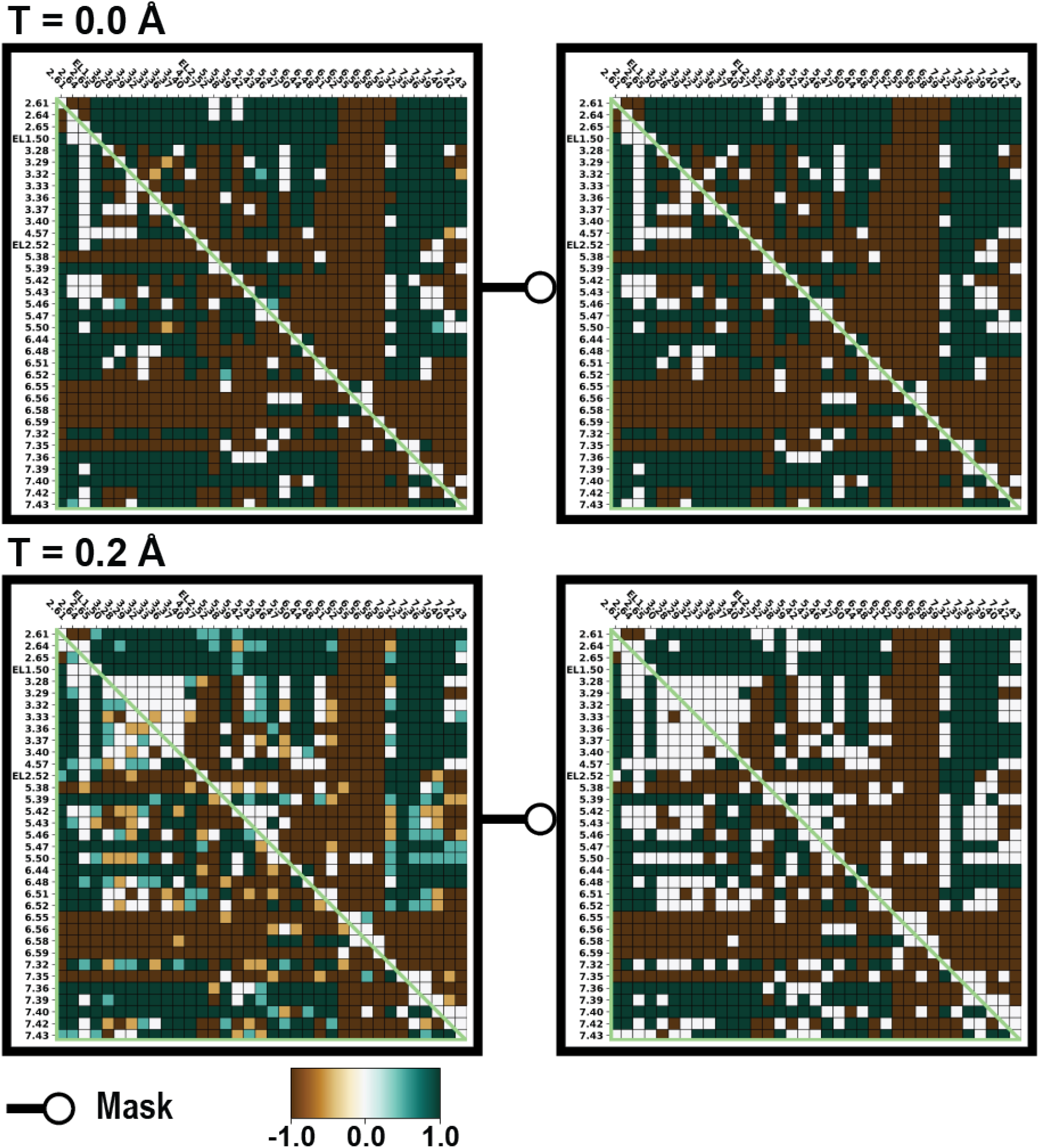
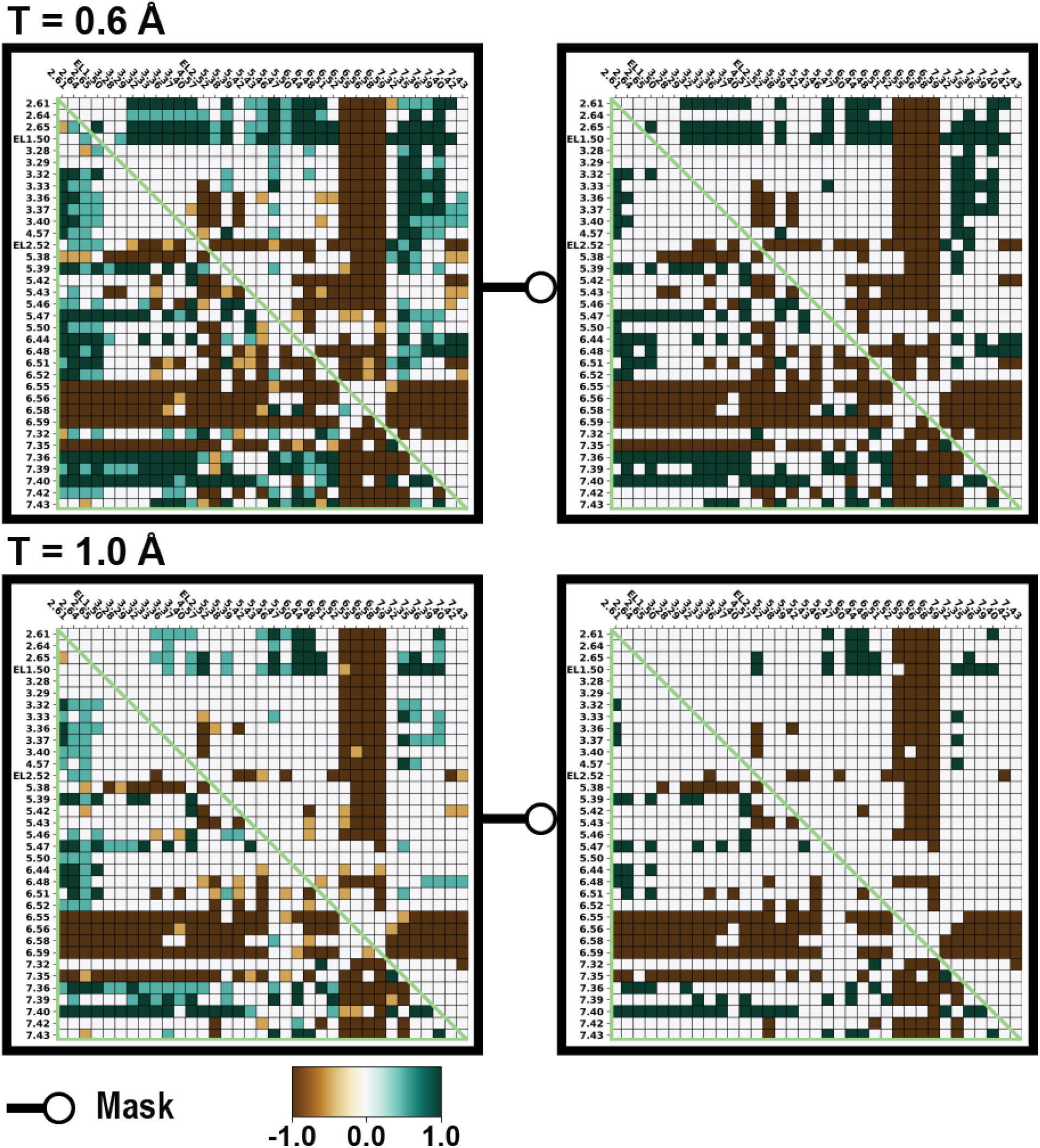
The binding site distance-difference category averages of the comparison between the active D3R and D2R structures. The heatmap shows the difference of pairwise distance among the binding site residues between the D3R and D2R using their active structures (D3R – D2R) and categorized based on various DDthreshold (T). Green represents the D3R conformation have a larger distance than D2R conformation, and brown represents the D2R conformation have a larger distance than D3R conformation. If the category average of the difference is smaller or equal to 0.9, it is colored as white in the masked heatmaps.

**Figure S17.**
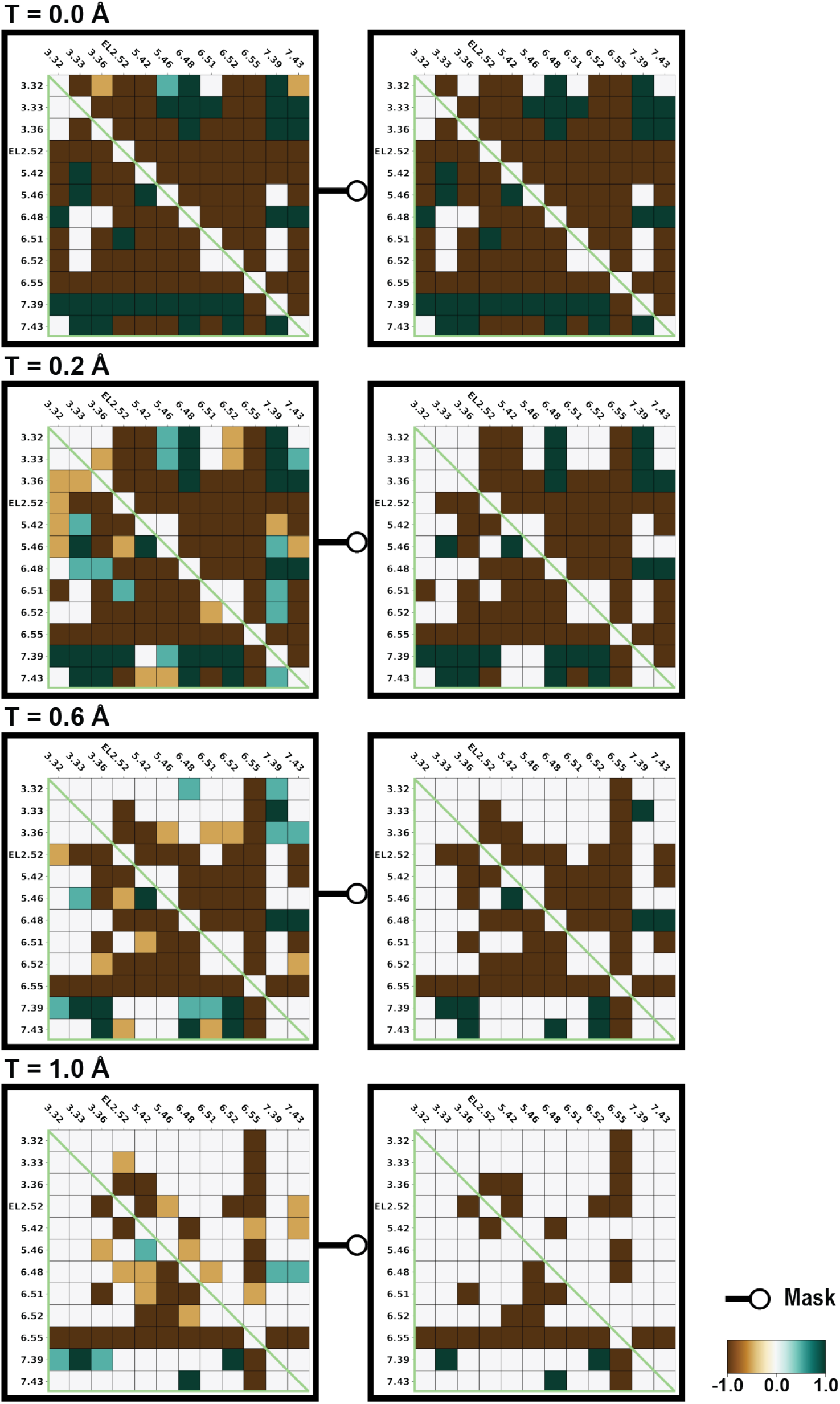
The OBS residue distance-difference category averages of the comparison between the active D3R and D2R structures. The heatmap shows the difference of pairwise distance among the OBS residues between the D3R and D2R using their active structures (D3R – D2R) and categorized based on various DDthreshold (T). Green represents the D3R conformation have a larger distance than D2R conformation, and brown represents the D2R conformation have a larger distance than D3R conformation. If the category average of the difference is smaller or equal to 0.9, it is colored as white in the masked heatmaps.

**Figure S18.**
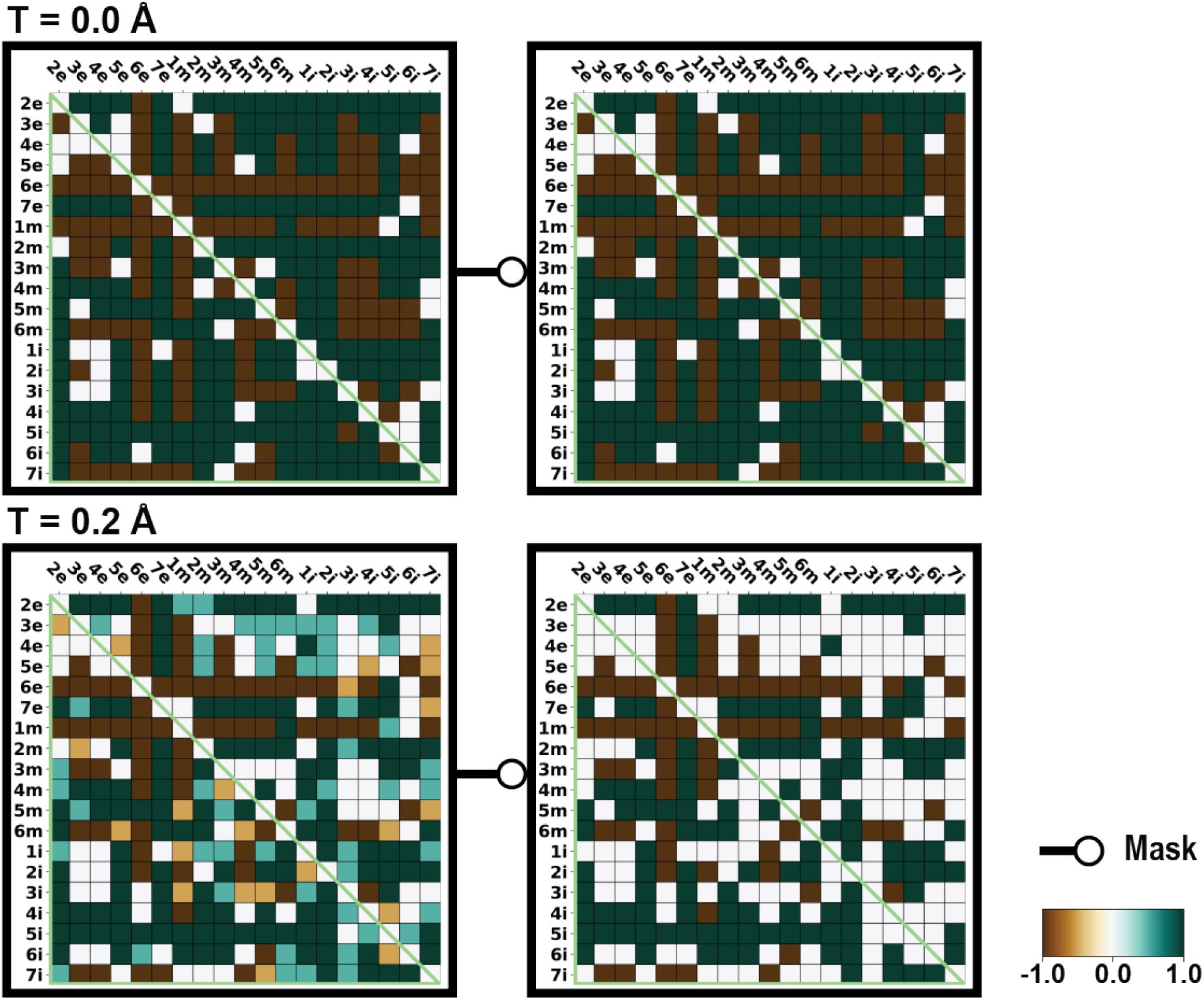
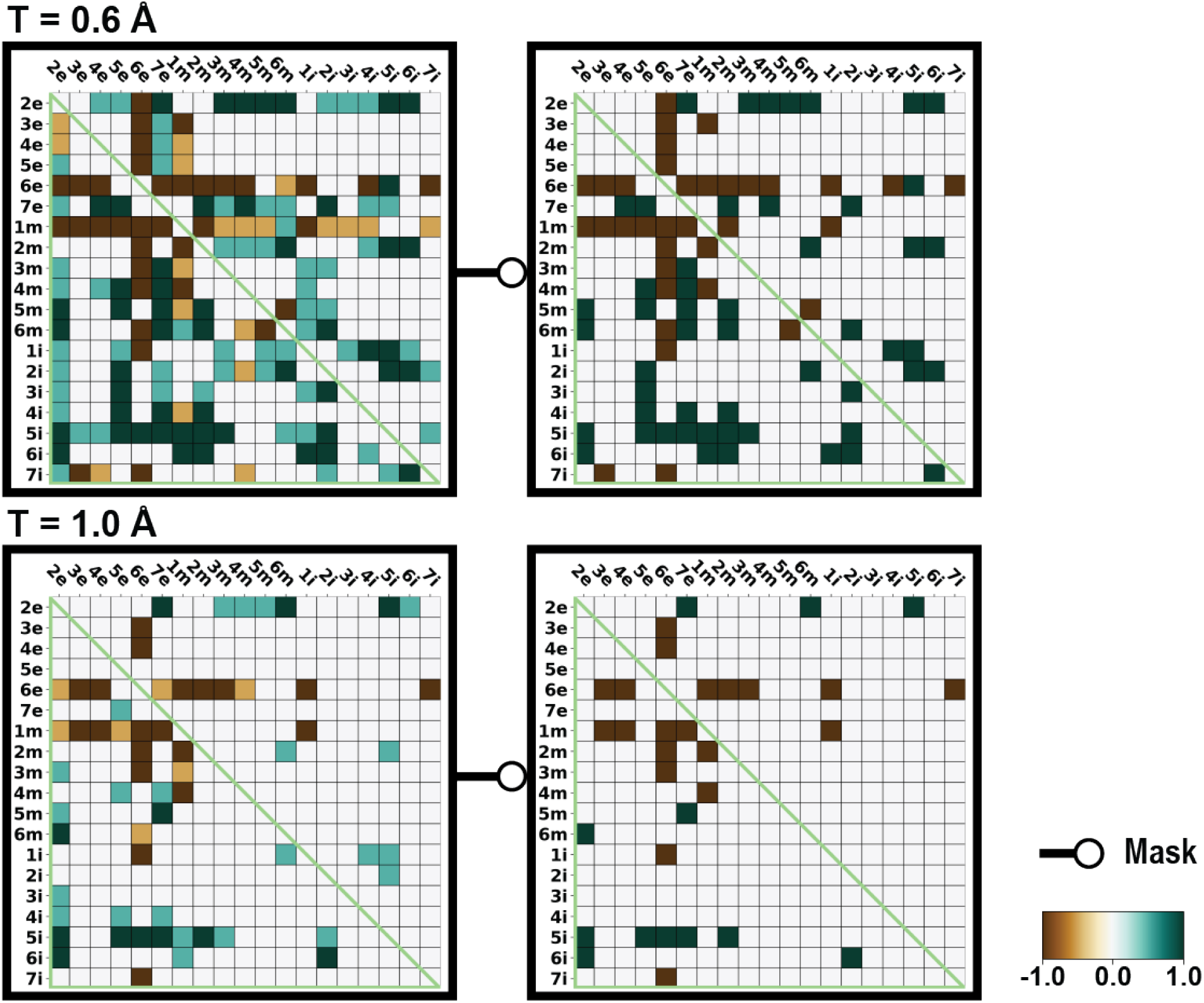
The subsegment distance-difference category averages of the comparison between the active D3R and D2R structures. The heatmap shows the difference of pairwise distance among the subsegments between the D3R and D2R using their active structures (D3R – D2R) and categorized based on various DDthreshold (T). Green represents the D3R conformation have a larger distance than D2R conformation, and brown represents the D2R conformation have a larger distance than D3R conformation. If the category average of the difference is smaller or equal to 0.9, it is colored as white in the masked heatmaps.

**Figure S19.**
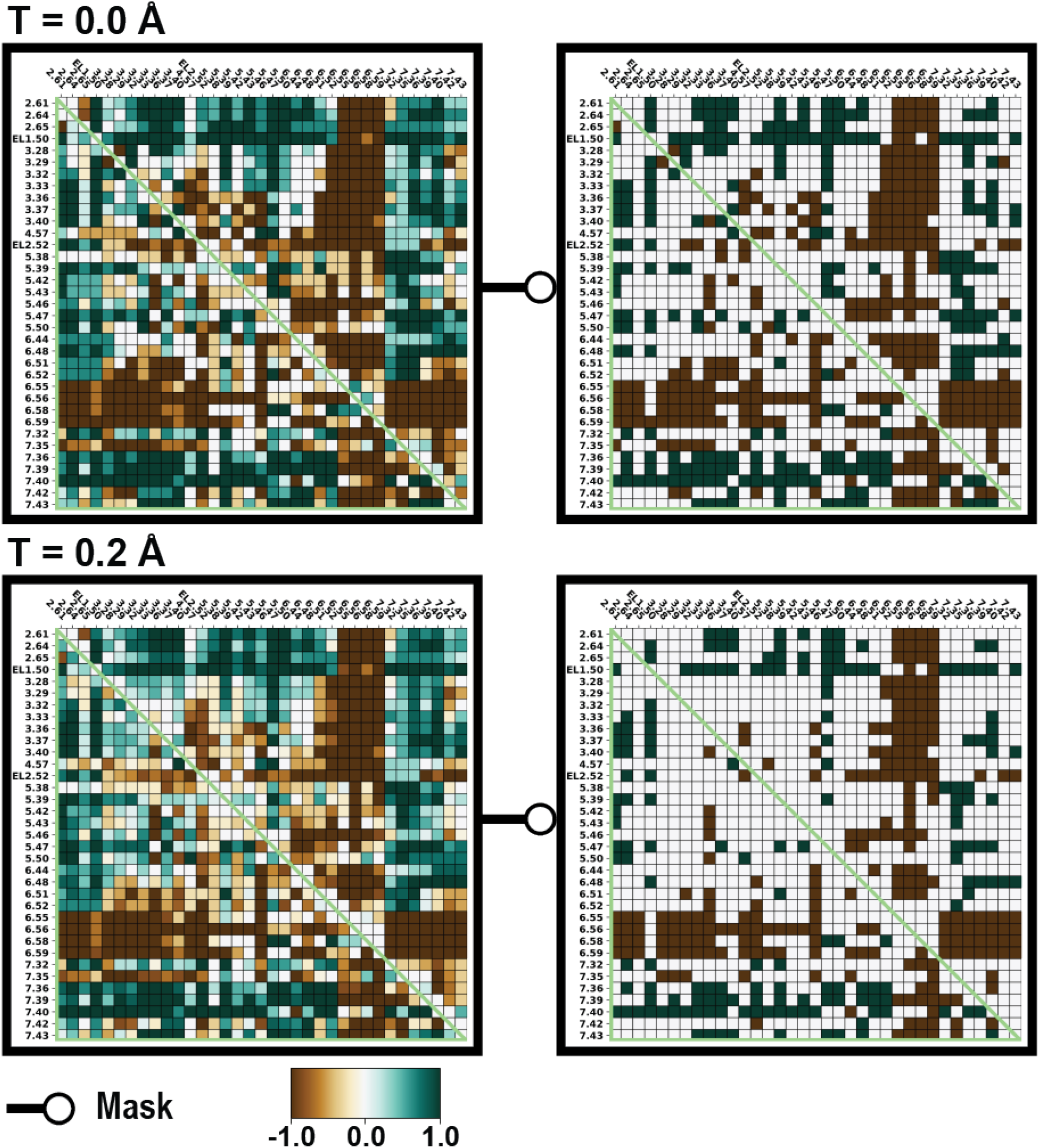
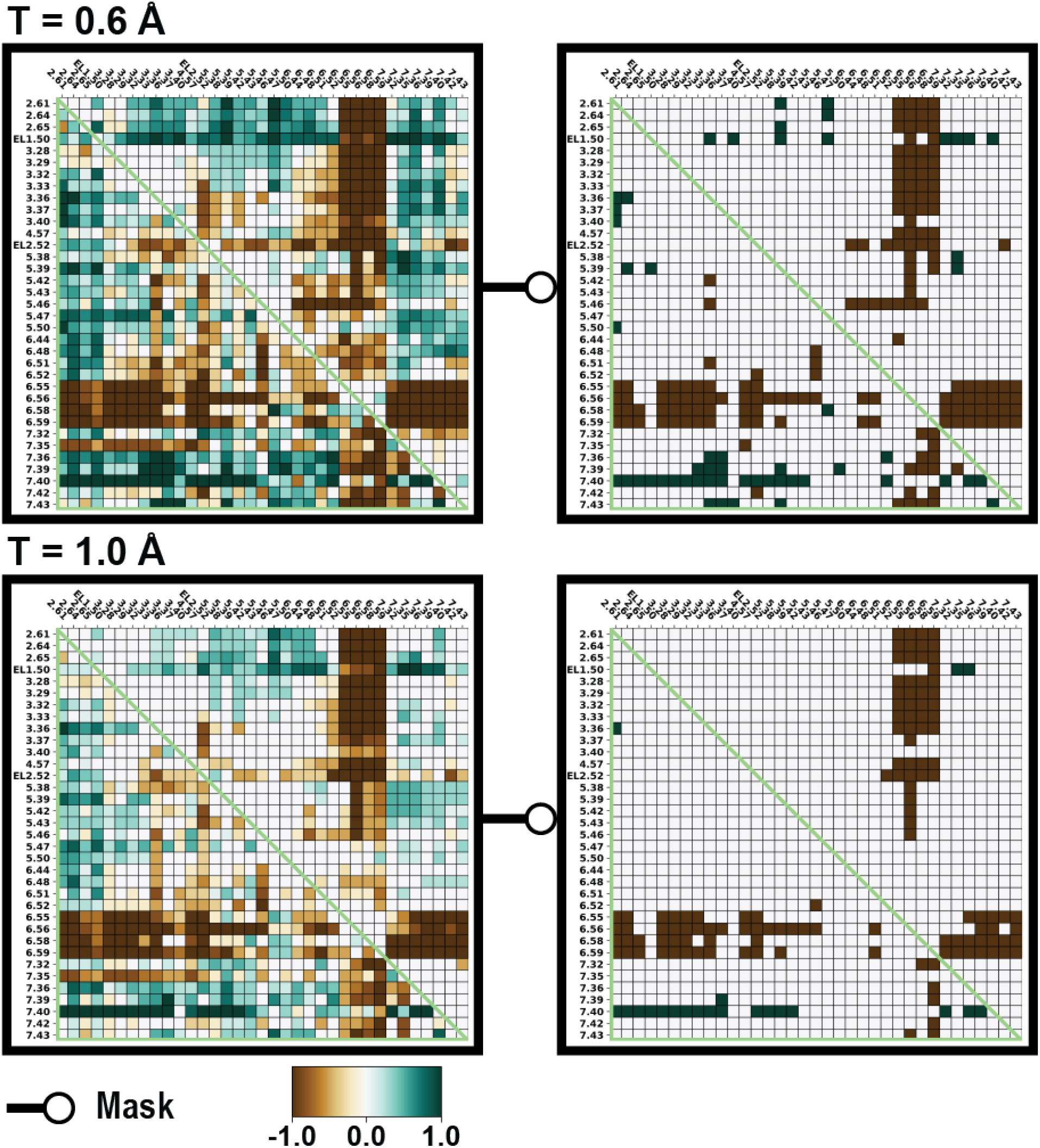
The binding site distance-difference category averages of both the comparisons between the active and between the inactive D3R and D2R structures. The heatmap was combining the derived categories from inactive pairs (**Fig. S13**) and active pairs (**Fig. S16**) with various DDthreshold (T). This heatmap revealed common patterns in distance changes (D3R – D2R) for both inactive and active states of these two receptors. The color green signifies a more distance change in D3R, while brown indicates a greater change in D2R. In instances where the category average of the difference is 0.9 or less, the corresponding region in the masked heatmaps is represented in white.

**Figure S20.**
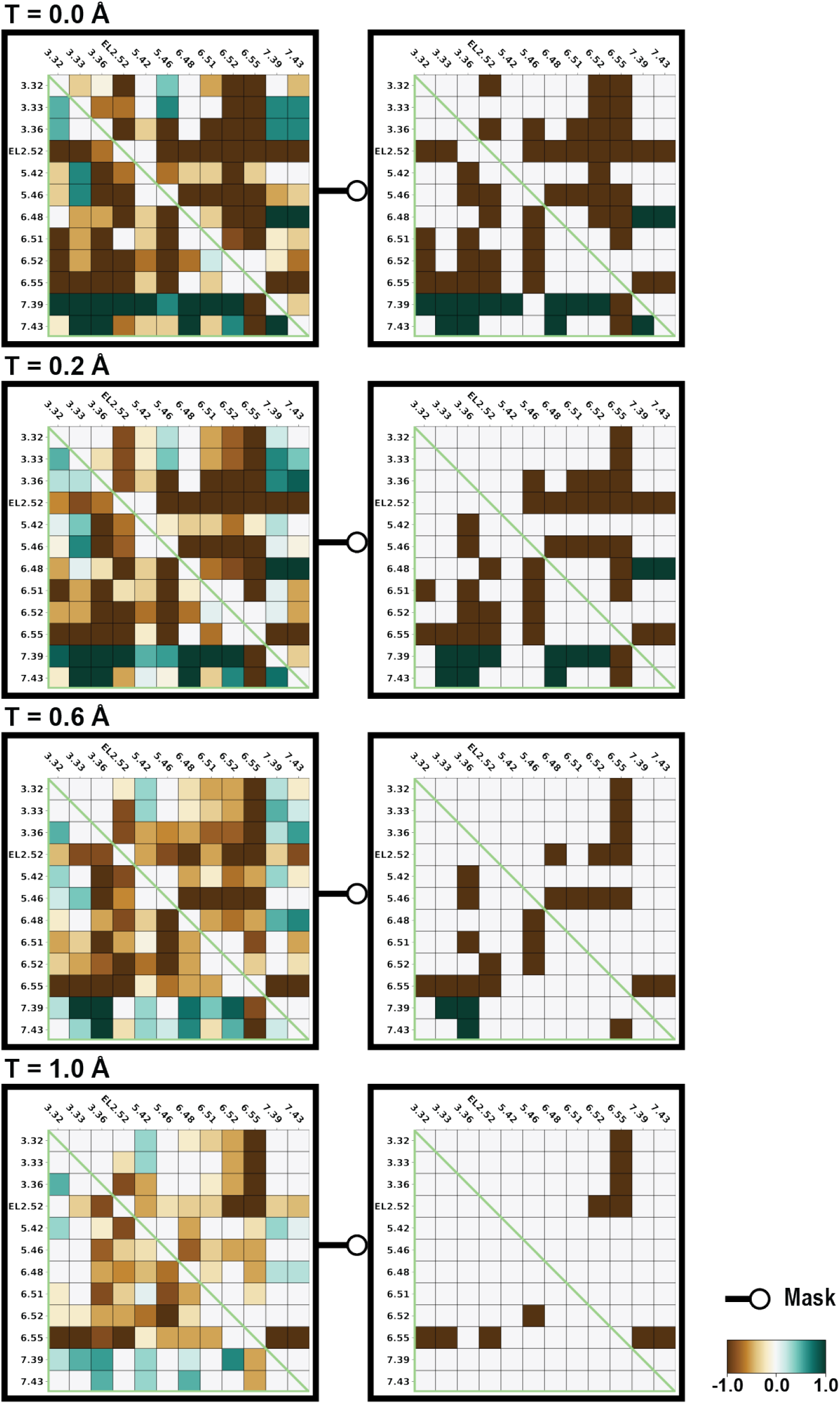
The OBS residue distance-difference category averages of both the comparisons between the active and between the inactive D3R and D2R structures. The heatmap was combining the derived categories from inactive pairs (**Fig. S14**) and active pairs (**Fig. S17**) with various DDthreshold (T). This heatmap revealed common patterns in distance changes (D3R – D2R) for both inactive and active states of these two receptors. The color green signifies a more distance change in D3R, while brown indicates a greater change in D2R. In instances where the category average of the difference is 0.9 or less, the corresponding region in the masked heatmaps is represented in white.

**Figure S21.**
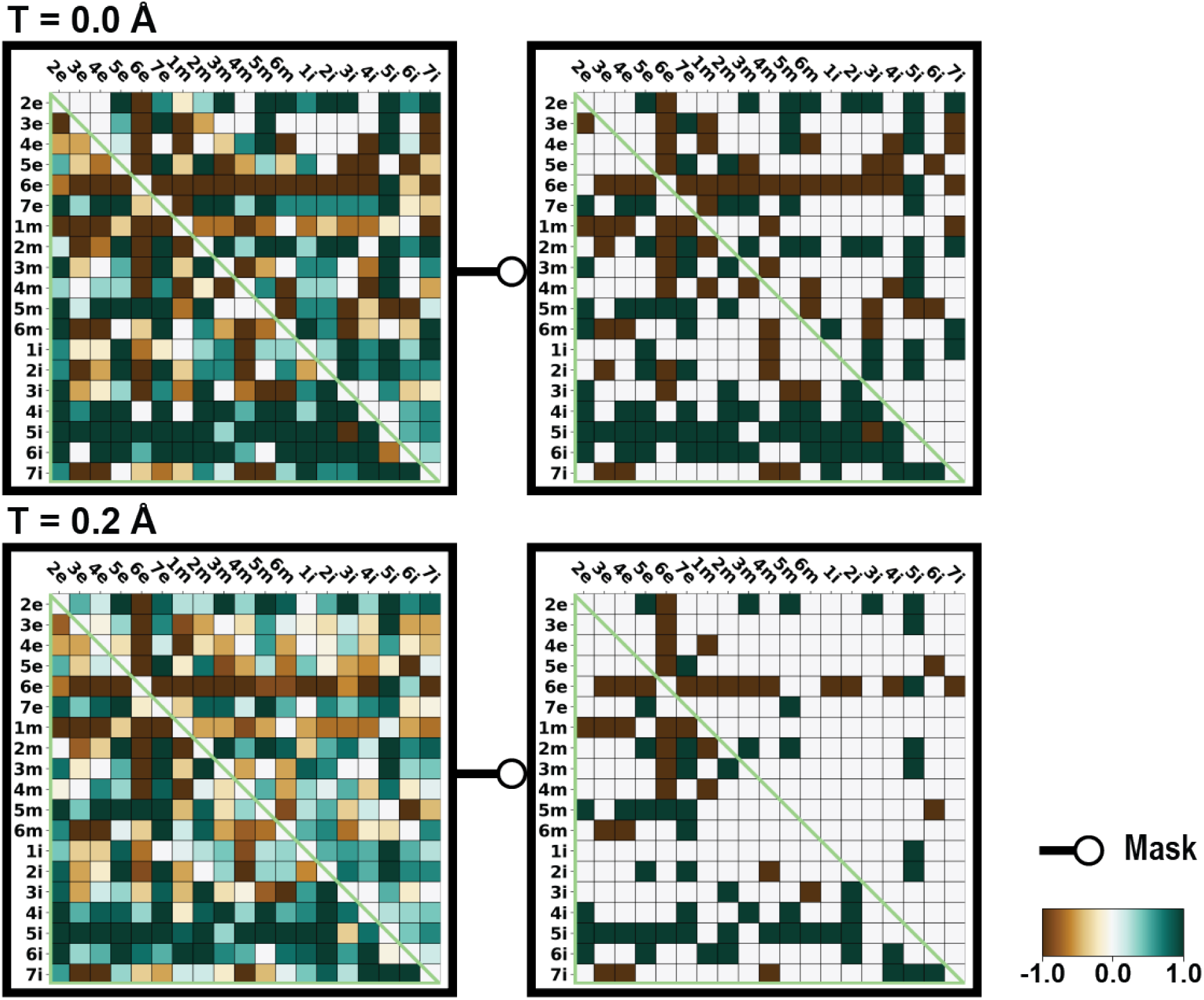
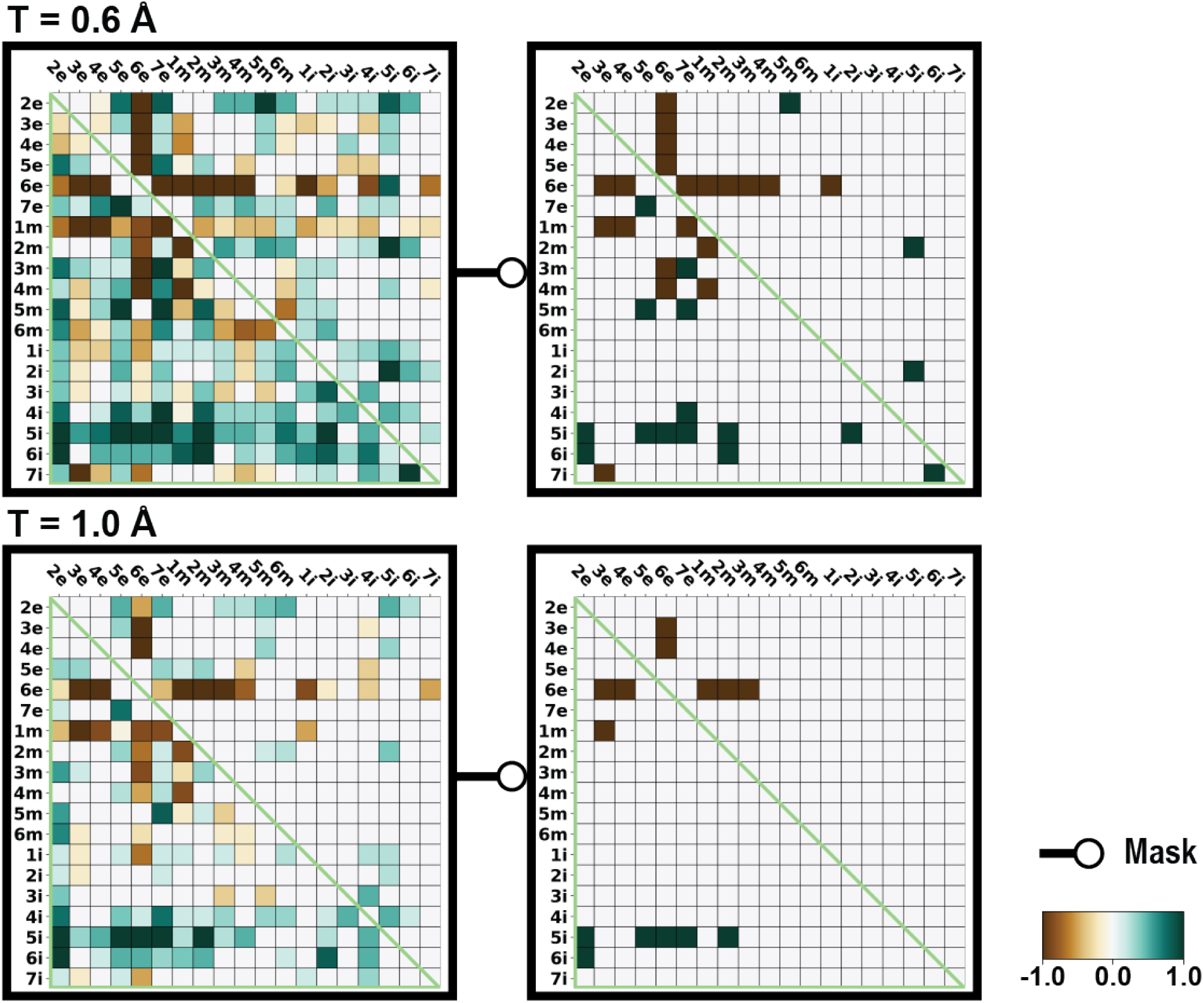
The subsegment distance-difference category averages of both the comparisons between the active and between the inactive D3R and D2R structures. The heatmap was combining the derived categories from inactive pairs (**Fig. S15**) and active pairs (**Fig. S18**) with various DDthreshold (T). This heatmap revealed common patterns in distance changes (D3R – D2R) for both inactive and active states of these two receptors. The color green signifies a more distance change in D3R, while brown indicates a greater change in D2R. In instances where the category average of the difference is 0.9 or less, the corresponding region in the masked heatmaps is represented in white.

**Figure S22.**
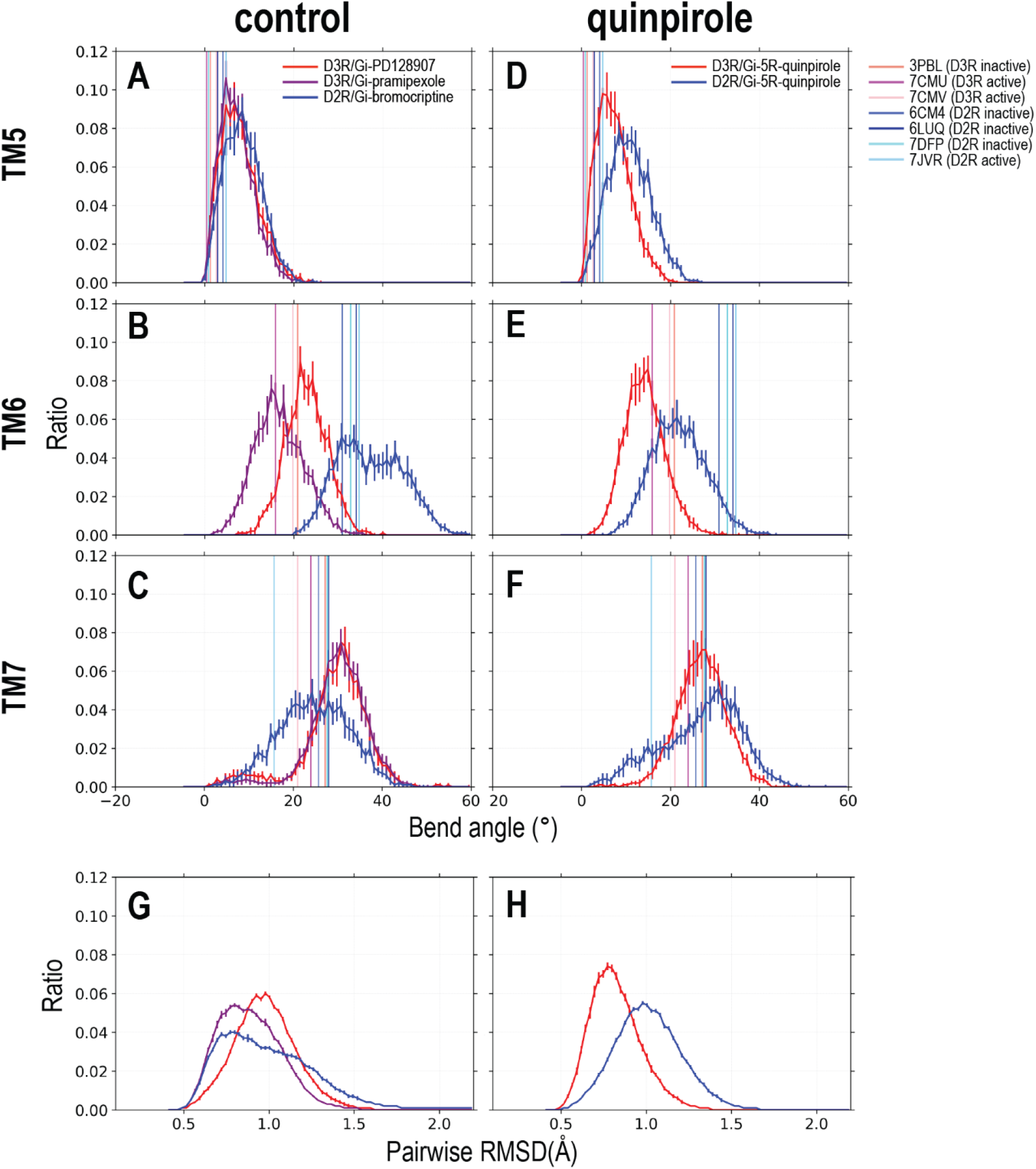
Proline kinks and the ligand binding residues dynamics in the MD simulations. This proline kink (Prokink) measurement was conducted for TM5, TM6, and TM7 using the indicated MD trajectories. Note that the control MD simulations refers to those of D3R/Gi- PD128907, D3R/Gi-pramipexole, and D2R/Gi-bromocriptine. Panel A, B, and C shows the Prokink of TM5, TM6 and TM7 of control simulations, respectively. In panel D, E, and F, the Prokink of TM5, TM6 and TM7 of quinpirole simulations were shown. We assessed the Prokink bend angle with Simulaid. In the calculations, seven residues preceding and succeeding proline were employed for TM5 and TM6, while for TM7, the calculations utilized five residues before and after the proline. The pairwise RMSD of backbone of ligand binding residues were calculated for the same bootstrapped ensemble for control simulations (G) and quinpirole simulations (H). For D3R/Gi-PD128907, D3R/Gi-pramipexole, and D2R/Gi-bromocriptine, the peak positions are 0.97 Å, 0.80 Å, and 0.76 Å, respectively. The peak positions for D2R/Gi- bromocriptine and D3R/Gi-5R-quinpirole are 0.97 Å and 0.77 Å, respectively.

**Figure S23.**
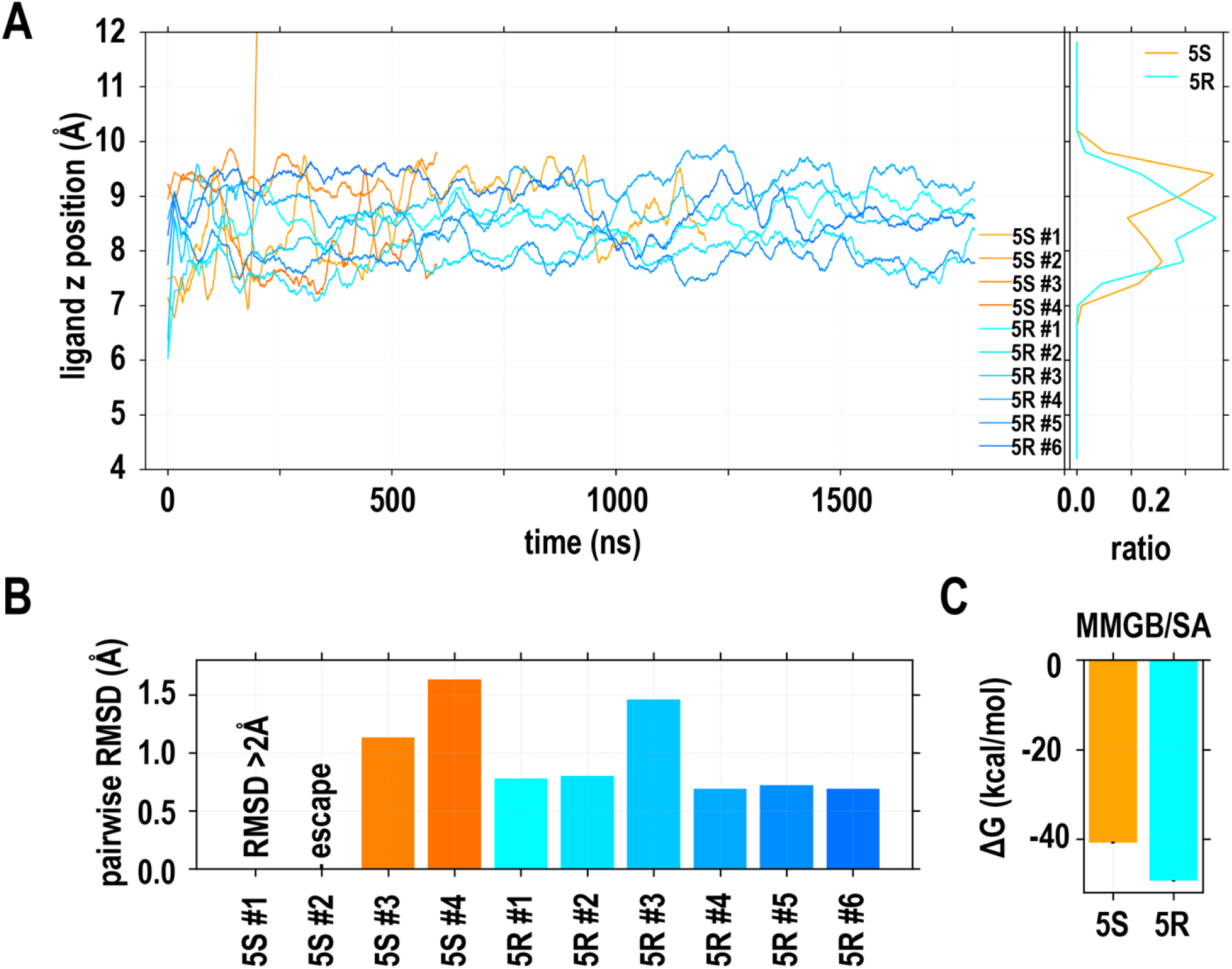
Quinpirole prefers to be in the 5R-isomer in the ligand binding site of D2R. In panel A, the evolutions of the z positions (i.e., along the axis perpendicular to the membrane) of the ligands in both 5S-isomer and 5R-isomer are shown, while the combined distributions of the z positions for these two isomers are shown on the right. When the binding pose of the 5S- isomer was relatively stable, it still exhibited higher z position values compared to the 5R- isomer. The averaged pairwise ligand RMSDs for each MD trajectory are presented in panel B, indicating that the 5R-isomer was more stable than the 5S-isomer. In panel C, the results of the MMGB/SA calculations using the stable trajectories of the 5S- and 5R-isomers are shown, demonstrating the 5R-isomer was bound tighter than the 5S-isomer.

**Figure S24.**
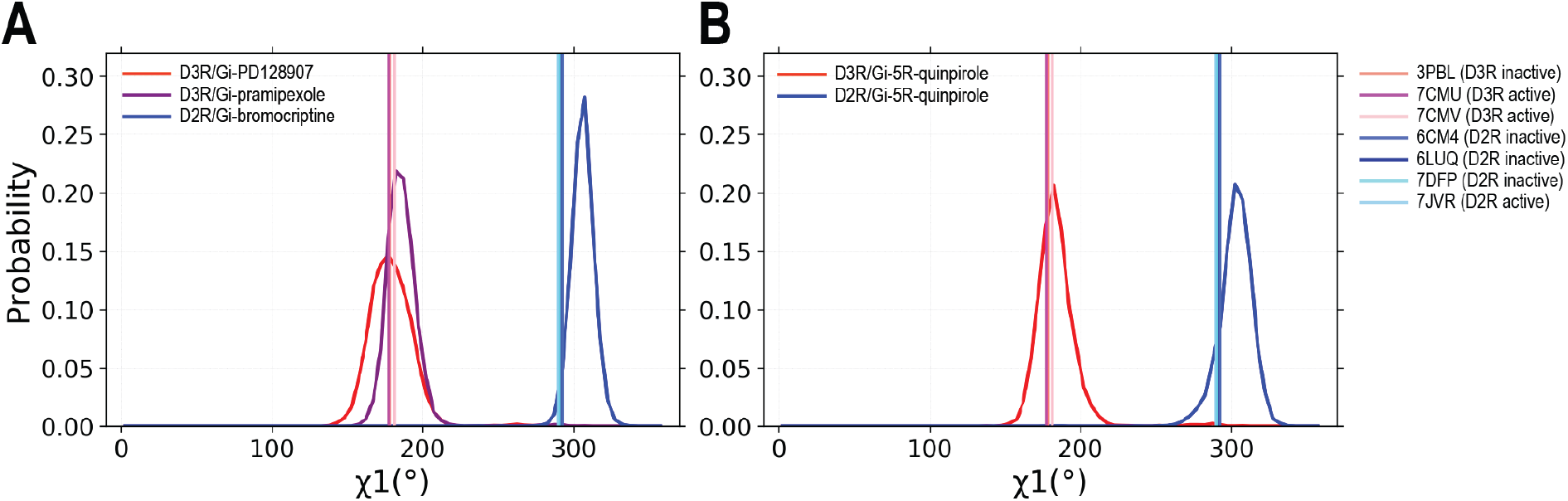
Trp^7.40^ χ1 angle distribution. The χ1 angle distribution was analyzed for two sets of the indicated simulations. For each set of simulations, the χ1 angle distribution was calculated for residue Trp^7.40^. In the plots, the vertical lines indicate the experimentally determined D2R and D3R structures, which serve as references for comparison of the distributions obtained from the simulations.

**Figure S25.**
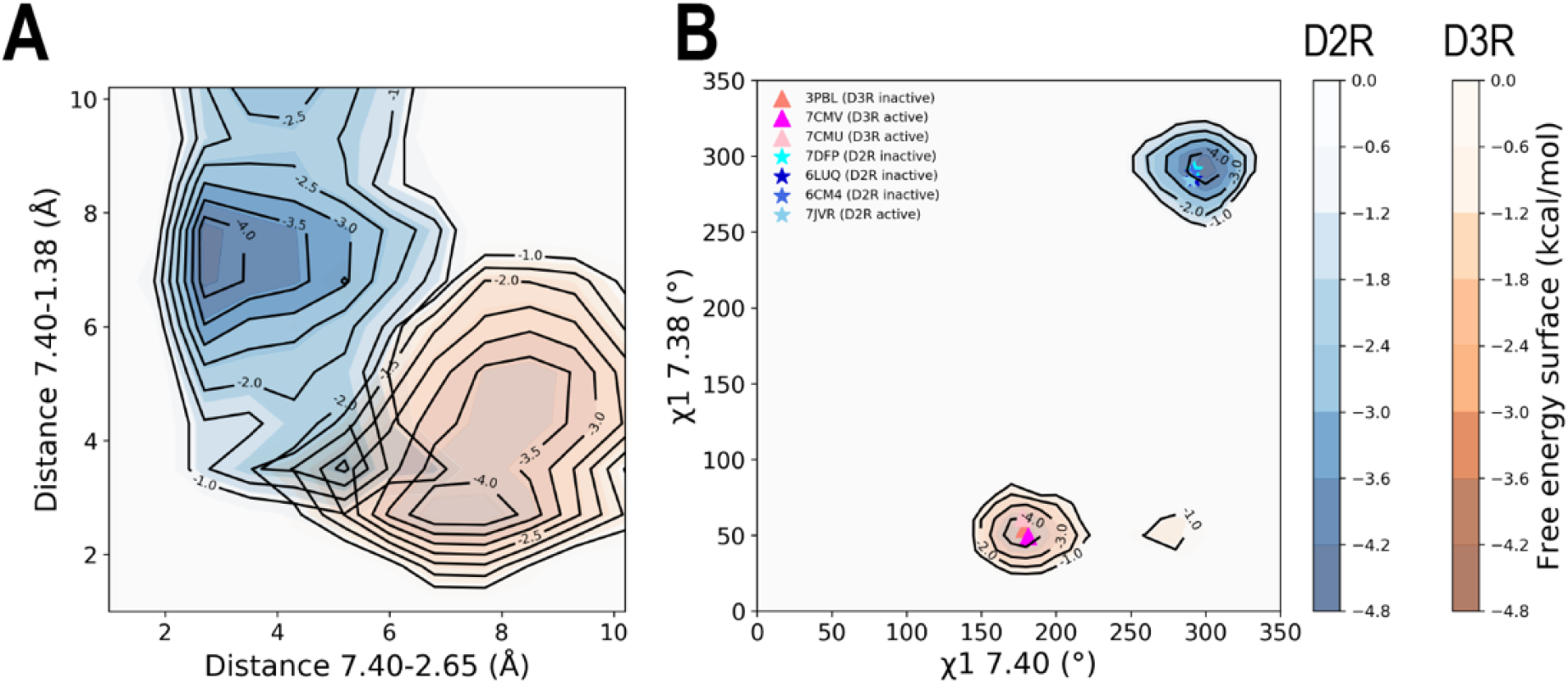
Different conformations of Trp^7.40^ in the 5R-quinpirole bound D3R and D2R. To investigate interactions between Trp^7.40^ and Ser^1.38^ in D3R or Leu^1.38^ in D2R, the closest heavy atom contacting distance between residues 7.40 and 1.38 was calculated. The results show that in the D3R, there is a tendency to form a water-mediated hydrogen bond between these two residues, whereas in the D2R, such an interaction is not observed. (A) The reaction coordinate is defined by the distance of Trp^7.40^– Glu^2.65^ and Trp^7.40^– Ser^1.38^ for D3R, and Trp^7.40^– Leu^1.38^ for the D2R. (B) The reaction coordinate is determined using the χ1 angles of Trp^7.40^ and Thr^7.38^ for the D3R, and Trp^7.40^ and Phe^7.38^ for the D2R. The various symbols indicate the corresponding values in the experimentally determined structures.

**Figure S26.**
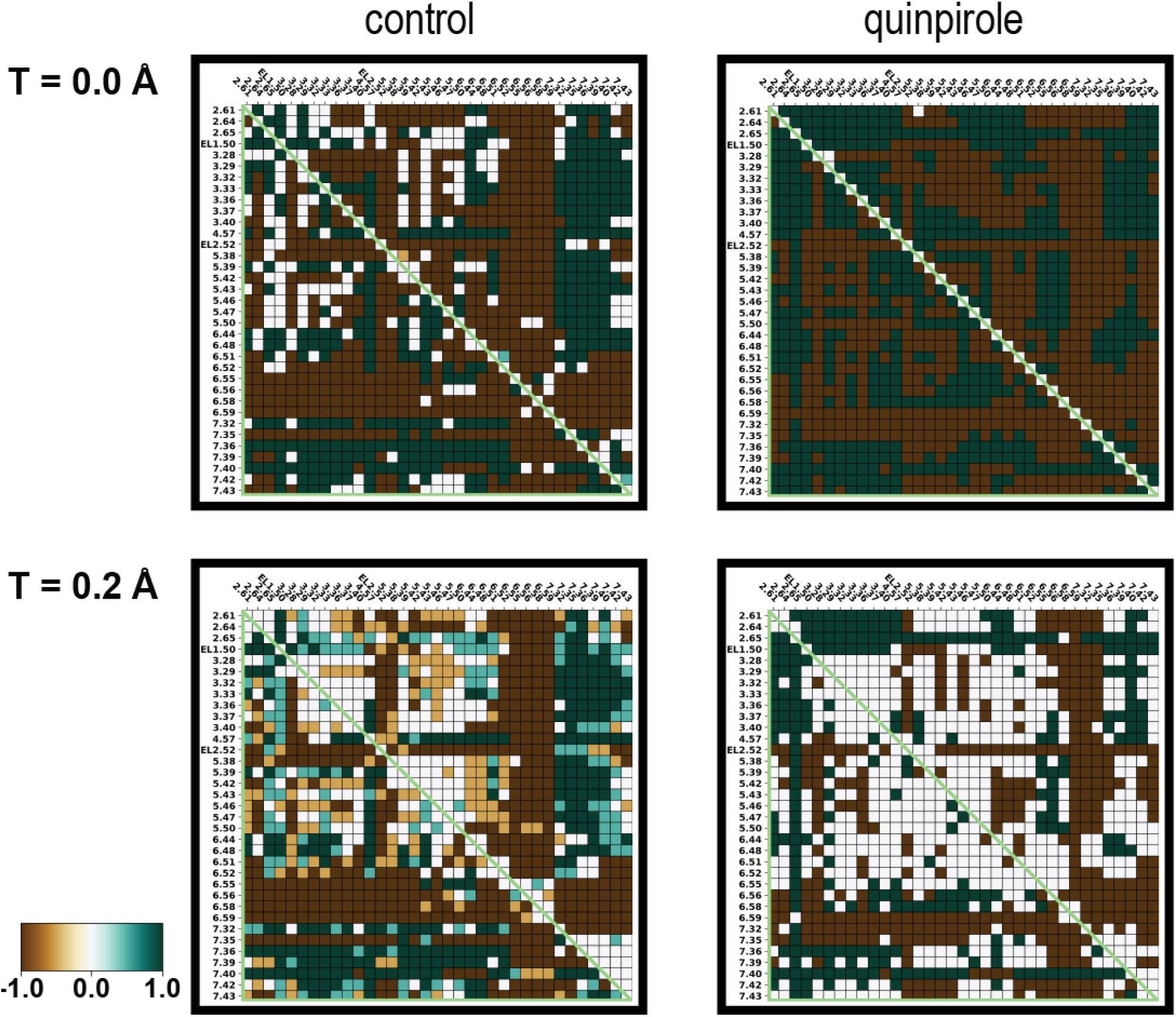
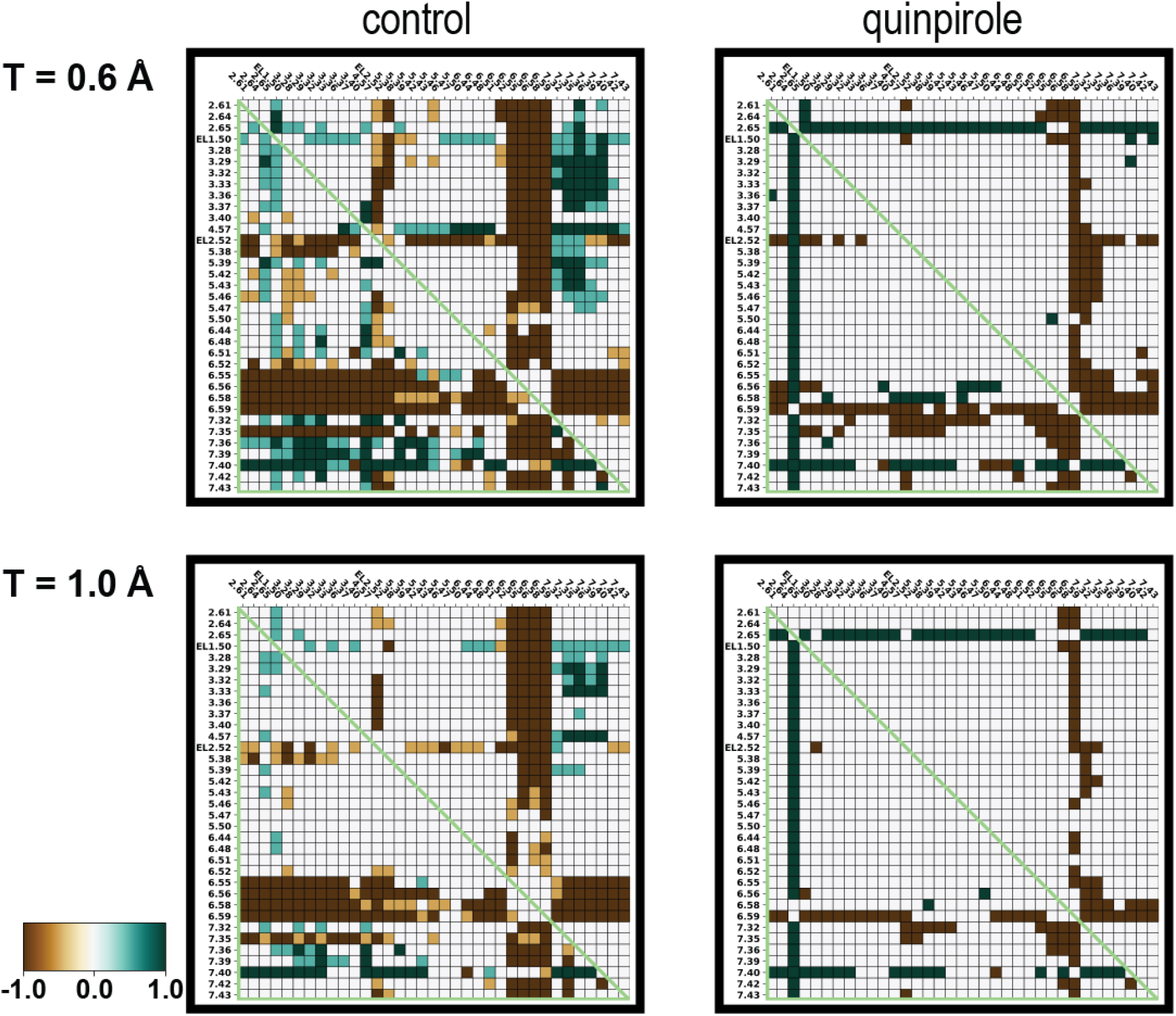
The binding site distance-difference category averages of the comparisons between the D3R and D2R MD simulations. The heatmap shows the difference of pairwise distance among the binding site residues between the D3R and D2R using their active structures (D3R – D2R) and categorized based on various DDthreshold (T). Green represents the D3R conformation have a larger distance than D2R conformation, and brown represents the D2R conformation have a larger distance than D3R conformation. Note that the MD simulations from D3R/Gi-PD128907, D3R/Gi-pramipexole, and D2R/Gi-bromocriptine are referred as “control”, and simulations from D3R/Gi-5R-quinpirole, and D2R/Gi-5R-quinpirole are referred as “quinpirole”.

**Figure S27.**
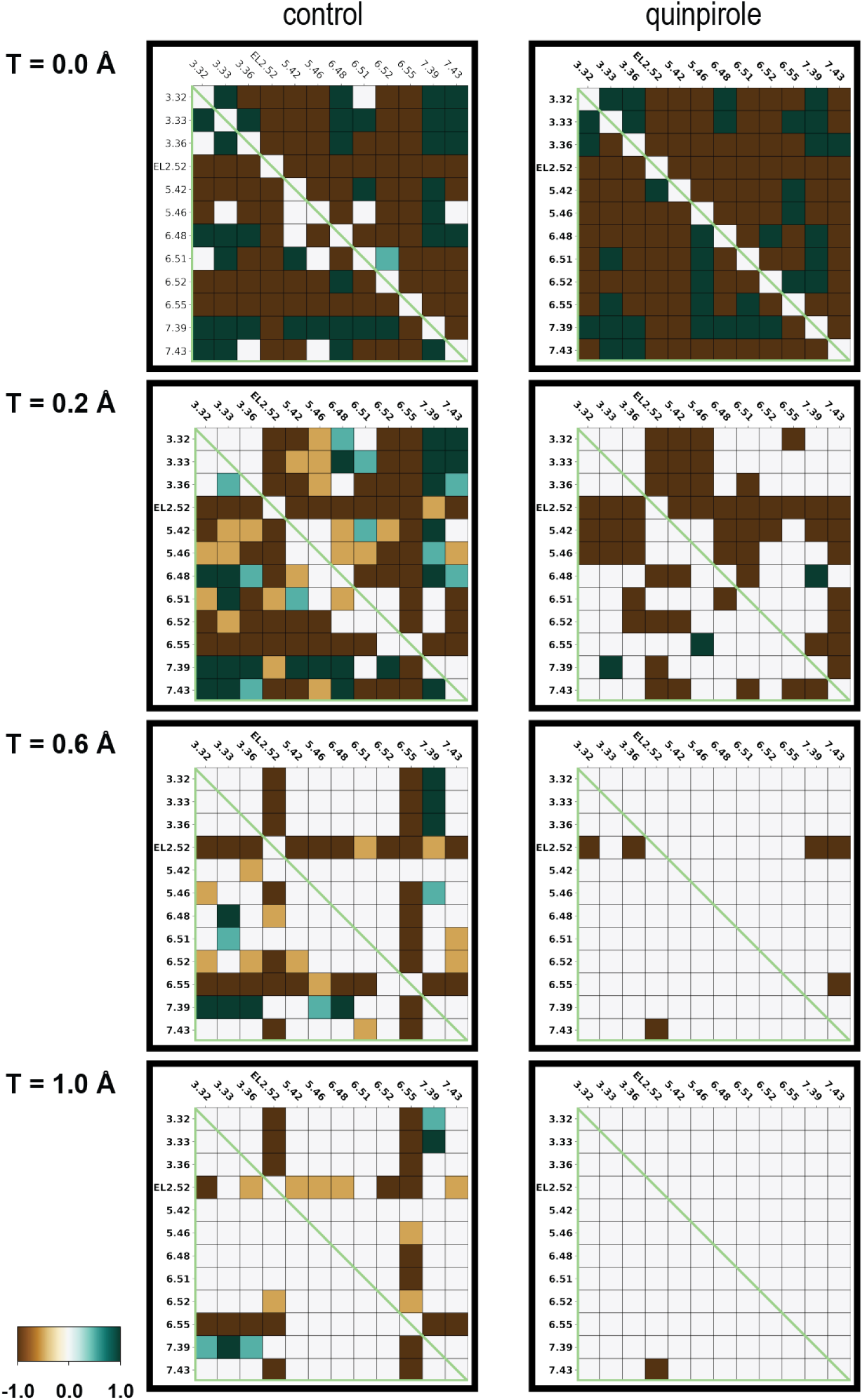
The OBS residue distance-difference category averages of the comparisons between the D3R and D2R MD simulations. The heatmap shows the difference of pairwise distance among the binding site residues between the D3R and D2R using their active structures (D3R – D2R) and categorized based on various DDthreshold (T). Green represents the D3R conformation have a larger distance than D2R conformation, and brown represents the D2R conformation have a larger distance than D3R conformation. Note that the MD simulations from D3R/Gi-PD128907, D3R/Gi-pramipexole, and D2R/Gi-bromocriptine are referred as “control”, and simulations from D3R/Gi-5R-quinpirole, and D2R/Gi-5R-quinpirole are referred as “quinpirole”.

**Figure S28.**
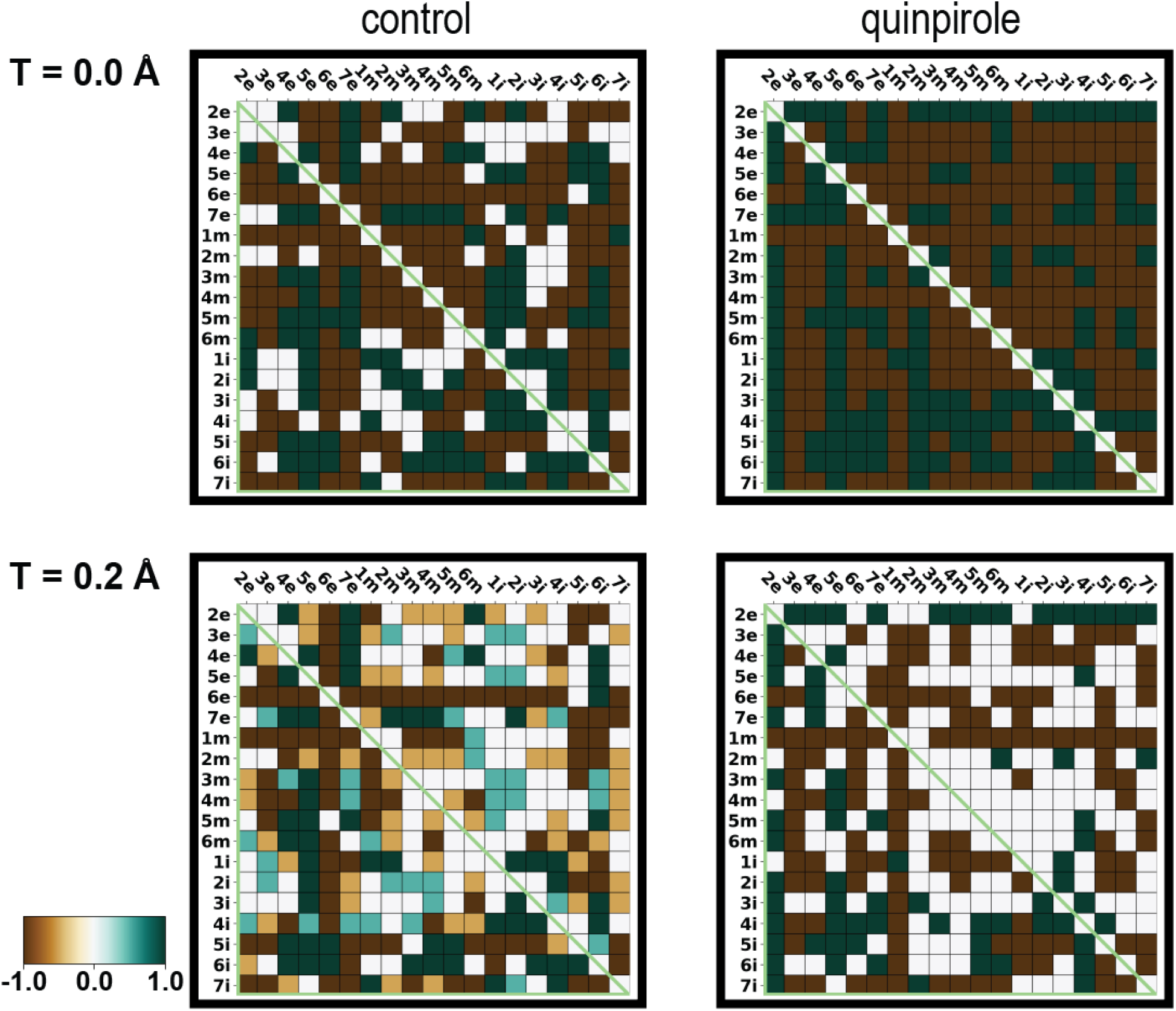
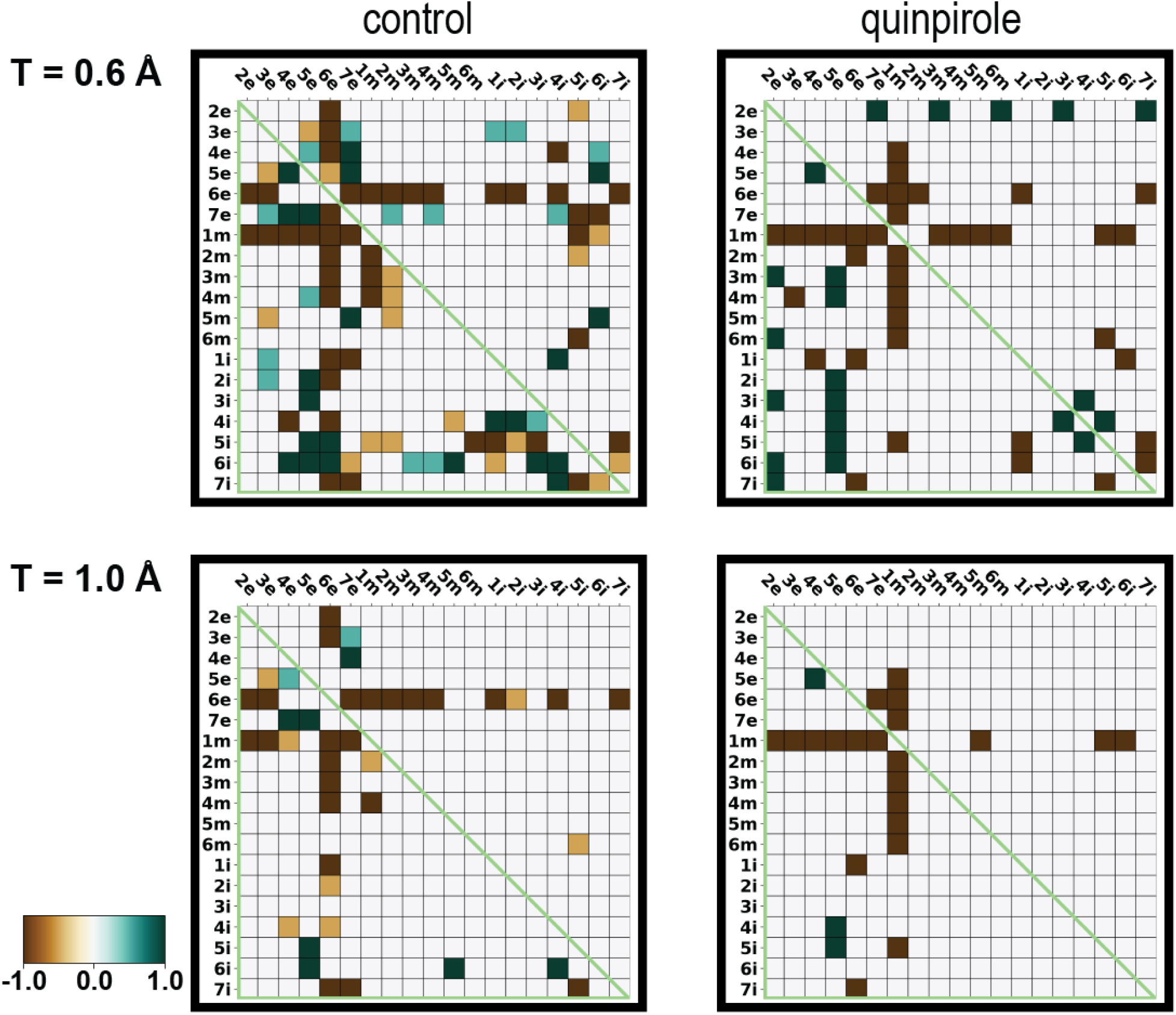
The subsegment distance-difference category averages of the comparisons between the D3R and D2R MD simulations. The heatmap shows the difference of pairwise distance among the binding site residues between the D3R and D2R using their active structures (D3R – D2R) and categorized based on various DDthreshold (T). Green represents the D3R conformation have a larger distance than D2R conformation, and brown represents the D2R conformation have a larger distance than D3R conformation. Note that the MD simulations from D3R/Gi-PD128907, D3R/Gi-pramipexole, and D2R/Gi-bromocriptine are referred as “control”, and simulations from D3R/Gi-5R-quinpirole, and D2R/Gi-5R-quinpirole are referred as “quinpirole”.

**Table S1.** The experimentally determined D2R and D3R structures used in this study.

| condition | PDB ID | publication year | resolution (Å) | bound ligand | reference |
| --- | --- | --- | --- | --- | --- |
| **D2R inactive** | 7DFP | 2020 | 3.1 | spiperone | 1 |
|  | 6LUQ | 2020 | 3.1 | haloperidol | 2 |
|  | 6CM4 | 2018 | 2.9 | risperidone | 3 |
| **D2R active** | 7JVR | 2021 | 2.8 | bromocriptine | 4 |
| **D3R inactive** | 3PBL | 2010 | 2.9 | eticlopride | 5 |
| **D3R active** | 7CMU | 2021 | 3.0 | pramipexole | 6 |
|  | 7CMV | 2021 | 2.7 | PD128907 | 6 |

Note that we chose to use the structure 7JVR because it has a higher resolution than the other D2R/G<sub>i</sub>-bromocriptine structure (PDB 6VMS)<sup>7</sup>.

**Table S2.**
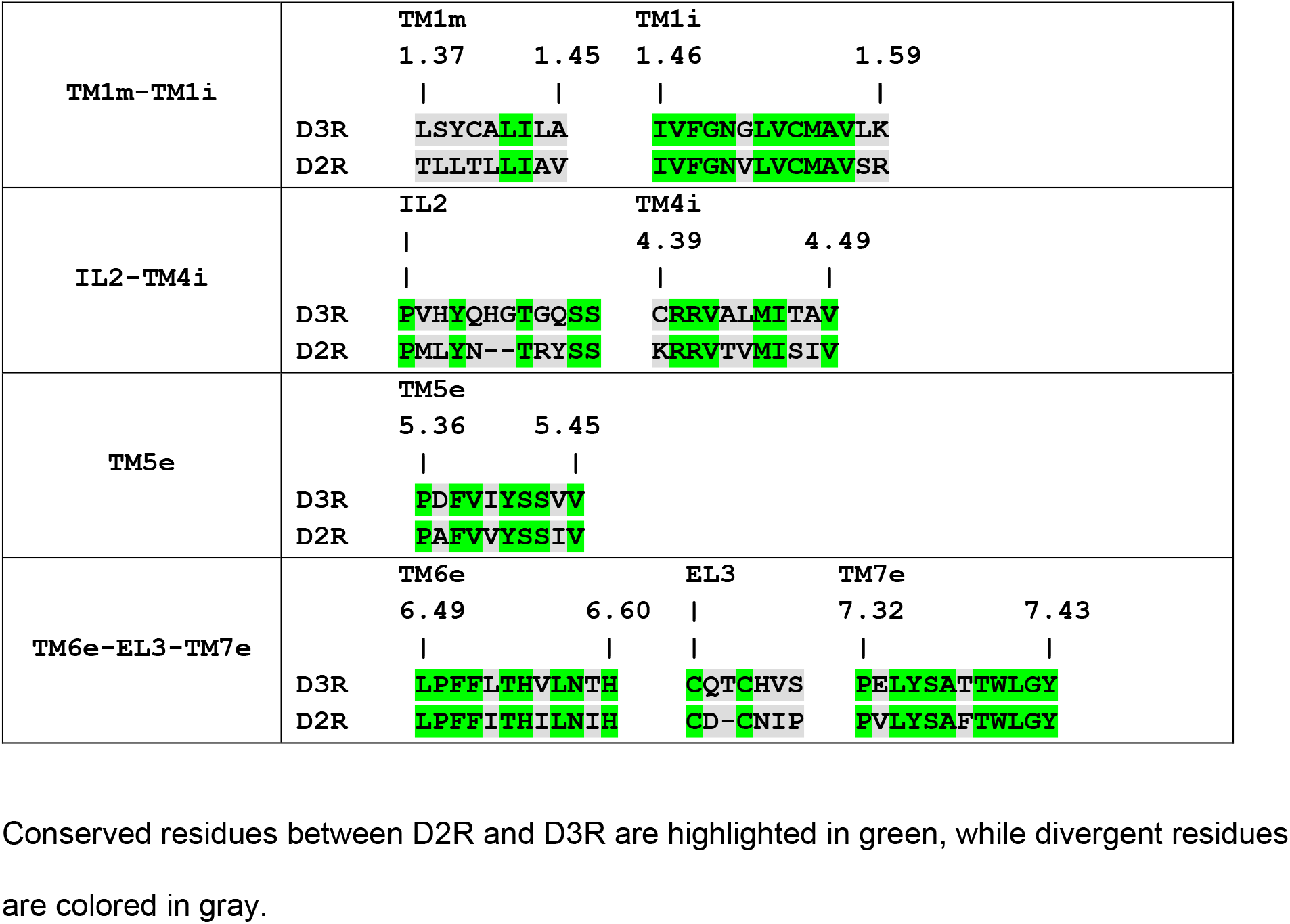
Sequence alignment of TM1m and TM1i.

**Table S3.** Prokink bend angles of TM5, TM6 and TM7.

| source | state | condition | TM5 | TM6 | TM7 |
| --- | --- | --- | --- | --- | --- |
| structure | D3R inactive | 3PBL | 1.3 | 20.9 | 27.2 |
|  | D2R inactive | 6CM4 | 4.1 | 30.9 | 25.7 |
|  |  | 6LUQ | 2.9 | 34.1 | 28.0 |
|  |  | 7DFP | 0.9 | 32.8 | 27.7 |
|  | inactive difference | 3PBL – 6CM4 | -2.8 | -10.0 | 1.5 |
|  |  | 3PBL – 6LUQ | -1.6 | -13.2 | -0.8 |
|  |  | 3PBL – 7DFP | 0.4 | -11.9 | -0.5 |
|  | **average of inactive difference** |  | **-1.3** | **-11.7** | **0.1** |
|  | D3R active | 7CMV (PD128907) | 2.6 | 19.8 | 21.0 |
|  |  | 7CMU (pramipexole) | 0.4 | 15.9 | 24.0 |
|  | D2R active | 7JVR (bromocriptine) | 4.8 | 34.7 | 15.7 |
|  | active difference | 7CMV (PD128907) – 7JVR (bromocriptine) | -2.2 | -14.9 | 5.3 |
|  |  | 7CMU (pramipexole) – 7JVR (bromocriptine) | -4.4 | -18.8 | 8.3 |
|  | **average of active difference** |  | **-3.3** | **-16.9** | **6.8** |
| MD | D3R active | D3R/G<sub>i</sub>-PD128907 | 6.6 | 22.4 | 31.4 |
|  |  | D3R/G<sub>i</sub>-pramipexole | 5.9 | 15.6 | 30.2 |
|  | D2R active | D2R/G<sub>i</sub>-bromocriptine | 7.8 | 32.6 | 23.6 |
|  | active difference | D3R/G<sub>i</sub>-PD128907 - D2R/G<sub>i</sub>-bromocriptine | -1.2 | -10.3 | 7.8 |
|  |  | D3R/G<sub>i</sub>-pramipexole - D2R/G<sub>i</sub>-bromocriptine | -2.0 | -17.0 | 6.6 |
|  | **average of active difference** |  | **-1.6** | **-13.7** | **7.2** |
|  | D3R active | D3R/G<sub>i</sub>-5R-quinpirole | 5.7 | 14.2 | 26.9 |
|  | D2R active | D2R/G<sub>i</sub>-5R-quinpirole | 10.0 | 20.0 | 30.7 |
|  | **D3R/G<sub>i</sub>-5R-quinpirole – D2R/G<sub>i</sub>-5R-quinpirole** |  | **-4.3** | **-5.7** | **-3.8** |

The Prokink bend angle of individual structure or MD frame was first calculated with Simulaid.<sup>8</sup>.
The inactive average is the average of 3PBL-6CM4, 3PBL-6LUQ, and 3PBL-7DFP. The active average is the average of 7CMU-7JVR, and 7CMV-7JVR. For the MD simulations, from the bend angle distributions of bootstrap ensembles for each condition, we identified the peak values of these distributions, and calculated the differences between the indicated conditions. Finally, we computed the control MD average using D3R/G<sub>i</sub>-PD128907 - D2R/G<sub>i</sub>-bromocriptine and D3R/G<sub>i</sub>-pramipexole - D2R/G<sub>i</sub>-bromocriptine.

**Table S4.** MD simulation summary.

| **protein complex** | **bound ligand** | **runs** | **length**<br>**(μs)** |
| --- | --- | --- | --- |
| D2R/Gi | 5S-quinpirole | 5 | 3.8 |
| D2R/Gi | 5R-quinpirole | 6 | 10.8 |
| D2R/Gi | bromocriptine | 6 | 10.8 |
| D3R/Gi | 5S-quinpirole | 8 | 11.3 |
| D3R/Gi | 5R-quinpirole | 6 | 10.8 |
| D3R/Gi | pramipexole | 6 | 10.8 |
| D3R/Gi | PD128907 | 6 | 10.8 |

**Table S5.**
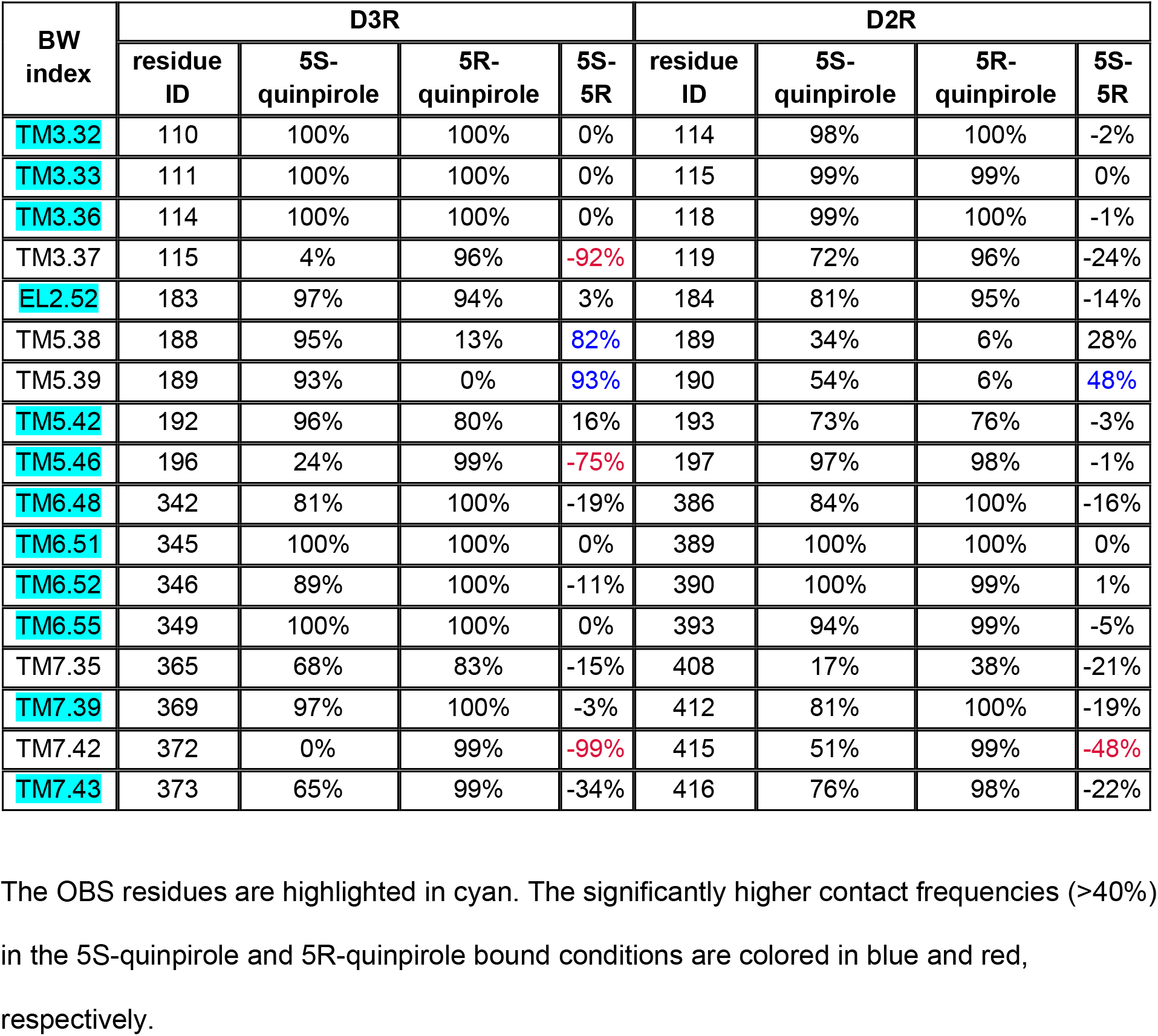
Ligand contact frequency observed with the 5S-quinpirole and 5R-quinpirole bound D3R and D2R MD simulations.

